# Structures of MeCP2 bound to nucleosomes reveal distinct mechanisms of Rett syndrome mutations

**DOI:** 10.64898/2026.09.08.750195

**Authors:** Liqi Yao, Carolina Valderrama-Hincapié, Jiarui Song, Iona L. Kelly, Edward B. Chuong, Vignesh Kasinath

**Affiliations:** Department of Biochemistry, University of Colorado Boulder, Boulder, CO, 80303; BioFrontiers Institute, University of Colorado Boulder, CO 80303; Department of Molecular, Cellular, and Developmental Biology, University of Colorado Boulder, CO 80303; Howard Hughes Medical Institute, University of Colorado Boulder, CO 80303

## Abstract

Methyl-CpG binding protein 2 (MeCP2) is a chromatin-associated regulator essential for neuronal gene regulation, and pathogenic mutations in MeCP2 cause Rett syndrome (RTT). However, the mechanisms governing MeCP2 engagement on chromatin and the effects of RTT mutations on nucleosome interactions remain poorly understood. We determined cryo-EM structures of MeCP2 bound to mono-nucleosomes with or without linker DNA methylation. In the absence of linker DNA methylation, the methyl-CpG binding domain (MBD) engages nucleosomal DNA near superhelical location ±7, whereas a methylated linker CpG redirects MBD to the linker methylation site. Quantitative EMSA, MNase footprinting, and fluorescence polarization show that both MBD and AT-hook regions cooperate to stabilize nucleosome binding, and that six common RTT variants (R133C, T158M, R306C, R168X, R255X, R270X) fall into four mechanistic classes. Loss-of-function truncations progressively weaken binding, whereas missense variants show near-WT DNA affinity. Among the variants, R133C shows near wild-type affinity for naked DNA but loses methylation-directed nucleosome engagement. Our analysis of human neuronal transcriptomic and chromatin occupancy datasets showed that RTT variants R133C and R168X lose CpG-island specificity and redistribute in the genome through distinct biochemical routes. Together, this work establishes a framework linking variant-specific defects in nucleosome binding to chromatin-targeting failure and transcriptional dysregulation in RTT.

## Introduction

Methyl-CpG-binding protein 2 (MeCP2) is expressed at near-histone stoichiometry in mature neurons and serves as a global regulator of transcription across a broad range of target loci^1,2^. Mutations in the X-linked *MECP2* gene cause Rett syndrome (RTT), one of the most common monogenic causes of intellectual disability in females, affecting approximately 1 in 10,000 girls^3,4^. Despite decades of study, the molecular mechanisms by which MeCP2 engages chromatin and how pathogenic mutations disrupt this engagement remain poorly understood.

MeCP2 is organized into an N-terminal domain (NTD), a methyl-CpG-binding domain (MBD), an intervening domain (ID) containing AT-hook regions, a transcriptional repression domain (TRD) harboring the NCoR/SMRT-interaction domain (NID), and a C-terminal domain (CTD)^5,6^. Apart from the MBD, MeCP2 is intrinsically disordered, which has precluded high-resolution structural studies with chromatin^7,8^. Early work established that MeCP2 reads methylated CpG through its MBD, recruits histone deacetylase complexes via the TRD, binds preferentially to nucleosomes with linker DNA, and competes with histone H1, and that an arginine-and lysine-rich region (aa 205-257) provides additional linker DNA contacts important for reading methylated cytosines within the nucleosome^9–14^.

Despite these advances, fundamental questions remain unanswered. There are currently no high-resolution structures of MeCP2 on a nucleosome. The basis for MBD positioning, and whether this positioning is driven by DNA sequence, methylation, or histone contacts, is uncharacterized. For the most frequently observed RTT mutations, quantitative nucleosome-binding data have been limited, and it is unclear whether they impair methylation readout, general nucleosome affinity, or co-repressor recruitment, and to what extent these mechanisms are separable.

Here, we report 2.9-8 Å cryo-EM structures of MeCP2 bound to nucleosomes with flanking linker DNA, in the presence and absence of CpG DNA hemi-methylation. We find that the MBD domain of MeCP2 is the primary nucleosome-binding module but is insufficient to engage nucleosomes on its own. The intervening domain containing the AT-hook regions is also required for stable nucleosome association. We defined plasticity in MeCP2 binding to nucleosomes as the presence of DNA methylation in the linker region forces the MBD to localize to the site of methylation without altering overall binding affinity. Additionally, new analyses of published RNA-seq and chromatin occupancy data in human neuronal cells for three RTT variants (R270X, R133C, R168X) reveal that these variants converge on the loss of CpG island specificity, demonstrating that mechanistically distinct biochemical defects produce a shared genome-wide redistribution phenotype.

## Results

### MeCP2 MBD anchors at SHL±7 and relocates to the site of linker methylation

To determine the molecular basis for MeCP2 interaction with nucleosomes, we reconstituted full-length human MeCP2 (isoform e2) with a mononucleosome assembled on the Widom 601 DNA sequence^15^ with 40-bp linker arms on each side (referred to as 40-N-40; N represents the 147 bp 601-Widom core sequence) (Supplementary Fig. 1) Single-particle cryo-EM study of this reconstituted complex using conventional methods yielded nucleosomes without MeCP2 density, likely from denaturation at the air–water interface^16^. To overcome this bottleneck, we used streptavidin–biotin affinity EM grids and biotinylated nucleosomes to obtain a 2.9 Å reconstruction of the MeCP2–unmodified nucleosome complex (Fig. 1a–b, Supplementary Fig. 2,3; Table 1)^17,18^.

**Table 1.**
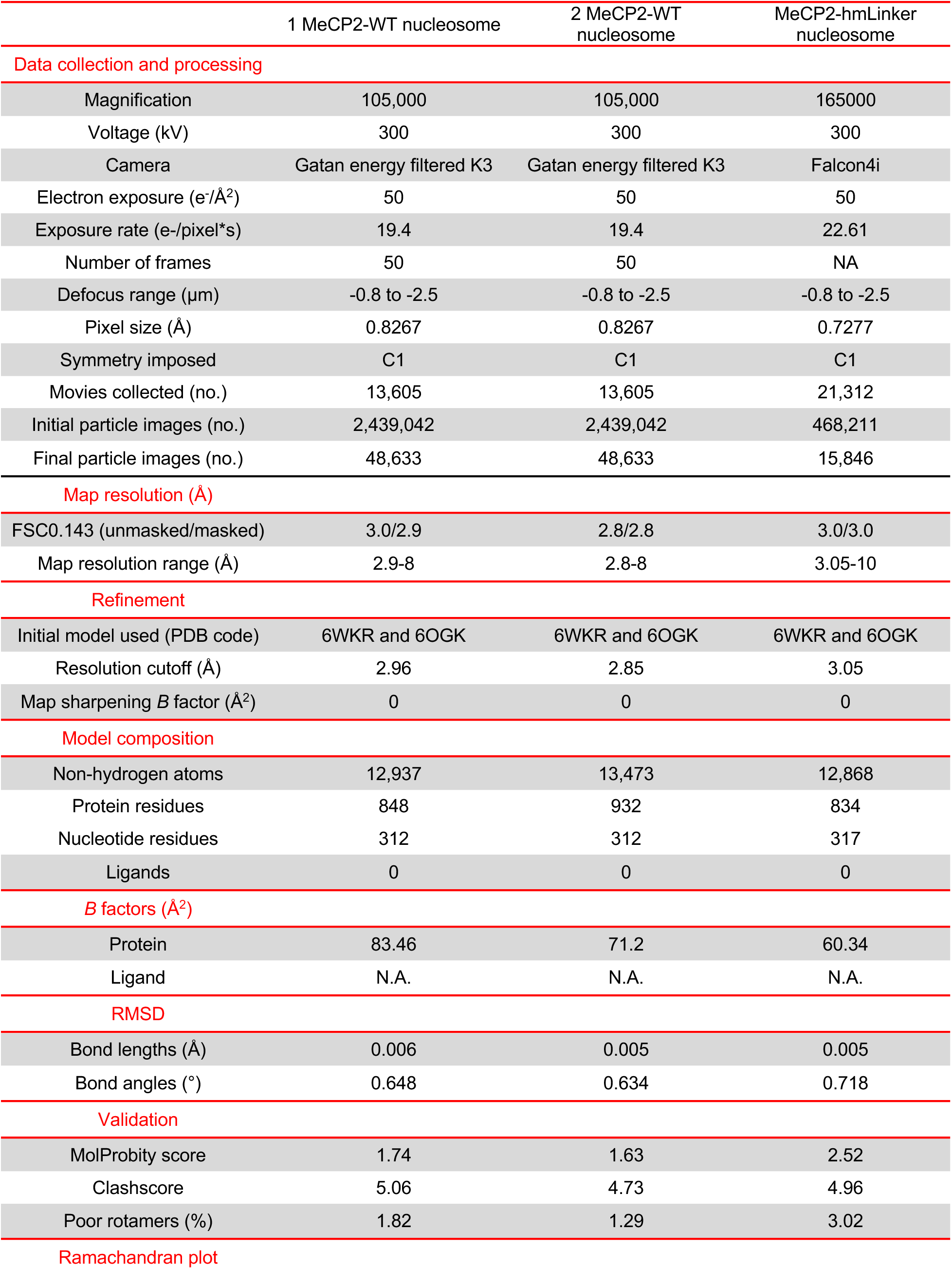
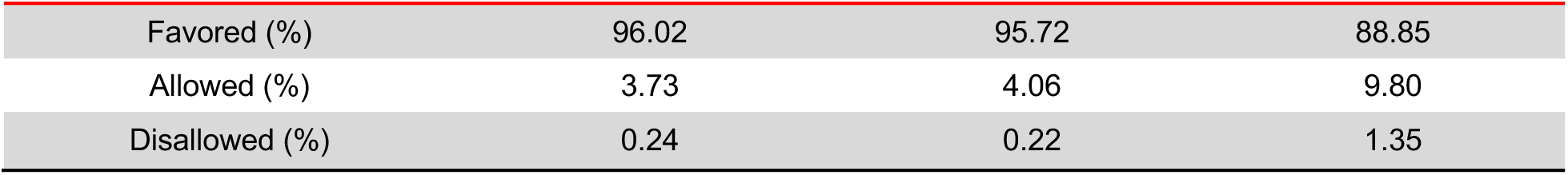
Cryo-EM data collection and refinement statistics.

|  | 1 MeCP2-WT nucleosome | 2 MeCP2-WT nucleosome | MeCP2-hmLinker nucleosome |
| --- | --- | --- | --- |
| **Data collection and processing** |  |  |  |
| Magnification | 105,000 | 105,000 | 165000 |
| Voltage (kV) | 300 | 300 | 300 |
| Camera | Gatan energy filtered K3 | Gatan energy filtered K3 | Falcon4i |
| Electron exposure (e-/Å<sup>2</sup>) | 50 | 50 | 50 |
| Exposure rate (e-/pixel*s) | 19.4 | 19.4 | 22.61 |
| Number of frames | 50 | 50 | NA |
| Defocus range (μm) | -0.8 to -2.5 | -0.8 to -2.5 | -0.8 to -2.5 |
| Pixel size (Å) | 0.8267 | 0.8267 | 0.7277 |
| Symmetry imposed | C1 | C1 | C1 |
| Movies collected (no.) | 13,605 | 13,605 | 21,312 |
| Initial particle images (no.) | 2,439,042 | 2,439,042 | 468,211 |
| Final particle images (no.) | 48,633 | 48,633 | 15,846 |
| **Map resolution (Å)** |  |  |  |
| FSC0.143 (unmasked/masked) | 3.0/2.9 | 2.8/2.8 | 3.0/3.0 |
| Map resolution range (Å) | 2.9-8 | 2.8-8 | 3.05-10 |
| **Refinement** |  |  |  |
| Initial model used (PDB code) | 6WKR and 6OGK | 6WKR and 6OGK | 6WKR and 6OGK |
| Resolution cutoff (Å) | 2.96 | 2.85 | 3.05 |
| Map sharpening *B* factor (Å<sup>2</sup>) | 0 | 0 | 0 |
| **Model composition** |  |  |  |
| Non-hydrogen atoms | 12,937 | 13,473 | 12,868 |
| Protein residues | 848 | 932 | 834 |
| Nucleotide residues | 312 | 312 | 317 |
| Ligands | 0 | 0 | 0 |
| ***B* factors (Å<sup>2</sup>)** |  |  |  |
| Protein | 83.46 | 71.2 | 60.34 |
| Ligand | N.A. | N.A. | N.A. |
| **RMSD** |  |  |  |
| Bond lengths (Å) | 0.006 | 0.005 | 0.005 |
| Bond angles (°) | 0.648 | 0.634 | 0.718 |
| **Validation** |  |  |  |
| MolProbity score | 1.74 | 1.63 | 2.52 |
| Clashscore | 5.06 | 4.73 | 4.96 |
| Poor rotamers (%) | 1.82 | 1.29 | 3.02 |
| **Ramachandran plot** |  |  |  |
| Favored (%) | 96.02 | 95.72 | 88.85 |
| Allowed (%) | 3.73 | 4.06 | 9.80 |
| Disallowed (%) | 0.24 | 0.22 | 1.35 |

**Figure 1.**
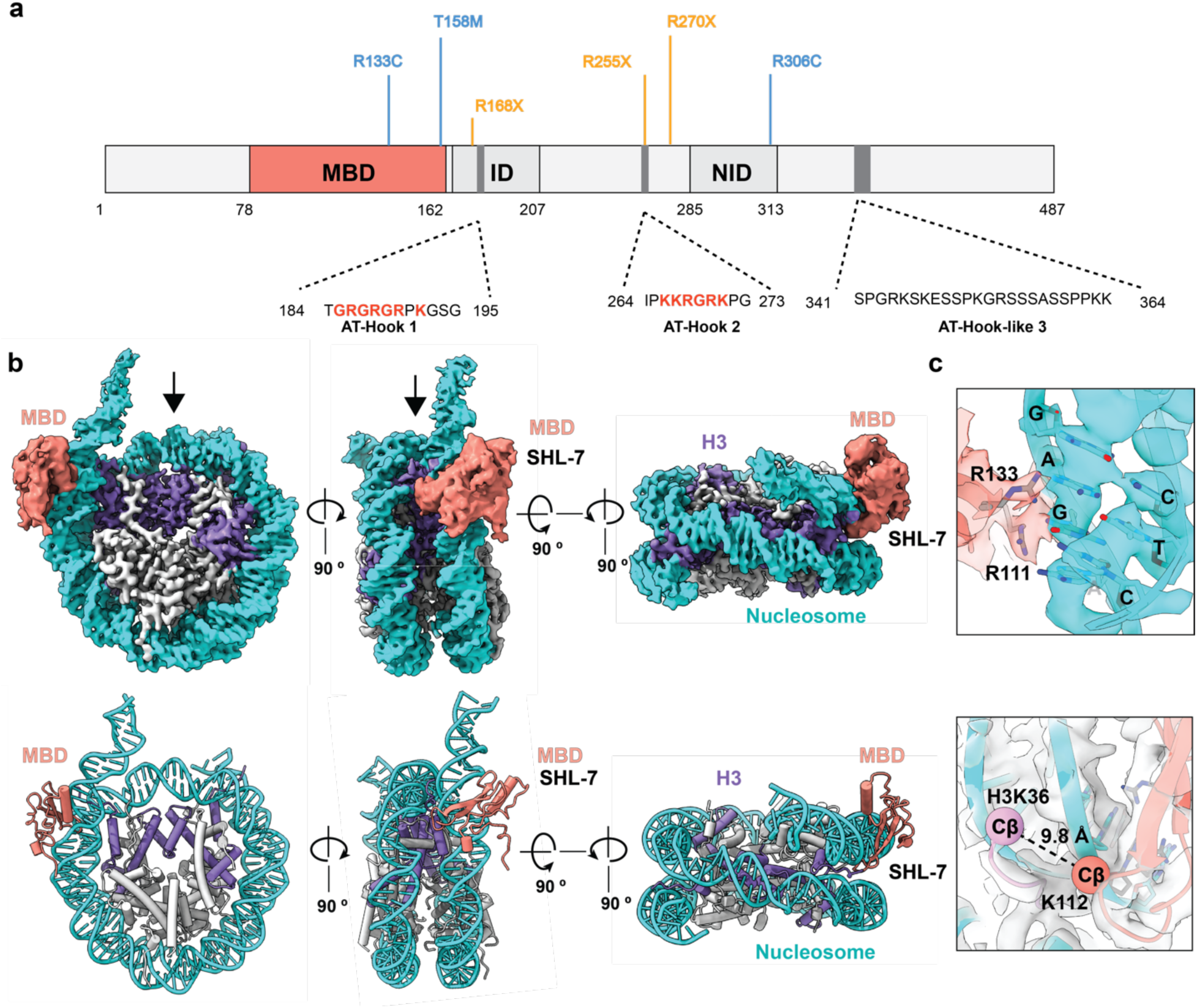
Cryo-EM structure of MeCP2 bound to an unmodified nucleosome. **(a)** Domain organization of MeCP2 (isoform e2), showing the methyl-CpG-binding domain (MBD, salmon), intervening domain (ID), NCoR/SMRT-interaction domain (NID), and AT-hook motifs (grey). The six Rett syndrome mutations characterized in this study are marked above: missense mutations (R133C, T158M, R306C) in blue and nonsense/truncation mutations (R168X, R255X, R270X) in orange. The sequences of the three AT-hook motifs are indicated. **(b)** Cryo-EM density map (top) and corresponding atomic model (bottom) of the single MeCP2(MBD)-bound nucleosome complex, shown in three orthogonal orientations. The MBD (salmon) is resolved, engaging the nucleosome near SHL7. DNA is shown in cyan, histone H3 is purple, and the remaining core histones are light grey. Arrows mark the dyad axis. **(c)** Close-up of the MeCP2 MBD–nucleosome interface. Top: the MBD arginine residues R133 and R111 contact the major groove of the nucleosomal DNA at SHL7. Bottom: van der Waals contact between histone H3K36 and MeCP2 K112 (Cβ–Cβ distance, 9.8 Å).

The reconstruction revealed two copies of MeCP2, each engaging super-helical location (SHL) ±7 near the dyad through its MBD (Extended Data Fig. 1a), a stoichiometry consistent with the complex formation observed by mass photometry (Supplementary Fig. 1). Further 3D classification revealed that one MBD domain was much better resolved, and refinement of the nucleosome with one MBD bound yielded a high-resolution map. The MBD engages the DNA major groove at SHL7, where a TCT/AGA sequence is read by the same arginine residues (R133, R111) that engage methylated CpG (Fig. 1c), with MBD K112 positioned near histone H3 K36 (Cβ – Cβ distance 9.8 Å) (Fig. 1c). This geometry resembles X-ray structures of the MBD on unmethylated DNA (Extended Data Fig 1b)^19^. AlphaFold3 prediction of MeCP2-nucleosome complex consistently positioned the MBD near the entry/exit site of nucleosomal DNA in the linker region similar to the cryo-EM structure but the prediction accuracy was poor and orientation of MBD on the DNA was different^19^ (Supplementary Fig. 4a). Furthermore, we observed additional density above the acidic patch of the nucleosome and on the linker DNA, albeit at a lower resolution threshold (Extended Data Fig. 2a, b). However, this density could not be refined at high resolution, likely due to increased flexibility. These results demonstrate that the MBD engages nucleosomal DNA through sequence and shape readout at SHL±7, with a minor histone H3 tail contact, and without requiring canonical CpG methylation for initial positioning (Fig. 1c).

To test whether positioning depends on methylation, we determined a 3-10 Å cryo-EM reconstruction of MeCP2 bound to a nucleosome containing a single hemi-methylation site (mCG(A/T)_4_) in the linker DNA 15 bp from the entry/exit site (Extended Data Fig. 3a, b, Supplemental Fig. 5, 6). We observed a clear shift in MBD density into the linker DNA by roughly one major groove (Extended Data Fig. 3c, d). However, due to the conformational flexibility of the linker DNA, we could not resolve the MBD density at high-resolution for model building. However, we were able to rigid-body dock the MBD domain into the density. The MBD geometry on the linker DNA hemi-methylation site was similar to that observed for binding to an unmodified nucleosome at SHL±7 (Extended Data Fig. 3c, d). AlphaFold3 prediction of the MeCP2(MBD)-hmLinker nucleosome, although of low prediction accuracy, placed the MBD on the linker DNA next to the hemi-methylation site (Supplementary Fig. 4a, b). Together, these data establish that the MBD anchors MeCP2 at nucleosomal SHL±7 through a sequence/shape readout independent of DNA methylation, and that linker DNA methylation is sufficient to reposition the MBD to the methylation site.

### MeCP2 binding position on nucleosomes is dictated by the location of the hemi-methylated CpG

To validate our structural findings, we performed MNase footprinting across increasing concentrations of MNase (12.5 U – 100 U) on three nucleosome: an unmodified 227-bp nucleosome (referred to as 227WTnuc; 40-N-40), a nucleosome with a single hemi-methylated CpG at SHL7 on the wrapped DNA (hmWrap; 14-N-40), and a nucleosome with a single hemi-methylated CpG in the linker DNA 15 bp from SHL7 (hmLinker; 23-N-40) (Fig. 2a,b, Supplemental Fig. 7,8). We designed these hemi-methylated nucleosomes such that the methylated CpG sits on the SHL+7 side, while the SHL-7 side lacks any CpG and therefore cannot be read as a methylation site. The hemi-methylation site matches the consensus MeCP2 site (mCG(A/T)_n_; n>3)^20^.

**Figure 2.**
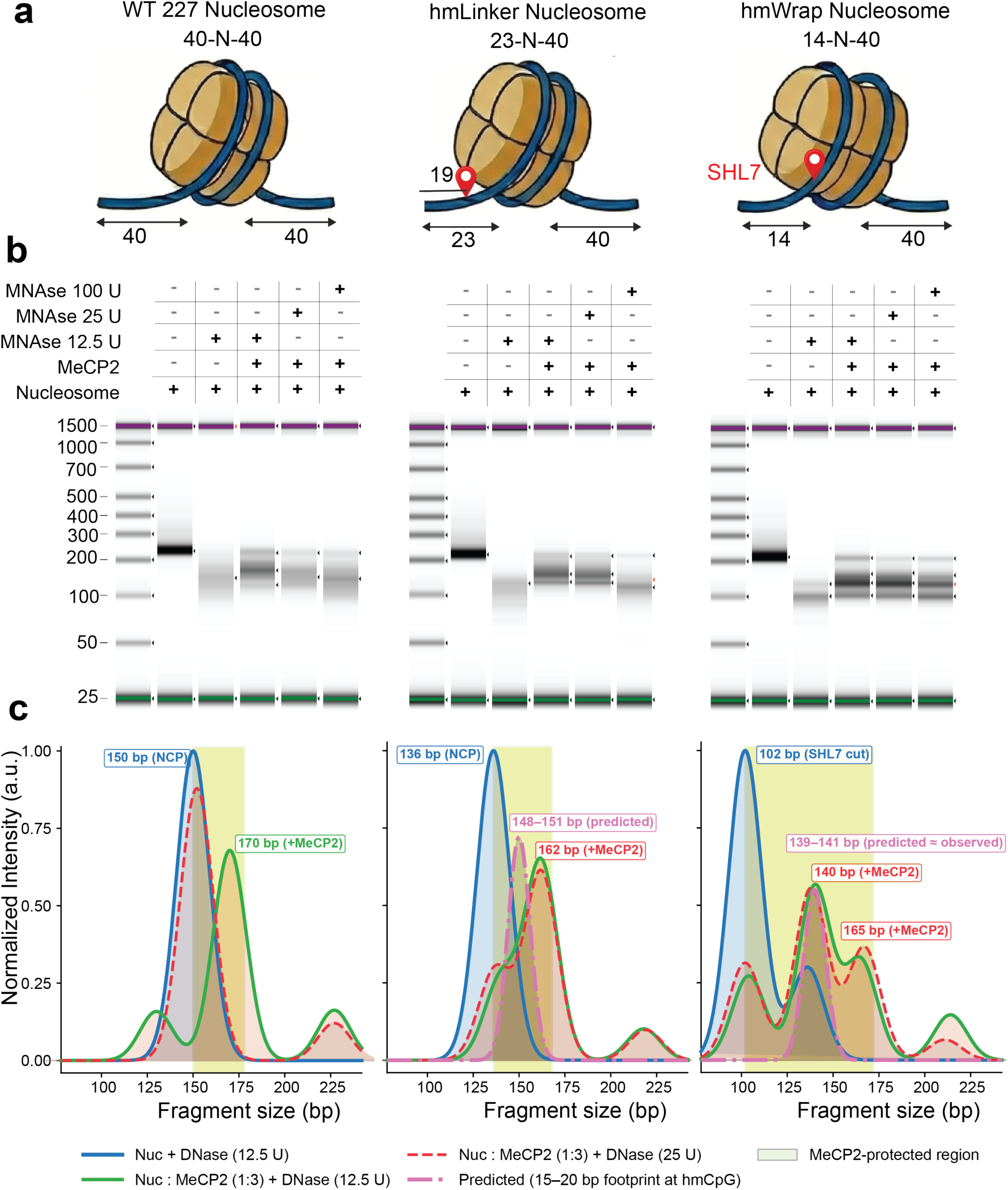
MeCP2 protects nucleosomal linker DNA from MNase cleavage. **(a)** Schematic representations of the three nucleosome substrates used throughout this study. The 227-bp WT nucleosome (40-N-40) contains 40-bp linker arms flanking the 147-bp nucleosome core particle (NCP). The hmLinker nucleosome (23-N-40) carries a single hemi-methylated CpG at position 19 within the 23-bp linker, with the opposing arm remaining 40 bp. The hmWrap nucleosome (14-N-40) is assembled on a DNA template in which the hemi-methylated CpG is positioned within the wrapped DNA at SHL 7 on the short-linker side. Hemi-methylated CpGs are indicated in red. **(b)** MNase footprinting of each nucleosome substrate in the absence or presence of saturating MeCP2. Reactions were performed at three MNase concentrations (100, 25, and 12.5 U) to identify protected regions across a range of cleavage intensities. Representative TapeStation images from two independent experiments are shown. The bands at 25 bp and 1500 bp correspond to internal TapeStation controls used for accurately measuring DNA fragment lengths. **(c)** Fragment-length distribution profiles derived from fluorescent capillary electrophoresis of MNase cleavage products. Normalized fluorescence intensity is plotted against DNA fragment size (bp). Shaded areas indicate the NCP protection boundary and MeCP2-dependent protection. The predicted footprint limits are shown for comparison with observed shifts. Data representative of two independent experiments.

In the absence of MeCP2, the 227 bp nucleosomal DNA of 227WTnuc was trimmed to give ∼ 140 bp DNA characteristic of the nucleosome core particle. On 227WTnuc, MeCP2 extended the core footprint by 20-30 bp, reflecting additional protection likely from the weak, sequence-non-specific ID/AT-hook binding to linker DNA. On hmLinker nucleosomes, MeCP2 produced a distinct, stable footprint (∼138 bp) at the linker-core boundary, and on hmWrap nucleosomes it occluded the entry/exit site (Fig. 2b). The observed footprint matches well with the predicted footprint of 137 bp when the MBD is anchored to the mCG on the linker. On hmWrap nucleosomes, MeCP2 occluded the entry/exit site, demonstrating binding to the SHL7 site (Fig. 2c). Taken together, MeCP2 binding provides robust, methyl-directed protection, suggesting that its binding position on the nucleosome is set primarily by the location of the methylated CpG.

### RTT mutations fall into three mechanistically distinct nucleosome binding classes

To investigate how RTT mutations affect nucleosome binding, we used electrophoretic mobility shift assays (EMSA) on the same three substrates, with a 27-bp unmethylated DNA competitor to reduce non-specific binding and obtain a more accurate readout (Fig. 3a, Extended Data Fig. 4, Supplementary Fig. 9-11).

**Figure 3.**
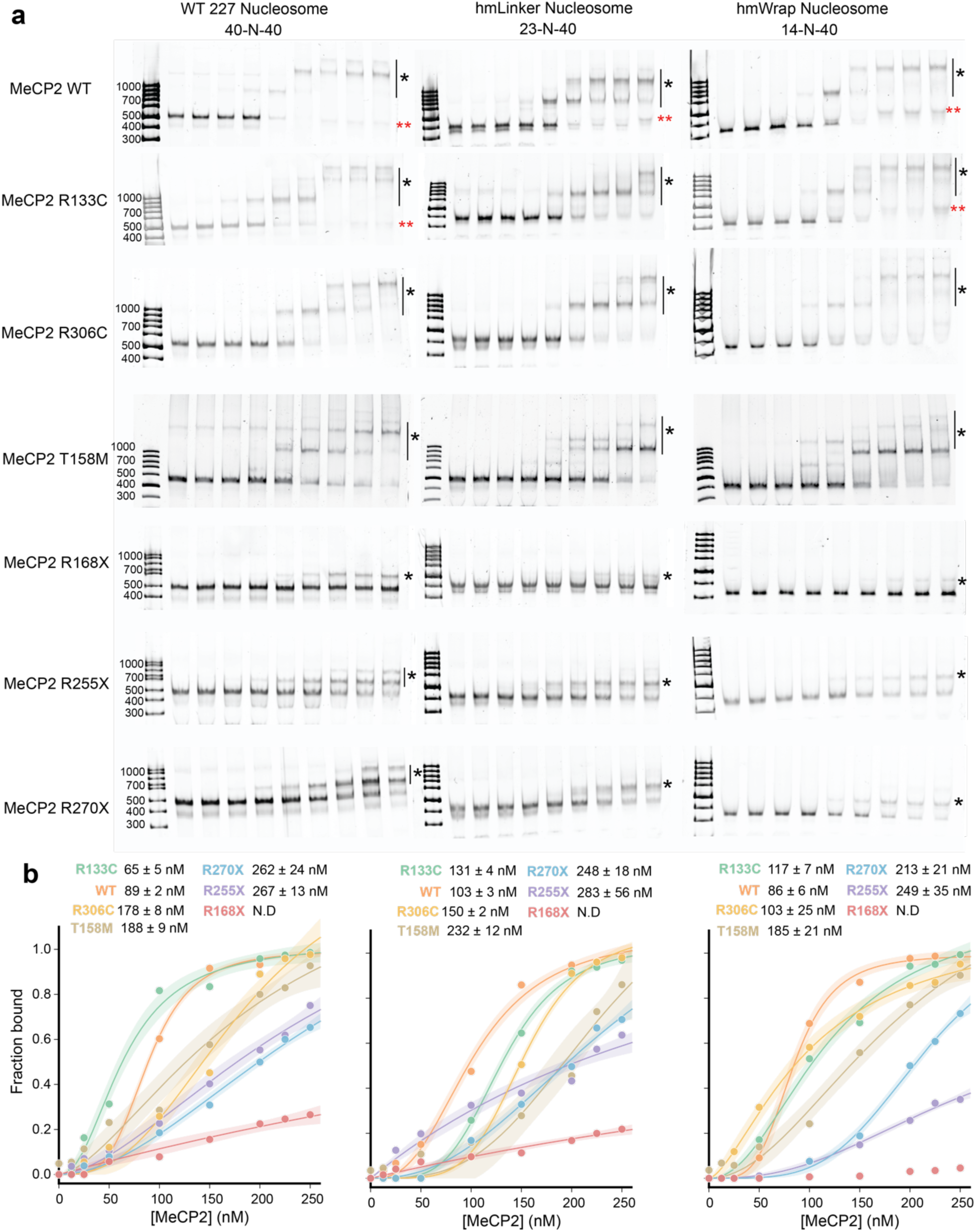
Rett syndrome variants display graded reductions in nucleosome-binding affinity. **(a)** Representative electrophoretic mobility-shift assays (EMSAs) for wild-type (WT) MeCP2 and six Rett syndrome variants (R133C, T158M, R306C, R168X, R255X, R270X) against all three nucleosome substrates (227WTnuc, hmWrap, hmLinker). Black asterisks (*) indicate MeCP2-bound nucleosome complexes, while red asterisks (**) indicate MeCP2-bound to the 27 bp non-specific competitor DNA. **(b)** Binding isotherms derived from EMSA gel quantification, grouped by nucleosome substrate (left, 227WTnuc; center, hmLinker; right, hmWrap). For each variant, points show the fraction bound (mean of three independent replicates) as a function of MeCP2 concentration, and solid lines are best-fit binding curves. Shaded regions are the 2σ (≈95%) confidence bands of each fit.

Wild-type MeCP2 bound 227WTnuc and hmWrap with similar apparent K_d_ (89 ± 2 and 86 ± 6 nM; Hill n ≈ 4) and with high cooperativity (Hill n = 4), consistent with previous observations of cooperative DNA binding^6,11^. (Fig. 3b, Supplementary Table S1). Unexpectedly, the R133C mutant bound 227WTnuc about 1.4-fold tighter than WT (65 ± 5 nM) with inverted selectivity, binding unmodified nucleosomes 1.8-fold tighter than hmWrap nucleosome. (117 ± 7 nM). Because the R133 amino acid makes a direct DNA contact in the methyl-CpG reading interface of the MBD^21^, R133C shifts MeCP2 toward a methylation-independent, shape-dependent mode rather than weakening binding.

The nonsense truncations weakened binding in proportion to C-terminal loss. R168X (NTD and MBD only) was severely impaired on all substrates (apparent K_d_ > 1 µM) with reduced Fmax, indicating near-complete loss of stable engagement and confirming the requirement for the ID and AT-hook regions for productive nucleosome binding^6^ (Fig. 3a, b). R255X, which retains the first AT-hook, and R270X, which additionally retains the N-terminal half of AT-hook 2, bound all three substrates weakly (∼ 2.5-to-3.5-fold weaker than WT). R270X showed a larger fraction of bound complexes with hmWrap compared to R255X (0.73 vs 0.35, p < 0.01), hinting at a residual preference for hmWrap due to the combinatorial role for AT-hook regions in stabilizing nucleosome engagement (Fig. 3a, b). The apparent Hill coefficients (∼1.7–1.9) for both truncations were lower than WT, consistent with a loss of CTD-dependent cooperative engagement^11,14^.

The NID missense variant R306C bound hmWrap (103 ± 25 nM) near WT (86 ± 6 nM) and retained a modest preference for hmWrap over the unmodified nucleosome, consistent with an intact MBD (Fig. 3c). T158M, the most common cause of RTT, bound all three nucleosome substrates about 2.2-to 2.4-fold weaker than WT with no methylation preference, a larger reduction than expected from its near-WT affinity for short DNA (see below), indicating that the destabilized MBD fold is more consequential on the nucleosome than on naked DNA^22–24^ (Fig. 3a, b). These data resolve the RTT variants into mechanistically distinct classes: gain of affinity with inverted selectivity (R133C), graded loss of binding with progressive C-terminal truncation (R270X > R255X > R168X), and modest, methylation-indifferent weakening from missense substitutions (T158M, R306C). Taken together, our data suggest that pathogenic mutations disrupt nucleosome recognition through several separable routes rather than a single shared defect.

### ID and AT hook contacts rather than the MBD set intrinsic DNA affinity

We next asked whether the nucleosome binding differences among MeCP2 variants reflect intrinsic changes in DNA affinity. Using fluorescence polarization (FP), we measured binding of WT MeCP2 and the six RTT variants to 27 bp or 15 bp double-stranded DNA that was unmethylated, fully methylated, or hemi-methylated at the CG(A/T)_n_ MeCP2 consensus binding sequence, each with and without the DNA-mimic competitor, poly (dI.dC) (Fig. 4a, b; Extended Data Fig. 5; Supplementary Table S2). Binding in the presence of competitor is biologically relevant, as in the cell nucleus, MeCP2 would need to discriminate between unmethylated and methylated DNA.

**Figure 4.**
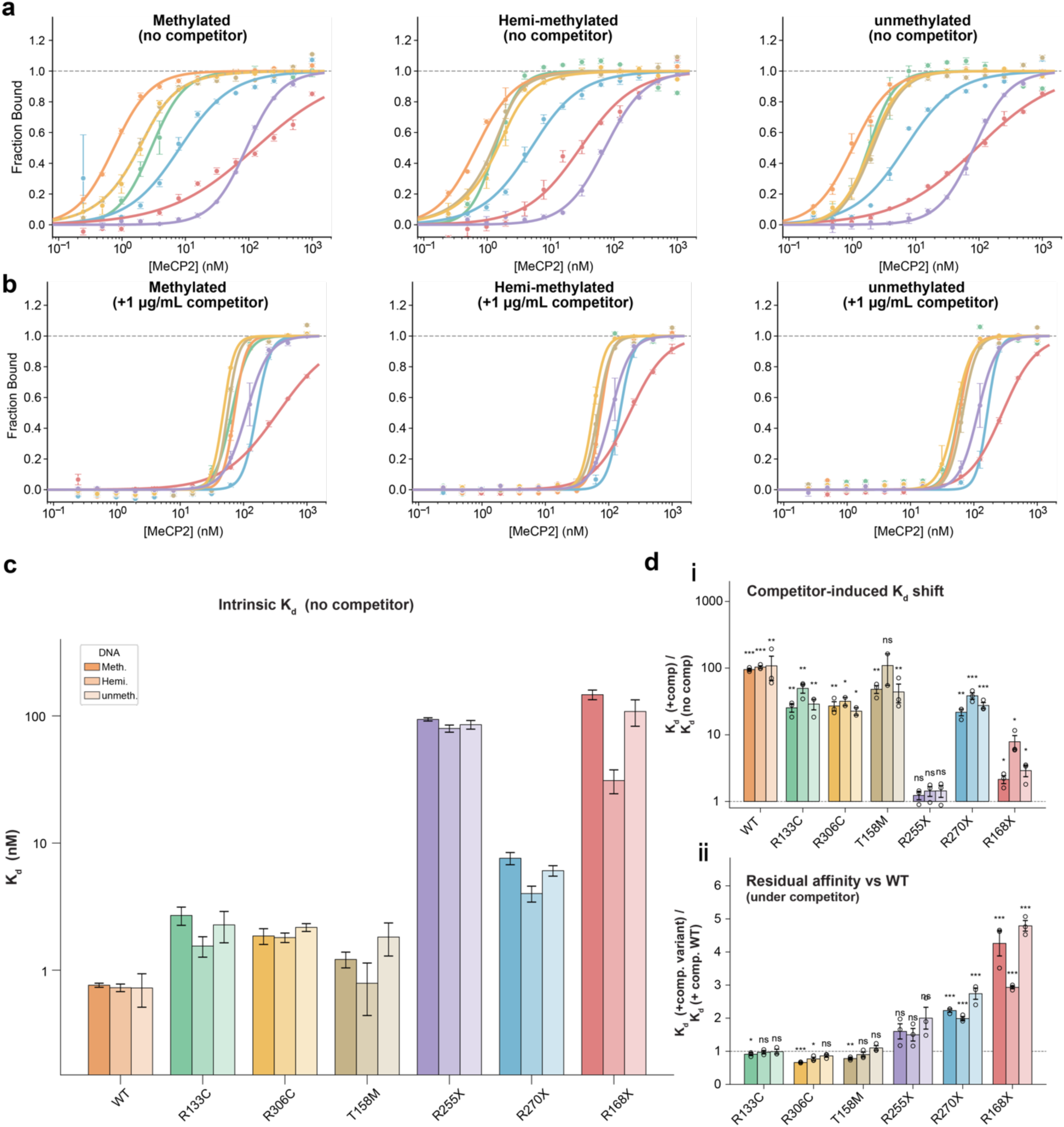
Fluorescence polarization reveals variant-specific DNA-binding affinities and altered competitor sensitivity. (a, **b)** Fluorescence polarization (FP) binding isotherms for WT MeCP2 and six Rett syndrome variants (R133C, R306C, T158M, R255X, R270X, R168X) in the absence and presence of 1 µg/mL poly(dI.dC) non-specific competitor, measured against a 27 bp FAM-labeled methylated (left), hemi-methylated (center), or unmethylated double-strand (right) DNA probe at 2 nM. Fraction bound is plotted on a linear scale against protein concentration (nM, log scale). Points represent individual replicates (n = 3), and lines show fits to the mean anisotropy trace. **(c)** Intrinsic apparent DNA binding affinities (K_d_, no competitor) for each variant and DNA type. Three grouped bars per variant represent methylated (Meth.), hemi-methylated (Hemi), and unmethylated (Unmeth) DNA. Bars show the mean K_d_ from three independent replicate fits ± s.e.m. on a log scale. **(d)** Quantitative assessment of competitor sensitivity. (**d-i**) Fold change in K_d_ upon addition of competitor (FC1 = K_d_(+comp)/K_d_(no-comp)). Three grouped bars per variant represent methylated, hemi-methylated, and unmethylated DNA. Bars are mean ± s.e.m. and individual replicate values are overlaid as dots. The statistical significance represented by * above each variant was calculated using a two-sided one-sample t-test. (**d-ii**) Residual affinity under competitor relative to WT (FC2 = K_d_(+comp, variant)/K_d_(+comp, WT)), shown as three grouped bars per variant for methylated, hemi-methylated, and unmethylated DNA. Bars are mean ± s.e.m. with replicate values overlaid as dots. FC2 above 1 indicates weaker binding than WT under competitor conditions, reflecting the residual DNA binding that remains after the competitor-sensitive contacts are suppressed. The statistical significance above each variant represented as * was calculated using a two-sided Welch t-test. ns, not significant; *, p < 0.05; **, p < 0.01; ***, p < 0.001.

Without competitor, WT MeCP2 and the three missense variants all bound 27 bp methylated DNA with similar affinity (WT 1 ± 0.1, T158M 1.2 ± 0.2, R306C 1.9 ± 0.3, R133C 2.7 ± 0.4 nM). The nonsense truncations, however, showed graded loss of affinity in proportion to the extent of C-terminal deletion. R270X was modestly weakened (8 ± 0.8 nM, about 10-fold), whereas R255X (94 ± 3 nM) and R168X (147 ± 13 nM) were severely impaired, in line with the near-complete loss of nucleosome engagement by R168X. All three truncations retain an intact MBD, so their loss of affinity cannot come from the MBD and must instead reflect removal of C-terminal DNA-contacting regions. These mainly include the intervening domain (ID; aa 163-206) and the AT-hook motifs, consistent with prior studies that assign roles for the ID in enhancing MBD-mediated DNA binding^11,25^ (Supplementary Table S3).

We then asked how each variant discriminates methylated from unmethylated DNA. On the 27 bp probe, none of the variants showed a clear preference. Discrimination appeared only on the 15 bp probe, which carries a single methyl CpG with little flanking DNA and therefore isolates the MBD readout from length-dependent nonspecific binding (Extended Data Fig. 5). On 15 bp DNA, R168X preferred methylated over unmethylated DNA by about 19-fold (16 nM vs 298 nM) and R270X by about 5-fold (1.4 nM vs 7 nM), while R255X, WT and the missense variants bound all three substrates similarly. The truncations therefore read methylation only on the minimal substrate and added flanking DNA masks that read out.

To ask how much of each protein’s binding comes from competitor-sensitive, largely non-specific DNA contacts, we measured the competitor-induced shift in K_d_ (FC1) and the apparent K_d_ under competitor relative to WT (FC2) (Fig. 4c, d; Supplementary Table S4). FC1 measures how much of MeCP2’s DNA-binding affinity comes from largely non-specific contacts that poly dI.dC competitor can strip away, while FC2 measures how much binding affinity is left after those contacts are stripped away, compared to WT MeCP2. WT showed the largest competitor effect (FC1 95 ± 4, p < 0.001). The three missense variants were 2-to-4-fold less sensitive (T158M 48 ± 6, R306C 27 ± 4, R133C 25 ± 4, all p < 0.01), so each point mutation removes part of the competitor-sensitive binding even though total affinity stays near WT. R270X grouped with the missense variants (FC1 22 ± 2, p < 0.01) rather than with the other truncations, while R255X and R168X were essentially competitor-insensitive (1.2 ± 0.2 and 2.1 ± 0.3). The break falls between R270X, which ends at residue 269 and keeps the N-terminal half of AT hook 2, and R255X, which removes it, mapping the competitor-sensitive contact to residues 255 to 269. R270X keeps a WT-like competitor response yet binds 10-fold weaker than WT, suggesting that competitor sensitivity and raw DNA affinity are distinct properties.

Comparing the with-and without-competitor conditions across all proteins provided a clear pattern. The intrinsic K_d_ on 27 bp methylated DNA spanned 193-fold (∼1 to 147 nM), but under competitor this collapsed to 6-fold (48 to 307 nM), and to 3-fold among the six proteins that retain the ID (48 to 161 nM). The intrinsic affinity ladder is therefore built almost entirely from competitor-sensitive ID and AT hook contacts, while the competitor-resistant residual is similar whenever the ID is present. FC2 tracks that residual, falling at or below unity for the missense variants (R133C: 0.9, T158M: 0.8, R306C: 0.7) and rising across the truncations (R255X: 1.6, R270X: 2.2, R168X: 4.3). R255X loses competitor sensitivity yet keeps a residual comparable to WT (116 nM), whereas R168X, the only variant lacking the entire ID, is also the only one whose residual degrades (307 nM). These ID and AT hook contacts therefore buffer MBD point mutations in the full-length protein, explaining why the isolated R133C MBD, measured previously by others, loses more than 100-fold in DNA binding and all methyl selectivity, while full-length R133C here is within 4-fold of WT^26–28^. Because the missense variants leave intrinsic DNA affinity largely intact, their nucleosome-level defect is the more likely route by which they alter MeCP2 function in cells.

### Biochemical binding properties predict variant-specific genome occupancy and transcriptional phenotypes

The mechanistic classification above makes testable predictions for how each variant could behave in neuronal cells. WT MeCP2 is expected to occupy mCpG-dense loci, including CpG islands and promoters, with high specificity. R270X, retaining the MBD and full ID, should preserve near-WT targeting with reduced overall chromatin occupancy. R168X is predicted to give the lowest specific occupancy, as the MBD alone is insufficient for stable nucleosome engagement. R133C is perhaps the most intriguing variant, as its affinity for naked DNA is near WT, and yet on nucleosomes it binds unmodified substrate 1.8-fold tighter than hmWrap and 1.4-fold tighter than WT (Fig. 5a). That inversion predicts that R133C redistributes away from mCpG-enriched targets toward non-specific loci including unmethylated chromatin, which would be mechanistically distinct from, and potentially more disruptive than, simple loss of function.

**Figure 5.**
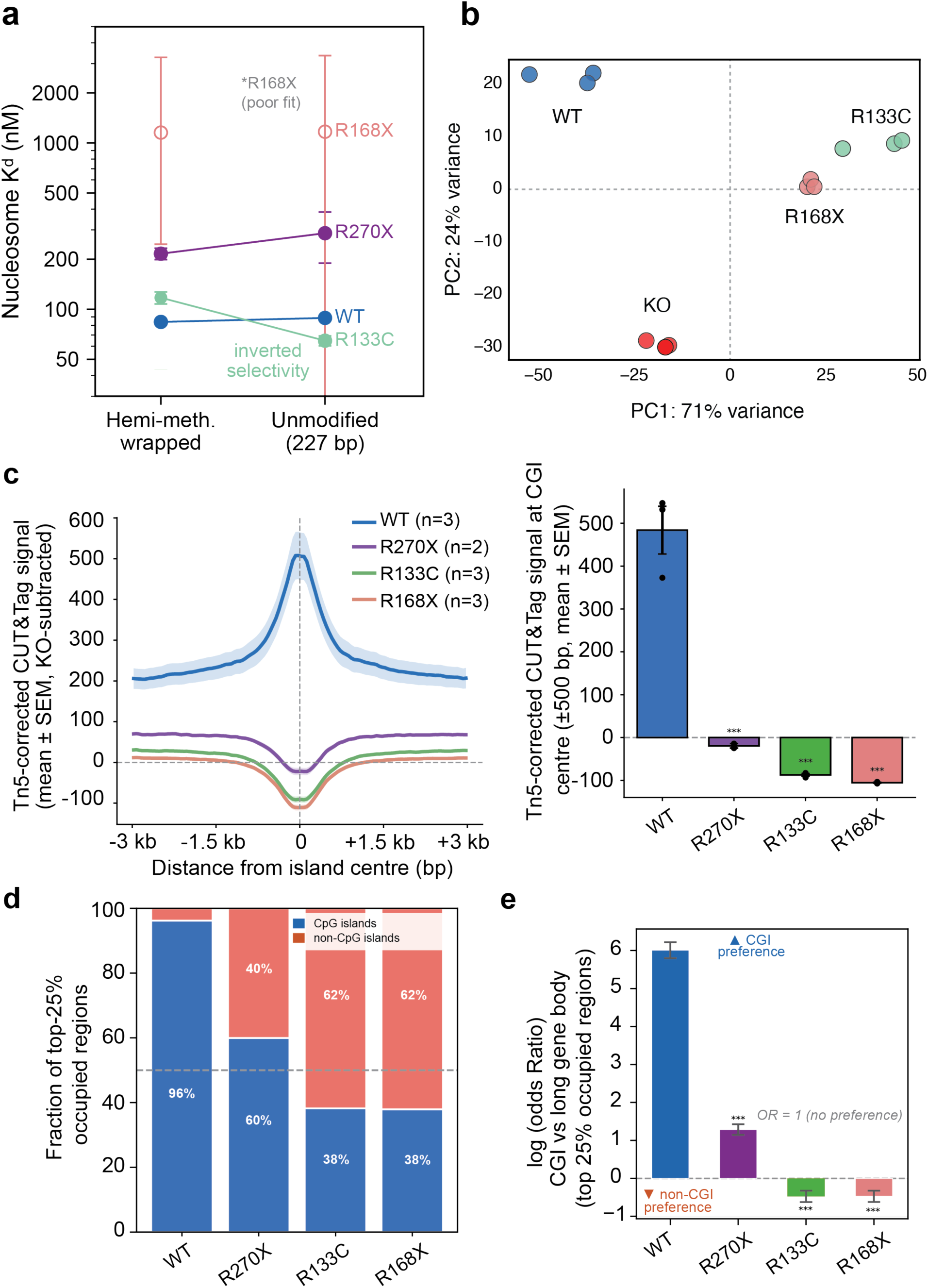
MeCP2 missense variants disrupt CpG island selectivity and nucleosomal methylation readout. **(a)** Nucleosome binding affinity (K_d_, nM) of recombinant WT (blue) and variant MeCP2 for hemi-methylated wrapped nucleosomes versus unmodified 227 bp nucleosomes. R133C (green) shows inverted selectivity, binding unmodified nucleosomes more tightly than methylated ones. Mean ± SEM; n = 3 independent experiments. **(b)** Principal component analysis (PCA) of DESeq2-normalized RNA-seq counts from hESC-neurons expressing WT (blue), R133C (green), R168X (salmon), or KO MeCP2 (red) (GSE230714). Each symbol represents one biological replicate (n = 3 per genotype). PC1 (71% variance) and PC2 (24% variance) separate genotypes along a WT–KO axis. **(c)** (left) Tn5-corrected MeCP2 CUT&Tag metagene profile centered on 3,000 CpG islands (±3 kb), KO signal subtracted; zero line indicates KO background (Tn5 accessibility). Mean ± SEM; n = 3 replicates. (right) Tn5-corrected mean CUT&Tag signal at CpG island center (±500 bp) per genotype. After KO subtraction, only WT retains a positive signal (+506.9 ± SEM); all three RTT variants fall below the KO background level, indicating their uncorrected CGI signal was primarily Tn5 accessibility rather than specific MeCP2 binding. *** p < 0.001, all variants vs. WT (Dunnett’s test). **(d)** Tn5-corrected stacked bar charts showing the fraction of each genotype’s top 25% most-occupied genomic regions that fall within CpG islands (blue) versus long gene bodies >100 kb (red), after MeCP2-KO signal subtraction (6,465 regions: 3,000 CGIs + 3,465 long gene bodies). After Tn5 correction, WT retains 96% CGI representation, R270X retains a CGI majority at 60%, while R133C and R168X shift below the pool baseline to 38%. The dashed line at 50% marks equal representation between compartments. **(e)** Tn5-corrected log₂(Odds Ratio) ± 95% CI quantifying CpG island over-representation among the top 25% most-occupied regions per genotype. After Tn5 correction, WT OR = 60.0 (log₂OR = 5.9); R133C and R168X show reduced selectivity with a shift toward long gene bodies (OR = 0.64 and 0.63); R270X retains weak CpG island preference (OR = 2.08). *** p < 0.001 for all genotypes (two-sided Fisher’s exact test vs. OR = 1).

To test these predictions, we performed new analysis of two publicly available human neuronal datasets: GSE230714, RNA-seq in hESC-derived neurons carrying isogenic MECP2-WT, -KO, -R133C, and -R168X alleles^29^; and GSE247071, a CUT&Tag experiment in the same background profiling GFP-tagged WT, R133C, R168X, R270X, and KO, with companion whole-genome bisulfite sequencing (WGBS) to validate methylation context^29^. Principal component analysis of DESeq2 spike-in normalized RNA-seq (PC1, 71% of total transcriptional variance) cleanly separated the four genotypes (Fig. 5b). WT occupied the most negative PC1 territory (∼ -35 to -52), while KO neurons separated on a distinct second axis (PC2, 24% variance). R133C neurons were the most positive on PC1 (mean score ∼ 39), more transcriptionally divergent from WT than either R168X (mean ∼21) or KO (mean -18 on PC1) along PC1. This ordering is not predicted by naked-DNA affinity, since R133C binds short DNA with near-WT affinity, but follows directly from its loss of methylation-directed nucleosome engagement, which redistributes it away from canonical mCpG sites and produces a qualitatively different, more extreme perturbation than MeCP2 absence. R168X was intermediate on PC1, consistent with the severely compromised nucleosome-binding of the MBD-only fragment.

CUT&Tag profiling of GFP-tagged MECP2 variants from Liu et al. 2024 provided direct genome-wide occupancy data linking nucleosome-binding to transcriptomic findings (Fig. 5c, Extended Data Fig. 6a, b). Using MeCP2-KO neurons as a Tn5 chromatin accessibility reference, we computed background-subtracted CUT&Tag signal at CGI centers. WT MeCP2 showed strong CGI enrichment, confirming preferential occupancy at mCpG-dense loci, whereas all three RTT variants fell at or below the MeCP2-null background (R270X near background; R133C and R168X depleted below it).

To quantify redistribution across genomic compartments, we compared occupancy at CpG islands versus long gene bodies by Fisher exact tests (>100 kb; 6,465 regions total; Fig. 5d, e; Extended Data Fig. 6a, c). Although long gene bodies carry nearly three-fold more CG methylation than CpG islands in neurons (WGBS mean mCG: 0.86 vs. 0.32; Lister et al. 2013; Gabel et al. 2015), WT MeCP2 occupancy was strongly enriched at CGIs (odds ratio (OR) = 60.05; 96.2% of top 25% occupied regions at CGIs), showing that WT does not simply track bulk methylation. All three RTT variants showed convergent loss or inversion of selectivity: R270X retained an attenuated CGI preference (OR 2.08), R133C (OR 0.64) and R168X (OR 0.63) inverted to a long gene bodies preference (Fig, 5d, e, Extended Data Fig. 6d, e). Uncorrected profiles gave the same rank order (WT > R270X > R133C ≈ R168X; Extended Data Fig. 6f, 7a, b), confirming the pattern is not an artifact of accessibility correction. Thus, pathogenic MeCP2 mutations converge on loss of CpG-island specificity in vivo, irrespective of their distinct biochemical mechanisms.

Taken together, these analyses reveal that the transcriptomic severity gradient (R133C > R168X > KO on PC1) and the convergent loss of CpG-island occupancy across all three RTT variants are not predictable from naked-DNA affinity alone but follow from how each variant engages the nucleosome. The convergence of inverted nucleosome methylation selectivity and maximal PC1 divergence in R133C defines a clear pathway, in which loss of methylation-directed nucleosome targeting drives redistribution to non-canonical loci and severe transcriptional dysregulation, even though R133C binds naked DNA with near-WT affinity. The truncations redistribute MeCP2 in the same way but they do so via different mechanisms involving progressive affinity loss, establishing that biochemically distinct defects produce a shared genome-wide phenotype.

## Discussion

The 2.9 Å cryo-EM structure reported here provides the first high-resolution view of the MeCP2 MBD positioned on a nucleosomal surface. The symmetric engagement of two MBDs at SHL±7 is notable for several reasons. First, SHL±7 presents a TCT/AGA nucleotide in the major groove at a location where DNA curvature is substantial, but the groove remains accessible, explaining how the MBD reads this site without displacing the wrapped DNA. Second, the contacts with the H3 N-terminal tail provide a minor, methylation-independent anchor, which may explain why MeCP2 shows only a modest affinity advantage for hemi-methylated over unmodified nucleosomes in our EMSA data. Third, the symmetric MBD placement is consistent with previous STEM mass analysis, which indicated that methylated 207-bp nucleosomes accommodate two MeCP2 molecules^11^. Critically, SHL±7 is not the preferred site once methylation is available. When methyl-CpG is present in the linker, the MBD leaves SHL±7 and anchors at the linker methylation site. MeCP2 therefore has two binding modes on the nucleosome: shape readout on the wrapped DNA and sequence-directed anchoring at linker methyl-CpG. MeCP2 reads the modification where it exists and falls back on DNA shape where it does not.

The mechanistic heterogeneity of RTT mutations revealed here has important implications for genotype–phenotype relationships, and our data support a classification into at least four mechanistic classes (Fig. 6). First, MBD methylation reading mutations exemplified by R133C leave intrinsic naked-DNA affinity within 2-to-4-fold of WT but disrupt methylation-directed engagement of the nucleosome. Because R133 makes a direct contact in the methyl-CpG reading interface, R133C binds unmodified nucleosomes 1.4-fold tighter than WT and acquires a 1.8-fold preference for unmodified over methylated nucleosomes that WT does not show, so it retains chromatin occupancy but engages at aberrant locations. Second, MBD stability mutations such as T158M keep near-WT DNA affinity yet pay the highest nucleosome-specific discrimination among the variants tested (Extended Data Fig. 8). Because T158M acts mainly through reduced protein levels via accelerated degradation, strategies that restore protein levels may be effective^24^. Third, progressive truncations (R168X, R255X, R270X) whose nucleosome-binding defect scales with loss of the ID, AT-hook, and CTD contacts, parallel the clinical severity gradient among nonsense mutations^30,31^. Fourth, R306C shows retained methylation preference and near-WT DNA affinity, so part of its pathology likely arises downstream of chromatin engagement, at the level of NCoR/SMRT recruitment and T308 phosphorylation^32^.

**Figure 6.**
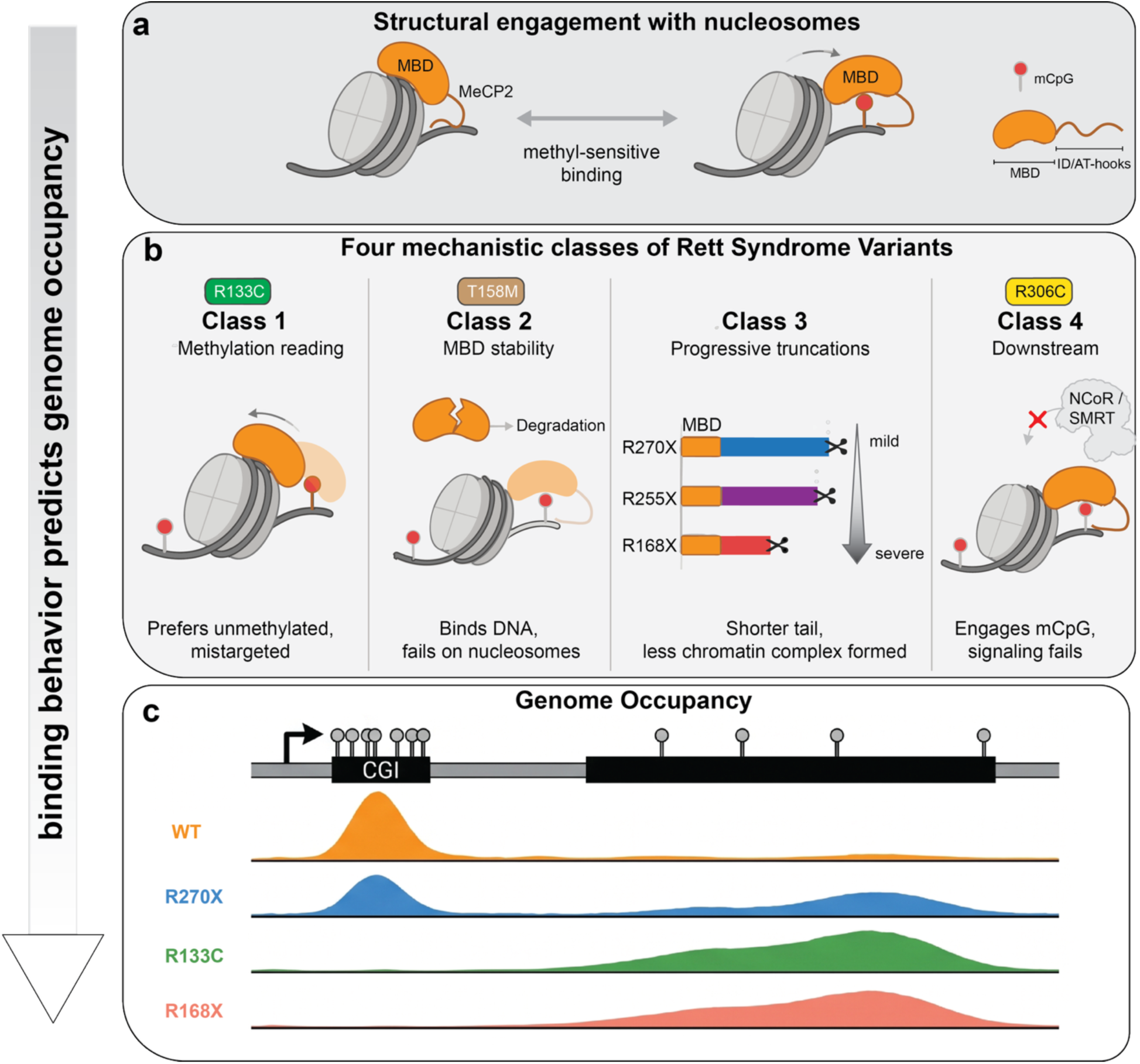
A unified model in which the DNA and nucleosome binding behavior of each MeCP2 variant predicts its genome occupancy and defines four mechanistic classes of Rett syndrome. **(a)** MeCP2 (orange, with the methyl-CpG binding domain MBD labeled, and the intrinsically disordered region and AT-hooks drawn as a tail) engages the nucleosome (grey) in a methylation-sensitive manner. **(b)** Rett syndrome variants fall into four mechanistic classes. Class 1 (R133C, green) binds DNA normally but inverts the methylation preference on the nucleosome and engages at aberrant locations. Class 2 (T158M, brown) destabilizes the MBD fold, shown as a cracked domain routed to degradation, and pays the highest nucleosome-specific cost. Class 3 (progressive truncations R270X, R255X, R168X) removes increasing lengths of the tail, and less of the nucleosome population forms a stable complex as the tail shortens. Class 4 (R306C, yellow) binds DNA normally yet fails to recruit NCoR/SMRT. MBD stability and NCoR/SMRT recruitment defect are inferred from prior studies. **(c)** These binding behaviors predict genome occupancy over a methylated gene locus with a CpG island promoter (CGI). Wild-type MeCP2 (orange) concentrates at the CGI, whereas the variants redistribute across the methylated gene body to differing extents (R270X blue, R133C green, R168X salmon).

Comparing each variant on the nucleosome against DNA of matched methylation state separates these classes by mechanism (Extended Data Fig. 8). WT loses only 1.1 to 1.5-fold on the nucleosome relative to competitor-matched DNA, so the nucleosome is as good a substrate as free DNA for a protein with the full set of contacts. T158M and R306C discriminate a further 1.6 to 2.5-fold, a nucleosome-specific defect that is invisible in the naked-DNA measurement, whereas R270X discriminates no more than WT and inherits its defect entirely from reduced DNA affinity. This further confirms that the MBD missense and truncation classes fail at different steps. R133C is the exception as it discriminates less than WT on the unmethylated nucleosome but more on both hemi-methylated substrates, which is likely what selective loss of methyl-CpG readout would look like when shape readout stays intact.

These predictions are corroborated by new analysis of two human neuronal genomics datasets. RNA-seq from isogenic hESC-derived neurons (GSE230714) showed R133C diverging from WT more than either R168X or KO. This ordering is not predicted by naked-DNA affinity, since R133C binds short DNA with near-WT affinity, but follows from the nucleosome-level data. R133C loses methylation-directed engagement and redistributes away from methylated canonical sites toward unmethylated chromatin. MECP2-KO neurons separate on a distinct axis (PC2, 24% variance), showing that a mutant MeCP2 that mistargets chromatin produces a qualitatively different perturbation than absence of the protein.

Our analysis of CUT&Tag data (GSE247071) confirmed loss of CpG-island occupancy across all three RTT variants, with R133C and R168X redistributing toward long gene bodies (38% of top-occupied regions at CGIs, vs 96% for WT and 60% for R270X). Two biochemically distinct defects, loss of methylation-directed nucleosome targeting in R133C and progressive loss of affinity in R168X and R270X, therefore converge on similar losses of CpG-island occupancy and similarly dysregulated transcriptomes and may require different therapeutic approaches.

Our work also highlights new questions. Future studies are needed to address how MeCP2 engages nucleosomes carrying mCA modifications, how post-translational modifications, including phosphorylation at S80, S274, T308, and S421^33^, modulate nucleosome affinity and selectivity, and how H1 and MeCP2 competition plays out at specific neuronal loci. Our results also suggest that naked-DNA affinity is a poor predictor of the cellular phenotype, since R133C binds short DNA with near-WT affinity yet produces the most extreme transcriptional departure from WT. Assays that report methylation-directed engagement in the nucleosome context are therefore likely to be more informative for interpreting novel MECP2 missense variants of unknown significance identified in clinical sequencing.

In summary, this work provides a structural and quantitative framework for understanding MeCP2 chromatin engagement and reveals that RTT mutations compromise this engagement through mechanistically distinct pathways. Our analysis of neuronal gene expression and chromatin occupancy data shows that these different biochemical defects lead to the same outcome. All three variants lose occupancy at their normal targets and disrupt gene expression, but they redistribute differently, which the CpG island and long gene bodies comparison separates. These findings suggest that effective therapeutic intervention may require mutation-specific strategies targeting different aspects of MeCP2 function.

## Materials and Methods

### Protein expression and purification

The MeCP2 genes were ordered from Addgene (Plasmid #48078) and cloned into a bacterial expression vector containing an N-terminal StrepII tag. All the mutations and truncations except T158M were made by Q5^®^ Site-Directed Mutagenesis Kit (NEB, E0554S). The MeCP2 T158M gene was ordered from Genewiz and cloned into the same bacterial expression vector. To purify the MeCP2 protein, mutations and truncations, the plasmid was transformed into Rosetta (DE3) competent cells. A few colonies were picked and cultured in 2 mL LB media with 1 mM Kanamycin and 0.5 mM Chloramphenicol. When the tube became turbid, the 2 mL culture was expanded to 50 mL and cultured until the OD reached 0.4. The 50 mL culture was later expanded to 100 mL. Once the OD reached 0.4, 100 mL of cells was expanded into 1.5 L of TB medium containing 1 mM Kanamycin and 0.5 mM Chloramphenicol. When the OD was 0.4, 1 mM IPTG was added, and the protein was expressed at 37 °C for 3 hours. The cells were harvested, flash frozen in liquid nitrogen, and stored in a -80 °C freezer.

To purify the protein, the frozen cells were thawed on ice and resuspended with resuspension buffer (25 mM HEPES, pH 8.0, 500 mM NaCl, 2.5 mM MgCl_2_, 10% glycerol, 10 uM Leupeptin, 1 uM Pepstatin, 0.2 mM PMSF. 1µM aprotinin, 1 Protease inhibitor tablet (Sigma, 11836170001), 1 mM DNASE, 5 µL Salt Active Nuclease (Sigma, SRE0015-5KU), and slowly rotated in the cold room for 30 min. After incubation, 1 mM TCEP and 0.05% NP40 were added and sonicated with a cycle of 10s on, 20s off for 15 min. Cell debris was then removed by centrifugation at 14,000 rpm for 30 min. The supernatant was incubated with Strep-Tactin®XT 4Flow® resin (IBA, 2-5010-025) for 40 min, and the resin was loaded into a column and allowed to settle for 30 min. The resin was washed with 6 CV of Wash buffer 1 (25 mM HEPES, pH 8.0, 500 mM NaCl, 10% glycerol, 1/3 Protease inhibitor tablet, 1 mM TCEP), 6 CV of Wash buffer 2 (25 mM HEPES, pH 8.0, 1 M NaCl, 10% glycerol, 1/3 Protease inhibitor tablet, 1 mM TCEP), and 6 CV of Wash buffer 1. The protein was eluted with 3 CV of Wash buffer 1 + 1X Buffer BXT (10X) (IBA, 2-1042-025). To the elution buffer, a final concentration of 10 mM ATP and 20 mM MgCl_2_ was added, and the sample was subjected to dialysis in dialysis buffer (25 mM HEPES, 300 mM NaCl, 10% glycerol, 5 mM 2-mercaptoethanol) for 1 hour. The AKTA-FPLC system was used for subsequent purification with a HiTrap Heparin HP column (Cytiva, 17040703) and a Superdex 200 Increase 10/300 column (GE Healthcare, 28990944). The Heparin column was equilibrated with buffer 1 (25 mM HEPES, pH 8.0, 250 mM NaCl, 10% glycerol, 1 mM TCEP), and the sample was eluted with a linear gradient of buffer 2 (25 mM HEPES, pH 8.0, 2 M NaCl, 10% glycerol, 1 mM TCEP). The Superdex 200 Increase 10/300 column was equilibrated and run with the final storage buffer (25 mM HEPES, pH 8.0, 500 mM NaCl, 20% glycerol, 1 mM TCEP). The protein was flash-frozen in liquid nitrogen as single-use aliquots and stored at -80 °C. Since the mutations and truncation variants were prone to degradation, all purification steps were performed on the same day.

### Histone purification and nucleosome reconstitution

The purification of individual histones and octamer refolding was carried out as previously described^34^. Briefly, individual lyophilized histones were resuspended in Unfolding buffer (6 M GuHCl, 20 mM HEPES, pH 7.5, 5 mM TCEP) to a final concentration of 2 mg/mL. After incubating the histones in the unfolding buffer for 1 hour at room temperature, the absorbance at A280 was measured for each histone and used to calculate the molar concentration. Each histone was then added in equal molar ratios and diluted to a final protein concentration of 1 mg/mL in Refolding buffer (10 mM HEPES, pH 7.5, 2 M NaCl, 1 mM EDTA, 5 mM 2-mercaptoethanol). The 1 mg/mL mixture was then transferred to a 10 kDa MWCO dialysis membrane and dialyzed against 600 mL of Refolding buffer for 6 hours at 4 °C, and the dialysis was repeated three times. The sample was then recovered from dialysis and centrifuged to separate any precipitated material. The supernatant was then concentrated to 500 µL using a 10 kDa MWCO AMICON spin column (Milipore, UFC5010). The concentrated sample was then loaded onto a Superdex 200 Increase 10/300 column equilibrated with refolding buffer plus 20% glycerol. Fractions containing intact, refolded octamers were pooled, concentrated, flash-frozen in liquid nitrogen, and stored at -80 °C.

### Generation of Nucleosome DNA

Plasmids containing 227bp WT mono-nucleosome DNA, hmLinker mono-nucleosome DNA, and hmLinker mono-nucleosome DNA were purchased from Twist Bioscience. Nucleosomal DNA was amplified and isolated using polymerase chain reaction (PCR) using the primers listed in the Supplementary Information.

The PCR products were purified on a Resource Q column (Cytiva, 17117901) and eluted with a salt gradient from 150 mM to 2 M KCl. Peak fractions were ethanol precipitated and resuspended in 2 M KCl buffer.

### Generation of Nucleosome Substrates

For the WT 227bp mono-nucleosome, 227bp DNA was combined with the histone octamer in a 1:1.1 (DNA to octamer) ratio. For the hemi-methylated Linker and wrap nucleosomes, a ratio of 1.0:1.0 was used. First, the appropriate amount of DNA is added to RB high buffer (10 mM HEPES, pH 7.5, 2 M KCl, 1 mM EDTA, 1 mM TCEP) and allowed to sit at room temperature for 30 min before adding the appropriate amount of octamer. The mixture is left to sit at room temperature for ∼30 min. After incubation, the sample is transferred to a 10 kDa MWCO dialysis cassette and dialyzed in RB high buffer against a linear gradient of RB low buffer (250 mM KCl, 10 mM HEPES pH=7.5, 1 mM EDTA, 1 mM TCEP) for at least 16 hours. Next, the reaction is dialyzed against RB low buffer for 4 hours, followed by a dialysis against HCS buffer (20 mM HEPES, pH=7.5, 1 mM EDTA, 1 mM TCEP) for 4 hours. The reconstituted WT, hmWrapped, or hmLinker nucleosome was then recovered and checked on both a 4% Native TBE PAGE gel and an 18% SDS PAGE gel. Nucleosomes are stored at 4°C.

### Mass Photometry

0.5 µM of nucleosomes and 1.5 µM MeCP2 are crosslinked with 0.05% glutaraldehyde in MP buffer (25 mM HEPES, pH 7.5, 50 mM NaCl, and 1 mM TCEP) for 10 min at room temperature. A final concentration of 25 mM Tris (pH 7.5) is used to quench the reaction. Samples are diluted to 20-100 nM nucleosomes in MP buffer. Mass photometry measurements were performed using a TwoMP mass photometer (Refeyn LTD, Oxford, UK). Data were acquired using the Acquire MP software package and analyzed using the Discover MP software, both from Refeyn. MP buffer was prepared fresh on the day of the experiment and checked by Mass Photometry for cleanliness. Glass coverslips (24 × 50 mm, Thorlabs Inc. 71861-054) were washed 3 times with Milli-Q water and HPLC-grade isopropanol, then dried with a clean stream of compressed air. Sample chambers were assembled by placing clean 6-well silicon gaskets (Fisher Scientific, NC2754003) on the cleaned coverslip, with the gasket positioned on the stage of the mass photometer. 20 uL of the sample was applied, and the mass photometer was focused. Data acquisition was started immediately to record a 60-second movie. The mass calibration was achieved using beta-amylase (Sigma-Aldrich, A3176, 10 nM in MP buffer; 56, 112, and 224 kDa). All measurements were performed at room temperature.

### EM sample preparation

Quantifoil Au 1.2/1.3 grids were converted to streptavidin-affinity grids using protocols described previously^17,18^. Grids were rehydrated in rehydration buffer (25 mM HEPES, pH 8.0, 50 mM NaCl, 1 mM TCEP) at room temperature for 1 hour. To assemble the MeCP2-nucleosome complex, 0.5 µM of nucleosome and 1.5 µM MeCP2 are crosslinked by 0.01% of glutaraldehyde at room temperature for 15 min. The reaction was quenched by 25 mM Tris, pH 7.5. The reaction was further diluted to a final concentration of 125 nM nucleosome with EM preparation buffer 1 (25 mM HEPES, pH 8.0, 50 mM NaCl, 2.5% glycerol, 1 mM TCEP) before use. After blotting away the remaining buffer with Whatman filter paper, 4 µL of MeCP2-nucleosome complex was loaded onto the grid. The grid was incubated for 15 min in a humidified chamber, washed twice with 50 µL of EM preparation buffer 1, and then washed with 50 µL of EM preparation buffer 2 (25 mM HEPES, pH 8.0, 50 mM NaCl, 2.5% glycerol, 1 mM TCEP, 0.01% NP40). After washing, the buffer was quickly blotted away with Whatman filter paper (cytiva, 1001-150), and 4 µL of EM preparation buffer 2 was added immediately. The grid was transferred to the Leica EM GP2 plunge freezer, blotted for 2-3 s at 10 °C and 90% humidity.

### EM data collection and processing

The Cryo-EM data of MeCP2-WT nucleosome were collected using a Titan Krios G3i equipped with a Gatan energy-filtered K3 camera and an energy filter set with a 10-eV slit width. Data acquisition was performed using SerialEM version 4.2.1 at 105,000x magnification and super-resolution mode (0.41335 Å/pixel) with a defocus range of −2.5 to −0.8 μm. Movies were collected in TIF format with a total dose of 50 electrons per square angstrom (e^−^/Å^2^) and an exposure time of 1.76 s.

The Cryo-EM data of MeCP2-hmLinker nucleosome were collected using a Titan Krios equipped with an energy-filtered Falcon4i camera. Data acquisition was performed using EPU version 3.16 at 165,000x magnification with a defocus range of −2.5 to −0.8 μm. Movies were collected in EER format with a total dose of 50 electrons per square angstrom (e^−^/Å^2^) and an exposure time of 4.62 s.

The data were processed in CryoSPARC^35^. Gain correction was applied during motion correction using Patch Motion Correction. The background streptavidin lattice of each micrograph was subtracted using in-house scripts^18^. Blob Picker is used for initial particle picking, followed by 2D classification to remove junk particles. Good 2D classes were used as templates for template picking. Template-picked particles were subjected to 2D classification, heterogeneous refinement, and non-uniform refinement. Global CTF refinement, Local CTF Refinement, and Subset particles are used to further filter particles. Several rounds of NU-Refine and 3D Variability are used to filter homogeneous particles. Local resolution filtering is used for the final map.

### Model building

The nucleosome is built using the nucleosome from PDB ID: 6WKR as a reference and the density map from our data collection^36^. The MBD domain is modeled using PDB ID: 6OGK as a reference^19^. The details of the structures are adjusted and rebuilt in the new map using COOT^37^. The MeCP2-nucleosome model was subjected to global refinement and minimization in real space using PHENIX^38^. The cryo-EM density maps and the molecular graphics were prepared with Chimera and ChimeraX^39^.

### AlphaFold 3 Prediction

MeCP2, two copies of each histone, and the nucleosome DNA sequence were provided to the AlphaFold web server for structure prediction^40^. The model graphics were prepared with ChimeraX^39^.

### Electrophoretic mobility shift assay (EMSA)

4% native TBE gels were cast using a 37.5:1 polyacrylamide solution. Gels were pre-run in 0.5X TBE buffer at 3 watts in an ice bath for 45 minutes prior to sample loading and production runs. Increasing concentrations of recombinant MeCP2 (0, 12.5, 25, 50, 100, 150, 200, 225, and 250 nM) were incubated with a fixed concentration of assembled nucleosome substrate (50 nM) and 250 nM 27 bp nonspecific dsDNA competitor in EMSA reaction buffer (25 mM HEPES, pH 7.5, 100 mM NaCl, 1 mM TCEP) for 30 min at room temperature. A final concentration of 4% sucrose was added to each sample before loading onto the pre-run 4% native gel. The gel was run at 3 watts for 90 minutes while in an ice bath. The Native gels were stained in 1x SYBR Gold Nucleic Acid Gel Stain (Invitrogen, S11494) for 20 minutes at room temperature and then imaged using a GelDoc Imaging System (Bio-Rad).

Three nucleosome substrates were used: WT 227 nucleosome (40-N-40 template, unmodified CpG): standard nucleosome; hmLinker nucleosome (23-N-40 template): hemi-methylated linker DNA, and hmWrap nucleosome (14-N-40 template): hemi-methylated wrapped DNA. Bound and free nucleosome species were resolved by native PAGE and quantified by band densitometry. The *fraction bound* at each MeCP2 concentration was calculated as [bound] / ([bound] + [free]). Three independent replicate experiments were performed for each variant x substrate combination, and the data from all replicates were pooled for model fitting. Binding isotherms for three independent replicates were fit to the Hill equation or, for weak-binding variants, to a Morrison tight-binding model. Each replicate was fit independently. For every variant and substrate combination, the binding isotherm was fit to both a Hill occupancy model and a Morrison tight-binding model, and the model was selected by the Akaike Information Criterion. We report apparent dissociation constants (apparent K_d_) since EMSA is performed under non-equilibrium electrophoretic conditions and in the presence of a non-specific competitor. The Hill or Morrison tight-binding model was selected for each curve by the Akaike Information Criterion (ΔAIC < −2 to prefer Hill).

Hill Equation:

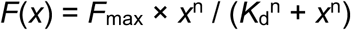

where *x* is the total MeCP2 concentration (nM), *F*_max_ is the maximum fraction bound, *K*_d_ is the apparent dissociation constant (nM), and *n* is the Hill coefficient.

Morrison Equation:

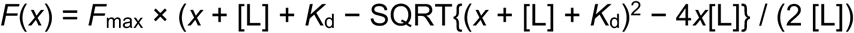

where *x* is the total MeCP2 concentration (nM), [L] = 50 nM is the fixed nucleosome concentration, *F*_max_ is the maximum fraction bound, and *K*_d_ is the dissociation constant (nM).

Non-linear least-squares fitting was performed using the Levenberg–Marquardt algorithm as implemented in *scipy.optimize.curve_fit* (Python 3; SciPy). Fraction-bound values from all replicates were concatenated into a single pooled dataset (i.e., each replicate contributed its 9 data points), and fitting was performed on the pooled data. This approach weights each replicate equally and provides a full residual distribution for parameter estimation. Initial parameter estimates were *F*_max_ = 0.95, *K*_d_ = 80 nM, *n* = 1.0 (Hill) and *F*_max_ = 0.95, *K*_d_ = 80 nM (Morrison). Both models were fit to every replicate dataset. Model selection used the Akaike Information Criterion (AIC), computed as:

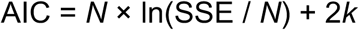

where *N* is the number of data points, SSE = Σ(*y*_i_ − *f*(*x*_i_))^2^ is the sum of squared residuals, and *k* is the number of free parameters.

The Hill model was selected when:

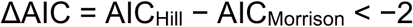

Otherwise, the Morrison equation was used. The threshold −2 penalizes the additional free parameter of the Hill model unless it provides a meaningfully better fit, consistent with standard AIC practice for model comparison differing by one parameter.

Shaded bands around fitted curves represent 2σ (95%) pointwise confidence intervals, propagated analytically from the parameter covariance matrix Σ returned by the non-linear fitting routine. At each concentration point *x,* the variance of the model prediction is:

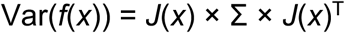

where *J*(*x*) is the row-vector of partial derivatives of the model with respect to each parameter, evaluated numerically by central finite differences at *x*. The 2σ band is:

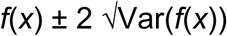

This propagation produces bands that faithfully reflect parameter correlation (e.g., K_d_ and F_max_ are typically anti-correlated) and scale appropriately with the curvature of the binding isotherm, narrowest at the half-saturation point and widest at the extremes.

### Fluorescence polarization (FP)

The FAM-labeled forward and reverse 27 bp or 15 bp DNA sequences with and without methylation are ordered from Genewiz. The dry powder is dissolved in Nuclease-Free water (Growcells.com, NUPW-0050). 20 µM of forward and reverse DNA were prepared with annealing buffer (20 mM Tris, pH 7.5, 50 mM NaCl). The DNA was heated at 95 °C for 5 min and then slowly cooled down to room temperature for annealing. The annealed double-stranded DNA was stored at -20°C until use. To set up FP reactions, 40 µL reactions were prepared in FP buffer (20 mM Tris, pH 7.5, 100 mM NaCl) containing FAM-labeled 15-bp or 27-bp DNA (unmethylated, hemi-methylated, or methylated). MeCP2 variants were titrated (0 to ∼1,000 nM, 14 concentration points spaced approximately log-linearly) against a fixed concentration of 2 nM fluorescently labeled double-stranded DNA probe. For the reactions with competitors, 1 µg/mL poly(dI·dC) (Sigma, P4929-5UN) is used as a non-specific competitor. The reactions were incubated for 30 min at room temperature in a black 384-well microplate (Corning, 3575) and then analyzed using a TECAN Spark microplate reader. Fluorescence anisotropy was measured in triplicate wells at each titration point with excitation of 481 *±* 20 nm and emission of 526 *±* 20 nm.

The observed change in anisotropy (Δr) was defined as:

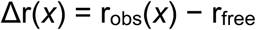

where r_free_ is the anisotropy at zero protein (mean of all zero-concentration replicates). Models were fitted directly to Δr rather than to r_obs_, absorbing the zero-point offset into the baseline.

The same two models used for EMSA data were applied, reparametrized in terms of Δr:

Hill equation:

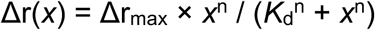

Morrison tight-binding equation:

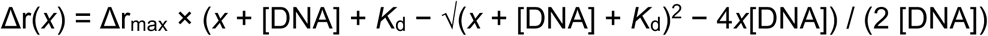

where [DNA] = 2 nM is the fixed probe concentration, Δr_max_ is the maximum anisotropy change at saturation, *K*_d_ is the dissociation constant, x is the MeCP2 concentration, and *n* is the Hill coefficient. Fraction bound was calculated for visualization by normalizing observed Δr by the fitted Δr_max_. For WT and the missense variants, the 27-bp and 15-bp K_d_ values sit at or below the 2 nM FAM probe concentration and are upper bounds. The 15-bp substrate is used to compare methylation states for the weaker-binding truncations, where the values sit clearly above the probe floor. The wide error bars on the missense FC1 values are informative rather than noise. For WT and the missense variants, the no-competitor K_d_ sits at or below the 2 nM probe floor, where it is only weakly constrained. Small replicate differences at the floor produce large swings in the ratio, so the FC1 spread is driven by a few high replicate values (the mean exceeds the median about three-fold for R133C, R306C, and T158M, versus near-equal for WT). The truncations bind well above the floor, so their no-competitor K_d_ is well determined, and their FC1 errors are correspondingly small. The large missense errors are thus a direct consequence of these proteins binding too tightly to resolve without competitor.

### MNase Footprinting

Protocol adapted from previous literature^41^, 0.5 μM dsDNA or nucleosome was combined with 1.5 μM MeCP2 and incubated at room temperature for 30 minutes in binding buffer (100 mM NaCl, 25 mM HEPES, pH = 7.9, 5 mM MgCl2, 1 mM TCEP). A control of just 0.5 μM dsDNA or nucleosome with no MeCP2 was also performed. The reactions were then diluted into MNase activity buffer (100 mM NaCl, 25 mM HEPES, pH = 7.9, 5 mM MgCl_2_, 5 mM CaCl_2_). 0.1 mg/mL BSA was then added to the reaction, which was then incubated at room temperature for an additional 15 minutes. The corresponding Units of MNase were added and allowed to digest for 10 minutes at 37℃, before quenching with an excess of EDTA.

To remove the protein for downstream DNA analysis, samples were treated with 20 μg of Proteinase K (Thermo Scientific, EO0491) at 55℃ for 30 minutes. The DNA was then purified using a NucleoSpin Gel and PCR Clean-up kit (Macherey-Nagel, 740609.250). The purified DNA was then quantified on the NanoDrop to give a rough ng/μL readout of the concentration, before being diluted to the appropriate range for quantification on the TapeStation (Agilent).

Once diluted, 1 μL of each DNA sample was mixed with 3 μL of the high-sensitivity D1000 sample buffer (Agilent, 5067-5585). A ladder was also run alongside each set of samples and prepared similarly: 1 μL of high-sensitivity D1000 Ladder (Agilent, 10597012) and 3 μL of high-sensitivity D1000 sample buffer. Data was analyzed within the TapeStation software.

## Re-analysis of published neuronal transcriptomic and genomic occupancy datasets

### Data acquisition

RNA-seq raw-read FASTQ files representing four isogenic hESC-derived neuronal genotypes (MECP2-WT, -KO, -R133C, -R168X; n = 3 biological replicates each) were obtained from GEO accession GSE230714 (Liu et al., Neuron 2024). Data was re-analyzed as specified in Liu et al., 2024, but using Hisat2 v2.1.0 instead of STAR for read alignment. Raw count matrix for 36,905 genes (including protein-coding and lncRNAs) across 12 samples was generated using featureCounts (subread v1.6.2) and analyzed in R v4.3.1 using DESeq2 v1.42.1. The spike-in normalization method was replicated as specified by the original authors (applying estimateSizeFactors (dds, controlGenes = ERCC)).

CUT&Tag data were re-processed from FASTQ files associated with GEO accession GSE247071 (Liu et al., Neuron 2024). Reads were aligned with bwa mem (v0.7.15) to a combined reference comprising the hg38 main chromosomes and the E. coli K-12 MG1655 genome; alignments were filtered with samtools (v1.16.1) view -q 10 -F 4 (MAPQ ≥ 10, mapped reads, all contigs retained), then sorted and indexed. PCR duplicates were quantified but retained, as duplication rates were high and non-uniform across genotypes and removal would bias between-genotype comparisons. A per-sample scale factor was calculated as 10,000 divided by the number of reads aligned to the E. coli contig, and applied to hg38-only reads via deepTools bamCoverage (bin size 50 bp; no CPM/RPKM/SPMR normalization, no blacklist) to generate per-replicate bigWig tracks; per-genotype mean tracks were produced with bigwigAverage. Fourteen replicate bigWig files were generated in total: GFP-tagged MECP2-WT, -R133C, -R168X, and -KO (3 biological replicates each), and -R270X (2 biological replicates). Because CUT&Tag signal reflects antibody accessibility, tagmentation efficiency, and relative variant expression rather than a stoichiometric measure of bound protein, occupancy values are interpreted as relative rather than absolute chromatin association.

Whole-genome bisulfite sequencing (WGBS) data were obtained from GEO accession GSE230715 (Liu et al., Neuron 2024), comprising three biological replicates of WT hESC-derived neurons. BigWig files were downloaded from the GEO FTP server as a single tar archive (GSE230715_RAW.tar, 269 MB). Upon extraction, replicate 1 (GSM8145251, 175 MB) was found to be a valid bigWig file reporting CG-context methylation fraction (values 0–1) at covered CpG sites genome-wide. Replicates 2 (GSM8145252) and 3 (GSM8145253) were found to have their first 344 bytes zeroed, corresponding precisely to the bigWig main header and zoom-level table, rendering them unreadable by pyBigWig.

### RNA-seq principal component analysis

The DESeq2 dataset was transformed using varianceStabilizingTransformation (blind = FALSE), which incorporates the dispersion estimates already fit under the ∼condition design rather than treating all samples as a single unstructured group. Principal component analysis was performed using DESeq2’s plotPCA function, which selects the 500 genes (from the 36,905) with the highest variance across samples from the variance-stabilized data and applies prcomp (centered, not scaled to unit variance) to this subset. The first two principal components and their associated percentage of variance explained were extracted (returnData = TRUE) and visualized with ggplot2 v3.5.1, with samples colored by genotype.

### CUT&Tag genomic analysis

#### Genomic region sets

##### CpG islands

CpG island coordinates were defined by the UCSC hg38 CpG island track (hg38_CpGislands.bed). Only standard chromosomes were retained (excluding chrM and unlocalized/unplaced contigs). Three region sets for CGIs were used. All 27,949 identified CGIs were used for the CGI profile plot (Fig. 5c). For the ATAC-seq correlation plot (Ext. Fig. 6f), only the 26,995 CGIs overlapping with KO-CUT&Tag signal were used. For CGI occupancy distribution and odds ratio plots (Fig. 5e,f), a random sample of 3,000 CpG islands was drawn without replacement (NumPy seed = 42) to define the CGI analysis set.

##### Long gene bodies

Gene body coordinates were obtained from refGene.txt.gz (UCSC hg38). Unique transcript loci longer than 100 kb on standard chromosomes were retained, yielding 6,803 transcript intervals, which were interval-merged to collapse overlapping isoforms of the same gene into unique loci, resulting in 3,465 total long gene bodies in the annotation. The combined regions comprised 6,465 sites (3,000 CGIs + 3,465 long gene bodies), written to SuppS3_region_list.bed.

##### Signal extraction

Multiple signal-extraction approaches were used, depending on the analysis. Metagene profiles (CpG islands and long gene bodies) were generated with deepTools computeMatrix v3.5.6 (reference-point mode, centered on each region’s midpoint), extracting per-bin signal from each biological replicate (n = 3, except R270X n = 2) and averaging across replicates to yield a mean ± SEM profile per genotype. The CpG-island profile used the validated annotation (27,949 islands), a ±3 kb window, 50 bp bins (120 bins total), and missing data set to zero; no islands were excluded from this average. The long gene bodies profile used the refGene.txt annotation (3,465 regions; see Genomic region sets), a ±10 kb window, 100 bp bins (200 bins total), and left missing data undefined rather than zero-filling it, so that for each replicate independently, regions with no defined signal across the entire window (all bins undefined, or a maximum bin value ≤0) could be identified and excluded before that replicate’s profile was averaged across the remaining regions (99.4-99.7% of the 3,465 regions retained per replicate) ; a 7-point moving-average kernel was applied to this profile for display only. Tn5 background subtraction (see below) was applied identically to both metagene profiles.

The CpG-island midpoint bar chart extracted signal directly with pyBigWig v0.3.25 (bw.stats, ±250 bp window centered on the island midpoint, exact=True) from the same replicate tracks used for the CGI profile, and averaged across replicates per genotype. The distribution and odds-ratio analyses used a hybrid annotation of 3,000 CpG islands and 3,465 long gene bodies. Per-region mean signal was extracted directly with pyBigWig from each genotype’s spike-in normalized mean bigWig track, in which replicate averaging (deepTools bigwigAverage) had already been performed upstream.

The ATAC-seq vs. KO CUT&Tag correlation plot used the validated CpG-island annotation (27,949 islands), extracting one mean-signal value per island directly from each bigwig. Of these, 26,995 islands (96.5%) had valid (non-missing) signal in both tracks and were used for the Pearson/Spearman correlation, while the remainder were excluded due to missing data in one track.

##### Tn5 background subtraction

Because no IgG control track was included in GSE247071, MeCP2-KO neurons —which express no MeCP2 protein but have identical chromatin accessibility — were used as the Tn5 tagmentation background reference. KO CUT&Tag signal therefore represents untargeted Tn5 insertion driven by chromatin accessibility rather than antibody-directed binding.

##### Metagene profiles

For each genotype *g* at each bin position *i*:

corrected_meanₑ[i] = meanₑ[i] − mean_KO[i]

corrected_SEMₑ[i] = sqrt( SEMₑ[i]^2^ + SEM_KO[i]^2^)

The KO-subtracted KO line equals zero by definition and is shown as a dashed reference in the metagene plot.

##### CGI center signal (bar summary)

The mean Tn5-corrected signal at CGI centers was computed via pyBigWig bw.stats(type=’mean’, exact=True). The individual replicate tracks were used (not the averaged tracks), hence error and test reflect biological-replicate variation. Dunnett’s test (scipy.stats.dunnett) comparing each variant to the control was computed, with built-in multiple-comparison control.

##### Per-region Fisher test

For each individual region *i*, the KO-corrected signal was computed with a floor at zero to prevent negative values from inflating the region-level ranking:
corrected_e,i_ = max(signal_e,i_ – signal_KO,_i_,0)

Fisher exact test for CGI vs long gene bodies selectivity

For each genotype, Tn5-corrected signals across all 6,465 regions (CGI and long gene body combined) were pooled into a single ranked distribution. The top 25% of regions by corrected signal were designated “high-occupancy” (threshold = 75th percentile of the joint distribution). A 2×2 contingency table was constructed:

|  | High-occupancy | Low-occupancy |
| --- | --- | --- |
| CpG island | $a$ | $b$ |
| Long gene body | $c$ | $d$ |

A two-sided Fisher exact test was applied (scipy.stats.fisher_exact). The odds ratio (OR) and 95% confidence interval on log_2_(OR) were computed using the normal approximation:

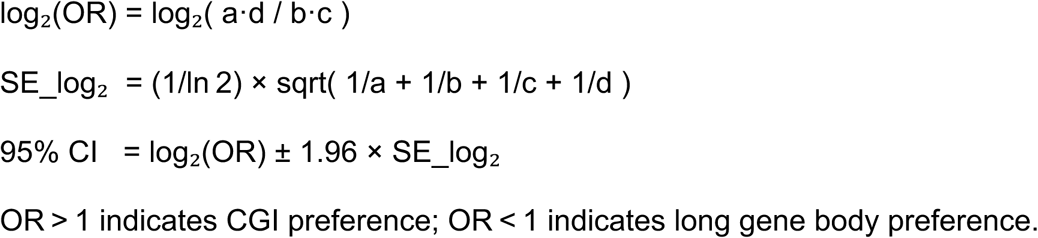

### WGBS methylation validation

To confirm the DNA methylation context of the two genomic feature classes used in the CUT&Tag analysis, mean CG methylation fraction was extracted from a WT WGBS bigWig (GSM8145251) at CpG island and long-gene-body intervals from the 3,000 CpG islands, 3,465 long gene bodies [≥100 kb] region set. Each interval was queried with pyBigWig using 5 bins per interval (bw.stats(type=’mean’, nBins=5)), and per-interval means were computed excluding bins with no covered CpGs (None/NaN). The difference between CpG islands and long gene bodies was assessed with a two-sided Mann–Whitney U test.

### Software and reproducibility

All analyses were implemented in Python 3.9+ with the following package versions: pandas 2.x, numpy 1.x, scikit-learn 1.x, scipy 1.x, matplotlib 3.x, pyBigWig 0.3.22. Analysis scripts will be made available at Kasinath Lab GitHub (https://github.com/KasinathLab/mecp2-EMSA-FP-fitting and https://github.com/KasinathLab/mecp2_reanalysis_scripts). WGBS signal extraction was performed on a single replicate (GSM8145251) due to file corruption of GSM8145252 and GSM8145253 in the GEO repository. All random operations used numpy seed 42 for reproducibility.

### Data availability

The cryoEM density maps and the atomic coordinates have been deposited to the Protein Data Bank. The accession number for the atomic model of one MeCP2 binding to a WT nucleosome is PDB ID 36WF. The corresponding density map is EMD-77909. The accession number for the atomic model of two MeCP2 binding to a WT nucleosome is PDB ID 36YN. The corresponding density map is EMD-77962. The accession number for the atomic model of MeCP2 binding to hmLinker nucleosome is PDB 37TE. The corresponding density map is EMD-78474.

## ACKNOWLEDGEMENTS

We thank E. Hartwick and S. Laursen (University of Colorado Boulder Biochemistry Krios Electron Microscopy Facility, RRID: SCR_019057) for cryo-EM data screening and maintenance of computational infrastructure, A. Herbst (University of Colorado Boulder Biochemistry Shared Instrument Pool Facility, RRID: R24OD033699-01) for training and access to shared instrument facilities, and T. Nahreini (University of Colorado Boulder Biochemistry Cell Culture Facility, RRID: SCR_018988). Cryo-electron microscopy data were collected at the Pacific Northwest Cryo-EM Center (PNCC), supported by NIH grant U24GM129547 and located at the Environmental Molecular Sciences Laboratory (EMSL), a DOE Office of Science User Facility sponsored by the Office of Biological and Environmental Research. We thank Danielle Guillen for help with MeCP2 purification, nucleosome binding assays, and footprinting. We thank the PNCC staff for assistance with microscope operation and data collection. We thank Dr. Karoline Luger for providing histone octamer as a gift for hmLinker nucleosome reconstitution.

## Author Information

### Authors and Affiliations

**Department of Biochemistry, University of Colorado Boulder, Boulder, CO, USA**

Liqi Yao, Jiarui Song, Iona L. Kelly & Vignesh Kasinath

**BioFrontiers Institute, University of Colorado Boulder, CO, USA**

Carolina V. Hincapie, Jiarui Song & Edward B. Chuong

**Howard Hughes Medical Institute, University of Colorado Boulder, CO, USA**

Jiarui Song

**Department of Molecular, Cellular, and Developmental Biology, University of Colorado Boulder, CO, USA**

Carolina V. Hincapie, Edward B. Chuong

### Contributions

L.Y. and V.K. conceived the study, analyzed data, and wrote the manuscript. L.Y. performed all the biochemical experiments, EM sample preparation, data collection, and data processing. I.L.K assisted with the molecular cloning of MeCP2 variant constructs.

J.S assisted with the preparation of streptavidin affinity grids and cryo-EM sample preparation. C.H., E.B.C., and V.K. reanalyzed RNA-seq and CUT&Tag data. L.Y and V.K wrote the manuscript. L.Y., C.H., E.B.C., and V.K. read and edited the manuscript.

### Corresponding authors

Correspondence to Vignesh Kasinath.

### Extended Data Fig

**Structures of MeCP2 bound to nucleosomes reveal distinct mechanisms of Rett syndrome mutations**

**Extended Data Fig. 1.**
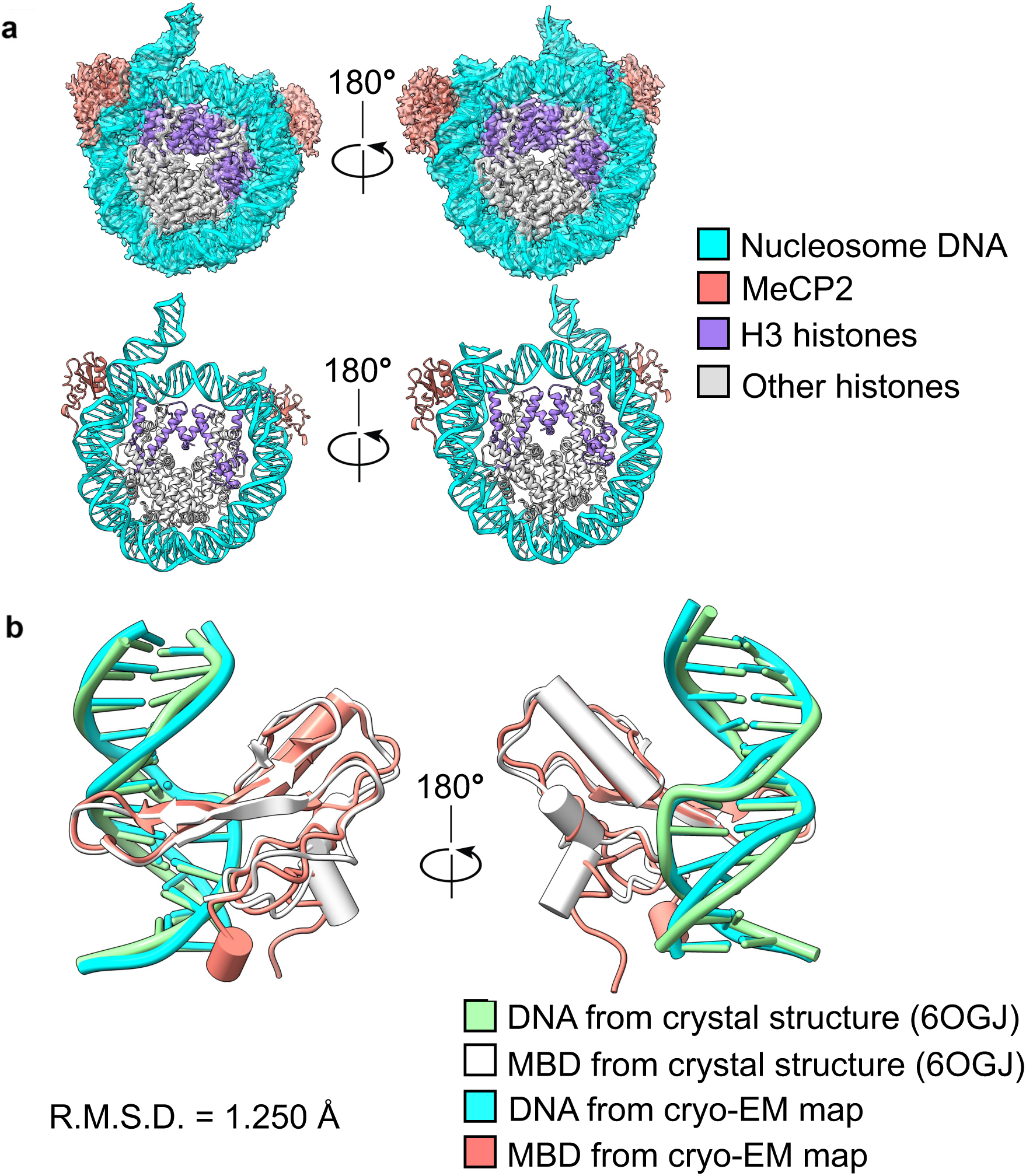
**(a)** Cryo-EM map and model of two MeCP2 (MBD) bound to a mono-nucleosome showing the MBD (salmon) bound at the SHL ± 7 location. Both MBDs bind to DNA in a similar geometry. **(b)** Structural comparisons of the MBD(white)-bound to DNA (green) between the crystal structure (PDB ID: 6OGJ) and the MBD (salmon) bound to nucleosomal DNA (cyan) from the cryo-EM structure of MeCP2-WT nucleosome in this study.

**Extended Data Fig. 2.**
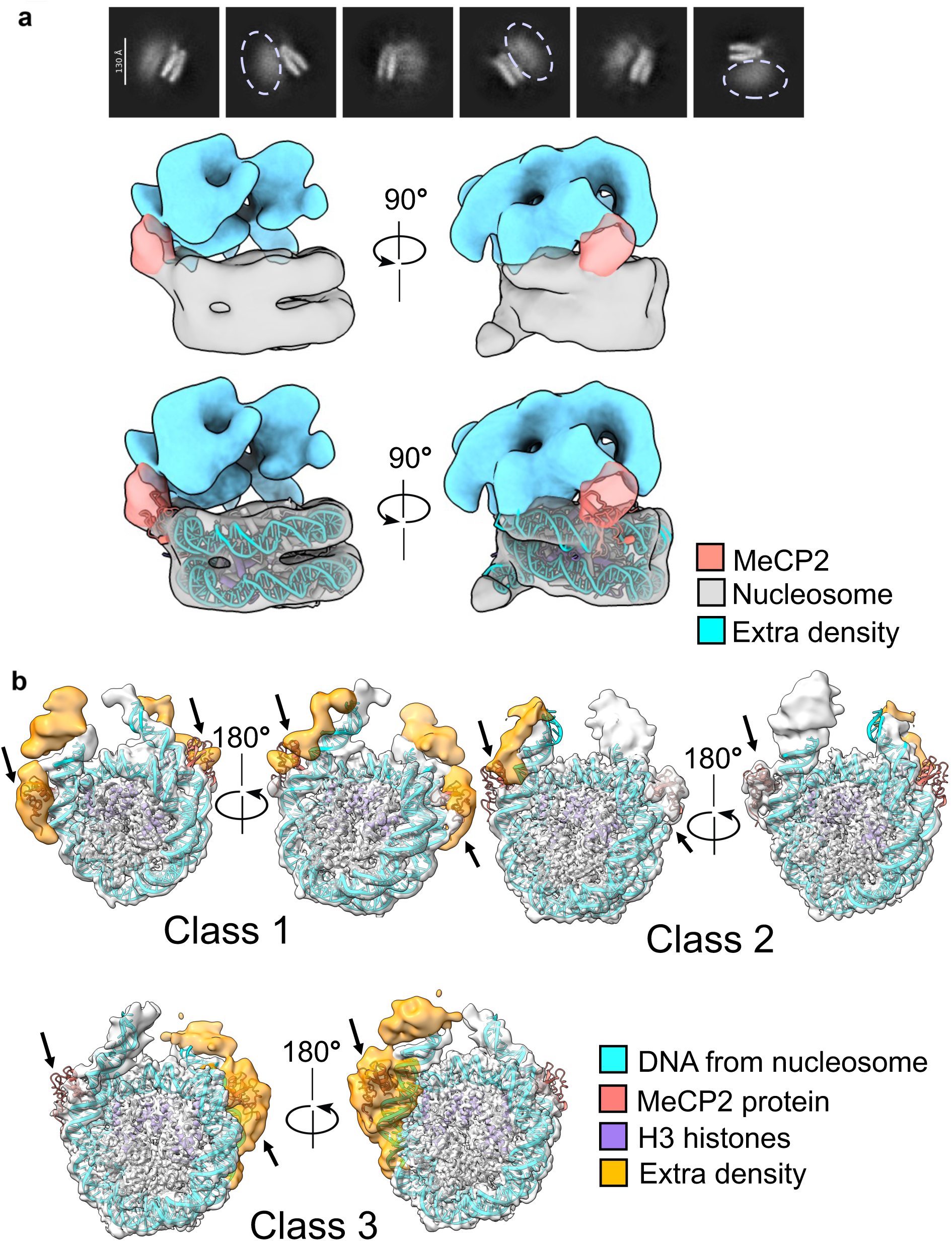
Analysis of conformational flexibility in cryo-EM reconstruction of MeCP2-nucleosome. **(a)** (top) Reference-free 2D classification reveals a subset of 2D class averages showing fuzzy density (dashed ellipse) above the nucleosome core. 3D refinement of the particles from these class averages revealed extra density (cyan) situated above the MBD (salmon) and nucleosome core (grey). Representative images of this reconstruction are shown at a low threshold (=0.08). **(b)** Non-uniform refinement and local resolution filtered cryo-EM maps of several distinct 3D classes obtained from 3D classification showing additional density (shown in yellow), proximal to the location of MBD (black arrow), on top of linker DNA. These additional densities were weak at high resolution and could not be modeled reliably.

**Extended Data Fig. 3.**
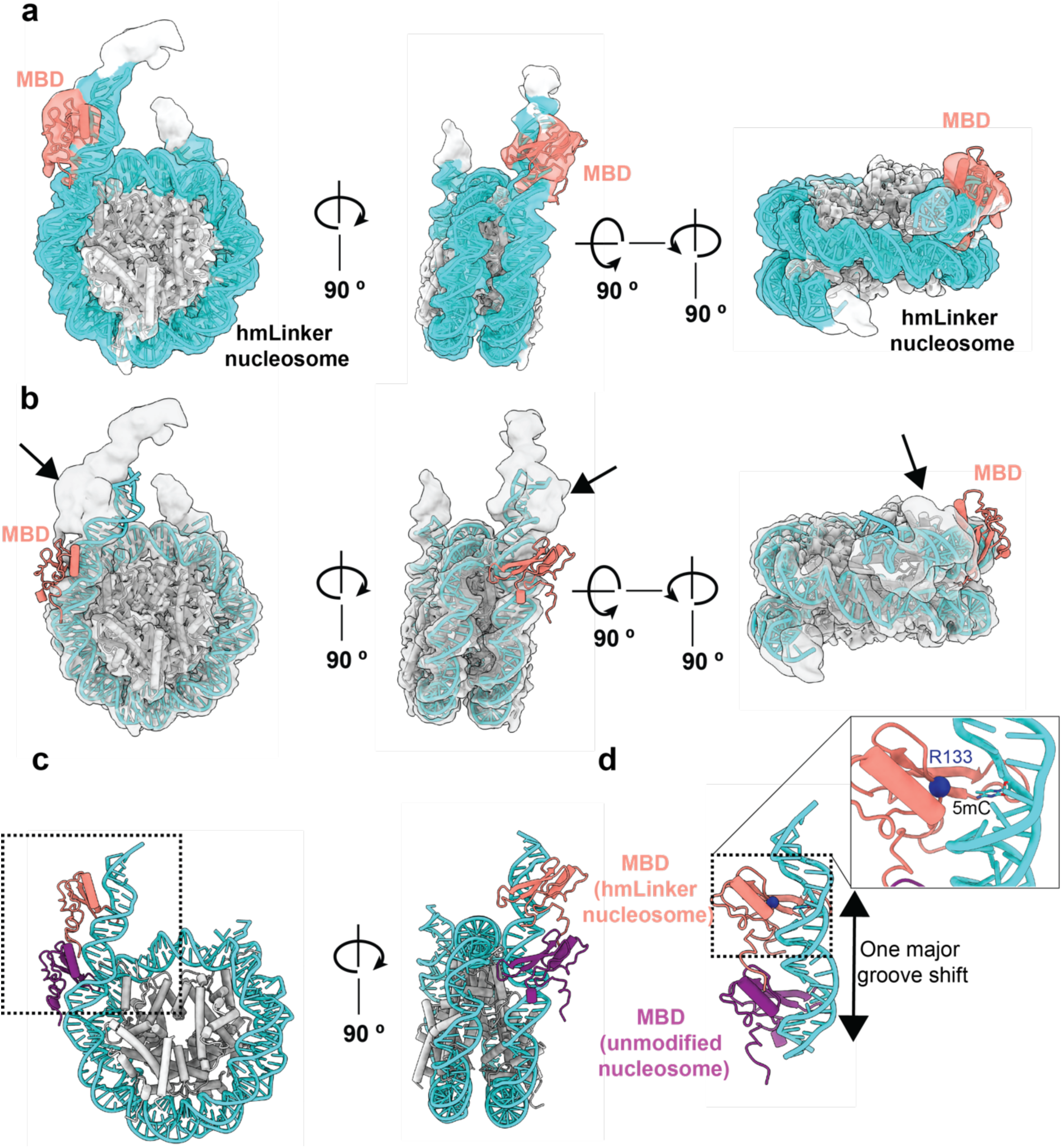
MeCP2 binds to the hmLinker nucleosome at the hemi-methylated DNA location. **(a)** Density map and model showing MeCP2 (salmon) binding to the hmLinker nucleosome (DNA: cyan, histones: gray). Only the MBD domain of MeCP2 was visible in the reconstruction. The model of the MeCP2(MBD)-hmLinker nucleosome was obtained by rigid-body docking the nucleosome and MBD domain, separately, into the cryo-EM reconstruction, followed by COOT and Phenix real-space refinement. **(b)** Representative view of the cryo-EM map of MeCP2 binding to the hmLinker nucleosome (light gray) with the model from the MeCP2-WT nucleosome rigid-body docked. The position of MBD is clearly different when bound to the hmLinker nucleosome. Black arrow marks the MBD density observed in the MeCP2-hmLinker nucleosome cryo-EM map. **(c)** Comparison of the MeCP2-hmLinker nucleosome model with the MeCP2-WT nucleosome model reveals the difference in MBD positioning on the nucleosomal DNA. The models were aligned with respect to the histone core. **(d)** A close look at the MeCP2-hmLinker nucleosome and MeCP2-WT nucleosome demonstrates that there is a major groove shift of the MeCP2(MBD) binding site on the nucleosomal DNA towards the linker region. The inset shows a closer view of the MBD binding orientation to the hmLinker nucleosomal DNA. The C*a* of Arg 133 in the MBD domain is shown as a blue sphere and is located 7.5 Å away from the 5mC of the DNA. Due to the low resolution of both the nucleosomal linker DNA and the MBD density, we cannot accurately model the side chain of R133, so we show these for illustrative purposes.

**Extended Data Fig. 4.**
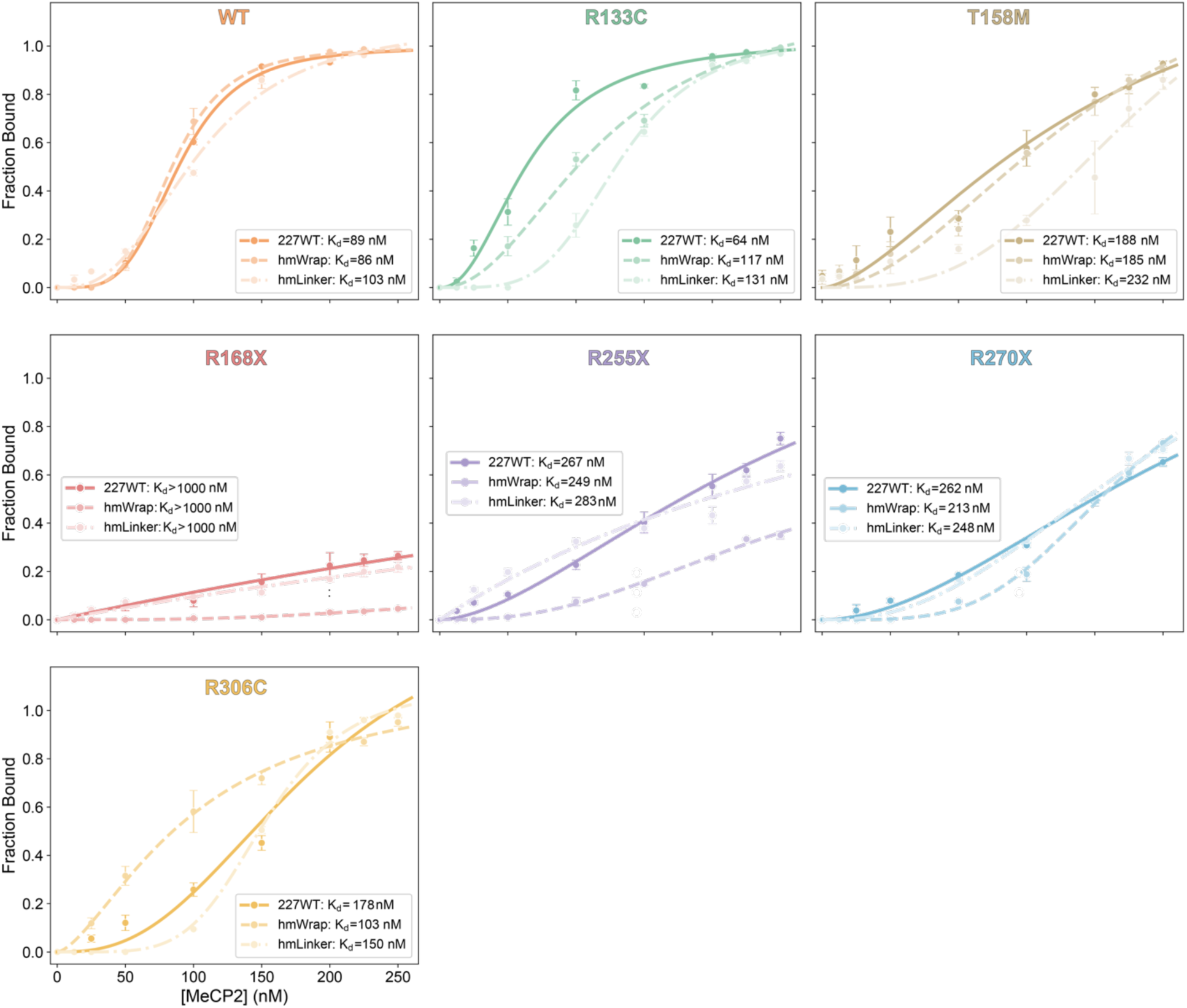
Binding isotherms for MeCP2 variants with the three nucleosomes. Individual binding curves (scatter plot; shown as points) and fit (solid or dashed lines) for all seven proteins characterized in this study (WT MeCP2 and six RTT variants: R133C, T158M, R168X, R255X, R270X, R306C) illustrate the differences in binding and cooperativity of each variant to the three different nucleosomes: unmodified (solid line), hmLinker (dot-dashed-dot), and hmWrap (dashed). The color scheme for each variant is the same as in Figures 3 and 4.

**Extended Data Fig. 5.**
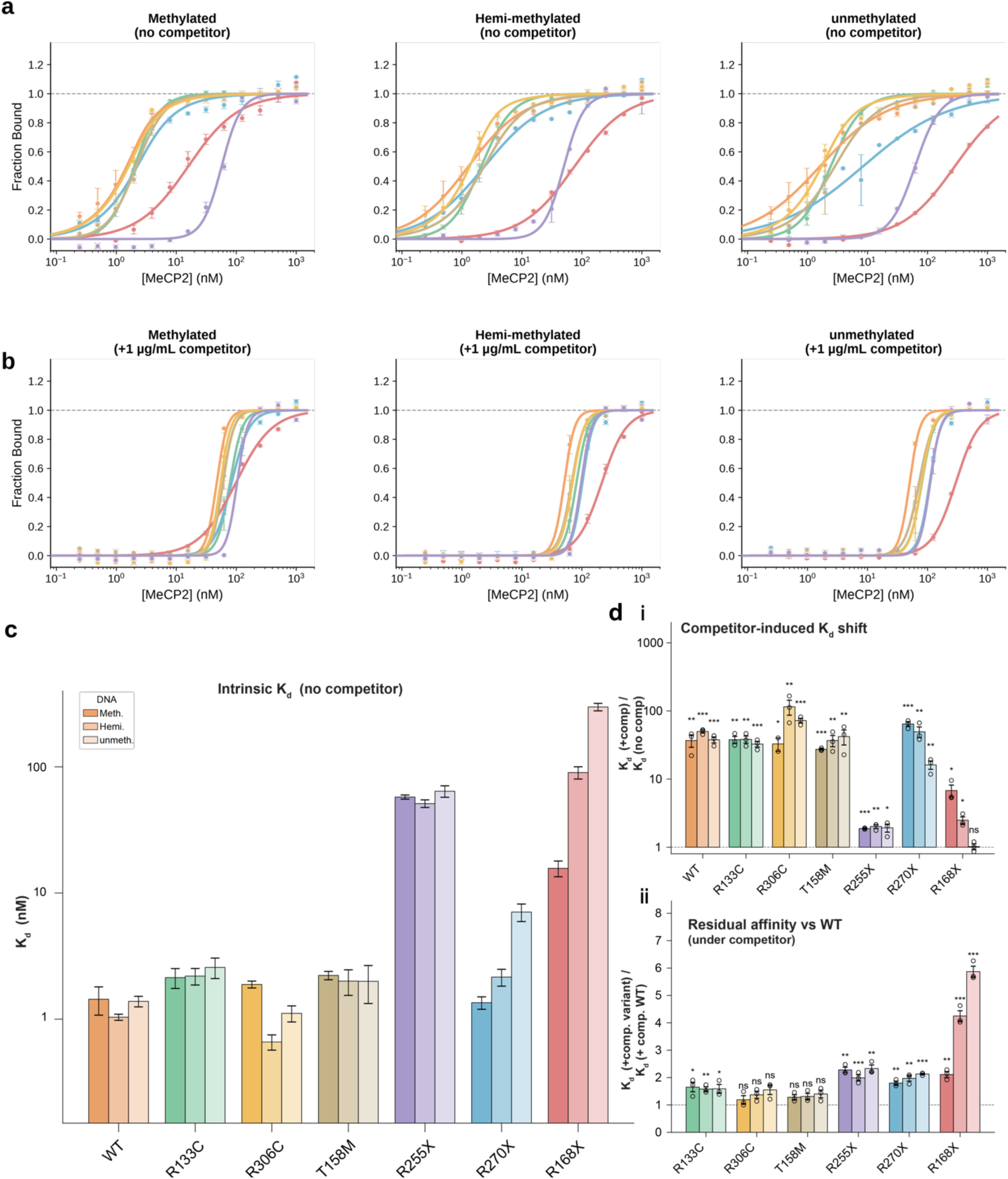
Fluorescence polarization reveals variant-specific DNA-binding affinities and altered competitor sensitivity. (**a, b**) Fluorescence polarization (FP) binding isotherms for WT MeCP2 and six Rett syndrome variants (R133C, R306C, T158M, R255X, R270X, R168X) in the absence and presence of 1 µg/mL poly(dI.dC) non-specific competitor, measured against a 15 bp FAM-labeled methylated (left), hemi-methylated (center), or unmethylated double-strand (right) DNA probe at 2 nM. Fraction bound is plotted on a linear scale against protein concentration (nM, log scale). Points represent individual replicates (n = 3), and lines show fits to the mean anisotropy trace. (c) Intrinsic DNA binding affinities (K_d_, no competitor) for each variant and DNA type. Three grouped bars per variant represent methylated (Meth.), hemi-methylated (Hemi), and unmethylated (Unmeth) DNA. Bars show the mean K_d_ from three independent replicate fits ± s.e.m. on a log scale. (d) Quantitative assessment of competitor sensitivity. (**d-i**) Fold change in K_d_ upon addition of competitor (FC1 = K_d_(+comp)/K_d_(no-comp)). Three grouped bars per variant represent methylated, hemi-methylated, and unmethylated DNA. Bars are mean ± s.e.m. and individual replicate values are overlaid as dots (n = 3 per bar). The statistical significance represented by * above each variant was calculated using a two-sided one-sample t-test. (**d-ii**) Residual affinity under competitor relative to WT (FC2 = K_d_(+comp, variant)/K_d_(+comp, WT)), shown as three grouped bars per variant for methylated, hemi-methylated, and unmethylated DNA. Bars are mean ± s.e.m. with replicate values overlaid as dots (n = 3 per bar). FC2 above 1 indicates weaker binding than WT under competitor conditions, reflecting the residual DNA binding that remains after the competitor-sensitive contacts are suppressed. The statistical significance above each variant represented by * was calculated using a two-sided Welch t-test. ns, not significant; *, p < 0.05; **, p < 0.01; ***, p < 0.001.

**Extended Data Fig 6.**
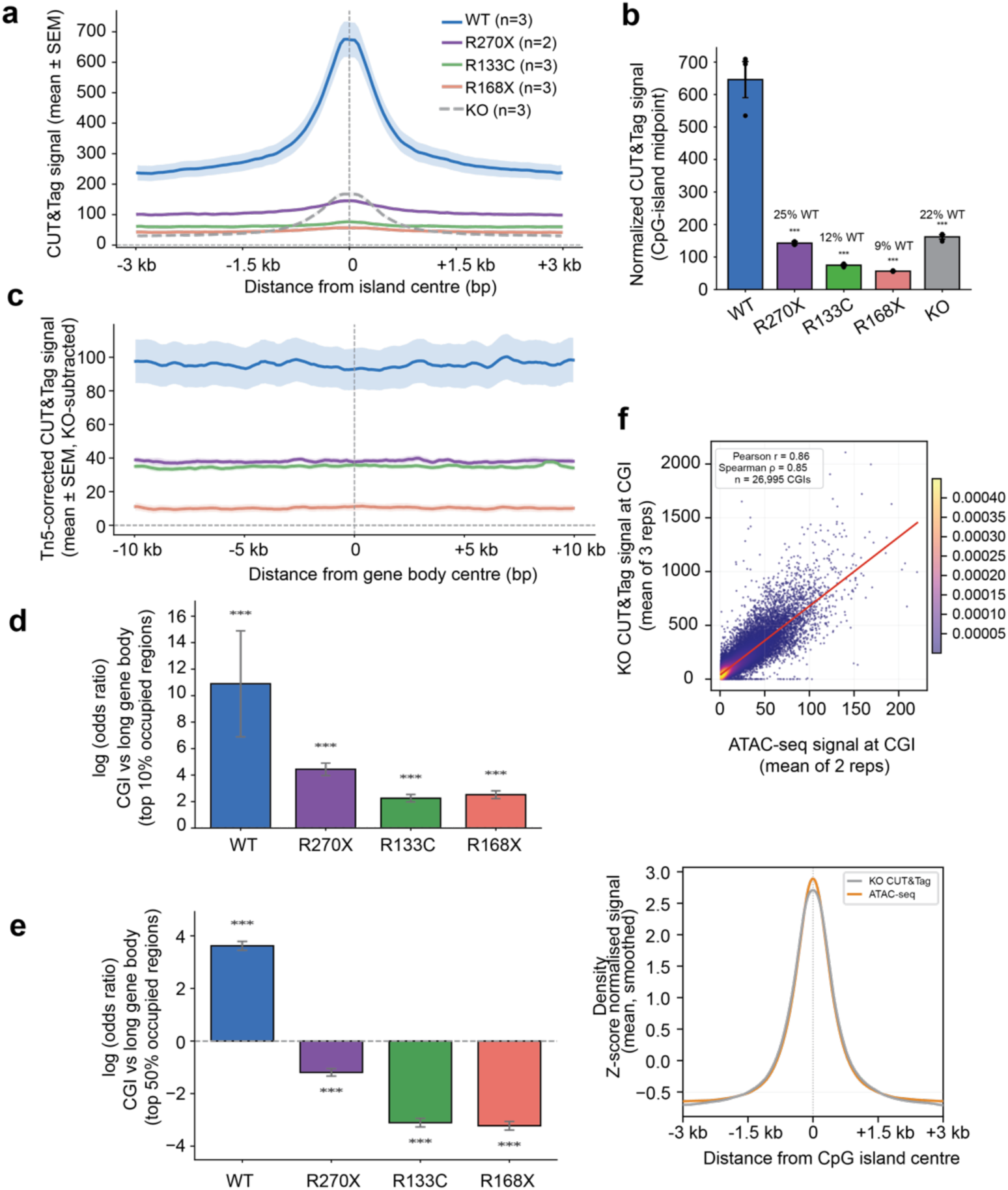
Validation of Tn5 background correction and robustness of MeCP2 chromatin occupancy redistribution in RTT variants. **(a)** Raw (uncorrected) CUT&Tag signal at CpG islands (±3 kb) for all five genotypes (WT -blue, R270X -purple, R133C -green, R168X -salmon, MeCP2-KO -gray), plotted as mean ± s.e.m. across three biological replicates. Prior to Tn5 background subtraction, KO neurons show substantial signal at CGI centers (dashed line, peak ∼ 167.4) reflecting untargeted Tn5 tagmentation driven by open chromatin. **(b)** Per-replicate CGI center signal for all five genotypes (raw, uncorrected; mean signal over ±500 bp from island midpoint across the 3,000-island CGI set). Each dot represents one biological replicate; the horizontal bar indicates the replicate mean. Tight clustering within genotype for RTT variants confirms that the low and negative Tn5-corrected values observed in the main analysis reflect genuine depletion rather than replicate-level outliers. WT replicate 3 shows a slightly elevated signal, reflected in the larger SEM visible in the main Fig. panel c bar graph. **(c)** Tn5-corrected (KO-subtracted) CUT&Tag signal at long gene body (±10 kb from the midpoint of each region; 200 bins, ∼100 bp/bin; isoform-deduplicated long gene body dataset, n = 3,465 regions, with regions lacking coverage excluded per replicate), plotted as mean ± s.e.m. The dashed line at zero represents the KO background level. In contrast to CpG islands, all four genotypes show positive corrected signal at gene body interiors (signal above Tn5 background), with the graded ordering WT > R270X > R133C > R168X. Critically, RTT variants that fall below the KO background at CGI centers retain detectable binding above background at gene body loci, consistent with redistribution of residual chromatin occupancy away from CGI-associated sites. **(d)** Fisher exact test log_2_ (OR) for CGI versus long gene body selectivity using a top-10% high-occupancy threshold (i.e., the top 10% of all 6,465 Tn5-corrected region signals). Error bars represent 95% confidence intervals computed by normal approximation on log_2_ (OR). At this stringent threshold, all four genotypes show positive log_2_ (OR) (CGI preference): WT = 10.89, R270X = 4.44, R168X = 2.52, R133C = 2.25. WT preference (OR = 1901.8, inflated by a zero-cell continuity correction) exceeded RTT variants (OR = 4.8-21.7) by roughly 90-to 400-fold, indicating that even among the very highest-signal regions, WT binding is overwhelmingly CGI-concentrated, while RTT variants showed modestly attenuated preference to CGI relative to WT. **(e)** Fisher exact test log_2_ (OR) at a top-50% threshold, corresponding to regions above the median corrected signal. OR=1 is shown as a dashed gray line. At this permissive threshold, WT retains strong CGI preference (OR = 12.29), whereas all three RTT variants shift to significant long gene body preference (OR = 0.12-0.44, all below 1), demonstrating that the bulk of RTT variant occupancy is non-CGI-associated. For this analysis, raw KO-subtracted values (without a floor at zero) were used for ranking to avoid degenerate contingency tables that would arise when many regions collapse to zero under a lenient threshold. **(f)** (top) Scatter of mean ATAC-seq signal (x) vs. mean MECP2-KO CUT&Tag signal (y) at CGIs, one point per island, colored by local point density. ATAC-seq data are from WT neurons (GSM8144406, GSM8144407; two biological replicates, averaged); KO CUT&Tag signal is averaged across three biological replicates. Signal for each island is the mean track value over that island’s own annotated interval (of 27,949 total CGIs in our used annotation, 26,995 with a valid signal in both tracks were used). The strong positive correlation (r = 0.86, ρ = 0.848, p < 0.0001) indicates that KO CUT&Tag signal closely tracks independently measured chromatin accessibility, validating its use as a background-subtraction reference for isolating MeCP2-specific occupancy in the Tn5-correction pipeline. (bottom) Metagene overlay of KO CUT&Tag and ATAC-seq across the same CGIs, ±3 kb from the island center (200 bins), both z-score normalized to allow direct shape comparison despite their different absolute scales. The similar shape and peak position of the two profiles further support the use of KO CUT&Tag as a proxy for chromatin accessibility at these loci.

**Extended Data Fig. 7.**
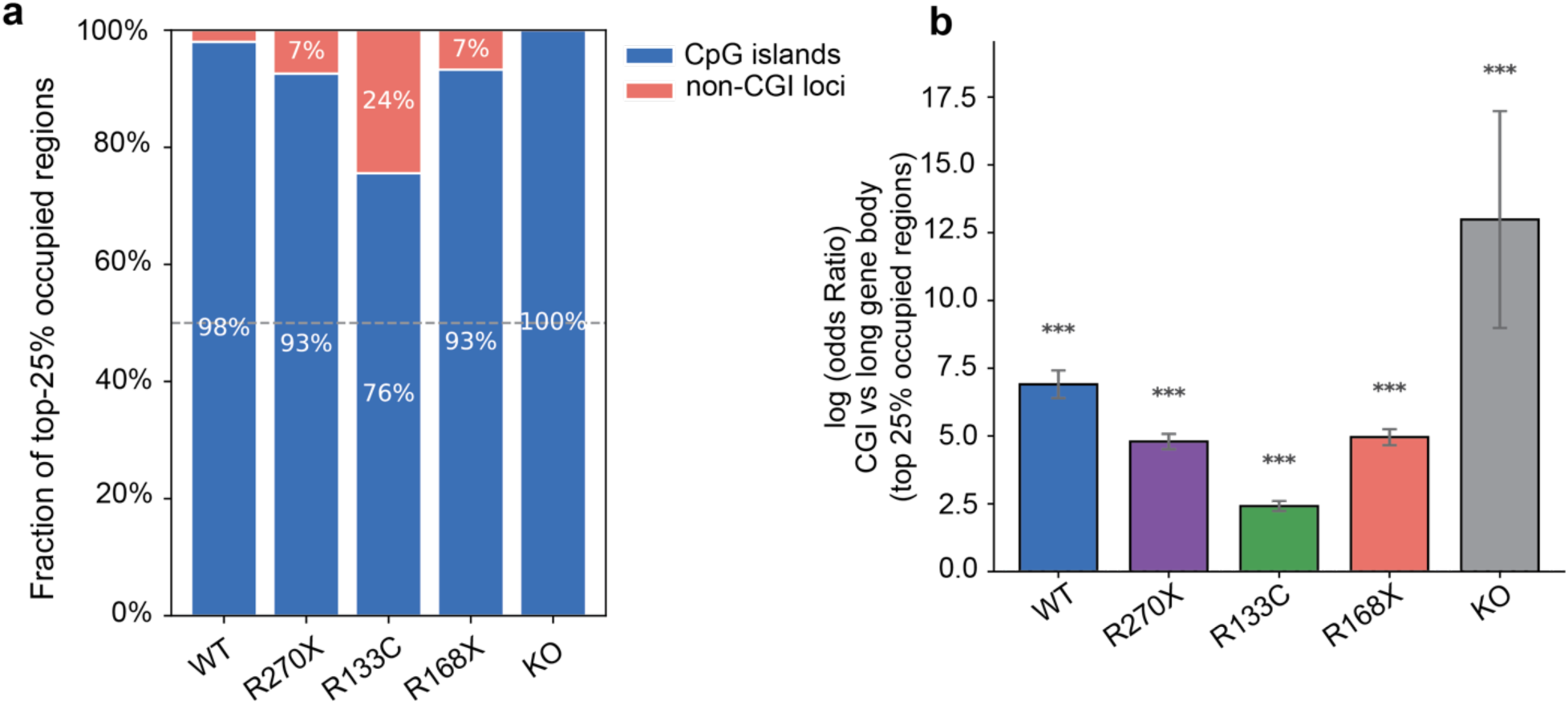
Reanalysis of MeCP2 chromatin occupancy from uncorrected CUT&Tag signal. **(a)** Stacked bar charts showing the fraction of each genotype’s top-25% most-occupied genomic regions (by CUT&Tag signal) that fall within CpG islands (blue) versus long gene bodies >100 kb (red), within a fixed Fisher exact set of 6,465 regions (3,000 CGIs + 3,465 long gene body). Dashed line at 50% indicates equal representation. **(b)** Fisher exact log odds ratios (log OR ± 95% CI) quantifying CGI over-representation among the top 25% most-occupied regions for each genotype. Higher values indicate stronger preferential occupancy at CpG islands relative to long gene bodies. *** p < 0.001 (two-sided Fisher exact test). R133C still shows the least CGI selectivity.

**Extended Data Fig. 8.**
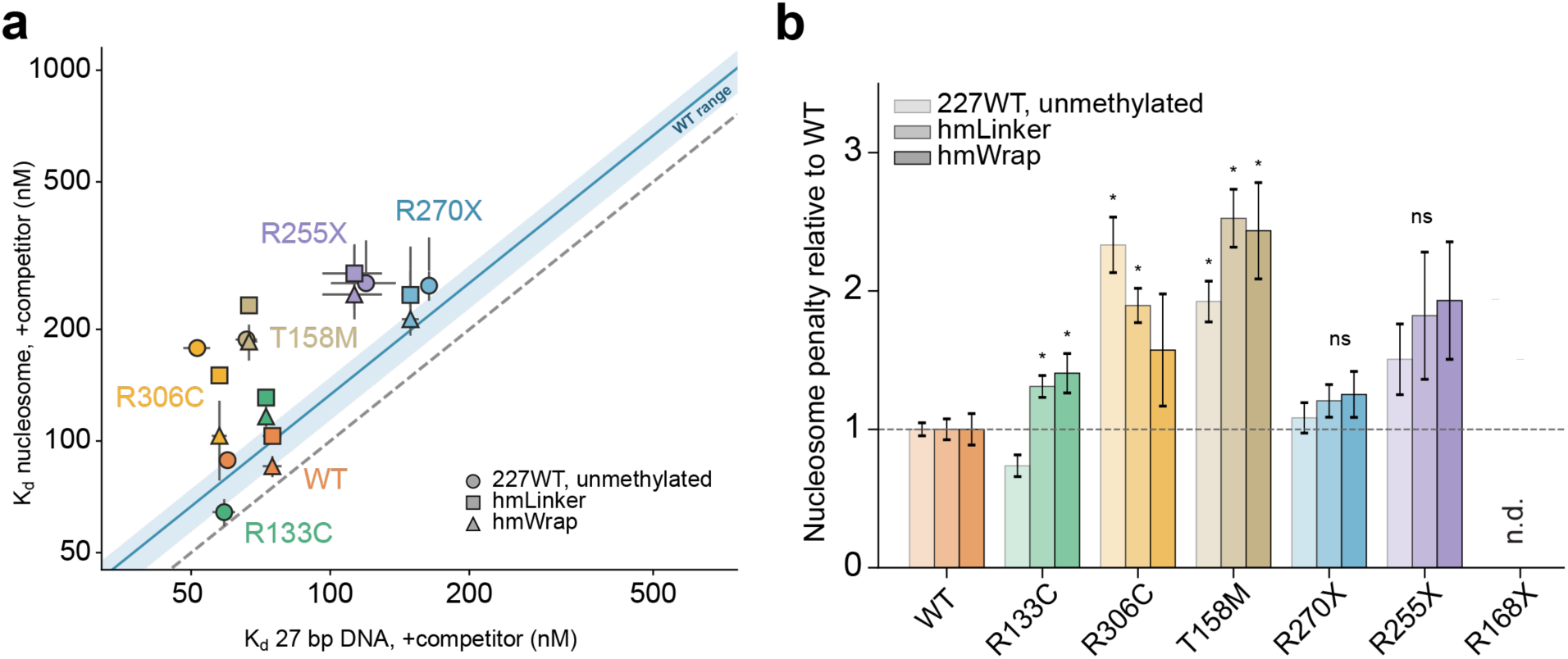
MeCP2 variants differ in how strongly they discriminate against the nucleosome relative to DNA. **(a)** Apparent K_d_ on each nucleosome plotted against apparent K_d_ on 27 bp DNA of matched methylation state with poly(dI.dC) competitor. The unmethylated 227WT (40-N-40) nucleosome is paired with unmethylated DNA, and hmLinker (23-N-40) and hmWrap (14-N-40) with hemi-methylated DNA. Dashed line, equal affinity on both substrates, meaning no discrimination between nucleosome and DNA. Shaded band, the range of discrimination measured for WT, 1.14 to 1.48-fold. Points above the band discriminate against the nucleosome more strongly than WT. **(b)** Nucleosome discrimination normalized to WT on the same substrate. Bars are the mean of n = 3 independent replicates, and error bars are propagated SEM. Asterisk represents p < 0.05 by Welch t-test. n.d., not determined. R133C shows a lower nucleosome penalty than WT on the unmodified nucleosome at 0.74-fold, p = 0.057.

