## Supplementary Information for "Structures of MeCP2 bound to nucleosomes reveal distinct mechanisms of Rett syndrome mutations"

**Primers and DNA sequences**

Primers and DNA sequence for FP:

| | |
|---|---|
| MeCP2_R133C_F | TAAAGCGTTTTGCAGCAAAGTGG |
| MeCP2_R133C_R | CCTTGCGGGTTGATC |
| MeCP2_R306C_F | GATCAAAAAGTGCAAGACCCGCG |
| MeCP2_R306C_R | GGCAGAACGGTTTCTTGG |
| MeCP2 R168X_F | TTAATAACATTGGAAGTGGATAACGG |
| MeCP2_R168X_R | CACGACGTGACGGGC |
| MeCP2_R270X_F | TAATTAATAACATTGGAAGTGGATAAC |
| MeCP2_R270X_R | ACGACCACGCTTTTTC |
| MeCP2_R255X_F | TAACTTAATAACATTGGAAGTGGATAACGGATCCG |
| MeCP2_R255X_R | GCGCTTGCGACCCGG |
| 27bpFP_F | FAM-GACTGTGTGCCCGCGATAATGGATAAT |
| 27bpFP_methylated_F | FAM-GACTGTGTGCCCG/5mC/GATAATGGATAAT |
| 27bpFP_R | ATTATCCATTATCGCGGGCACACAGTC |
| 27bpFP_methylated_R | ATTATCCATTAT/5mC/GCGGGCACACAGTC |
| 15bpFP_F | FAM-TGCCAGCGATAATGG |
| 15bpFP_methylated_F | FAM-TGCCAG/5mC/GATAATGG |
| 15bpFP_R | CCATTATCGCTGGCA |
| 15bpFP_methylated_R | CCATTAT/5mC/GCTGGCA |
| WTnucleosome_F | TACGTACGACGACGACGTACGTA |
| WTnucleosome_R | BioTEG – GCATCGATCGGGAGCTCC |
| Linker_nucleosome_F | CACGCGACTGTGTGCCCG/5mC/G |
| Linker_nucleosome_R | BioTEG – GCATCGATCGGGAGCTCC |
| New Linker_nuc_F | CACGCGA/5mC/GATATTGTCG |
| NewLinker_nuc_R | BioTEG – GCATCGATCGGGAGCTCC |
| Wrap_nucleosome_F | CACGCGACTGTGTGCTGGA/5mC/G |
| Wrap_nucleosome_R | BioTEG – GCATCGATCGGGAGCTCC |

**Nucleosome DNA and MeCP2 sequences**

(The underlined sequence is the 601-Widom sequence. The bold letters indicate the mutations from G-to-T or C-to-T to eliminate CA/GT sequence at SHL+-7.)

WT nucleosome
CACGCGACTGTGTGCCCGTCAGACGCTGCGCTGCCGGCGGCTGGAGAATCCCGG
TGCCGAGGCCGGCTCAATTGGTCGTAGACAGCTCTAGCACCGCTTAAACGCACGT
ACGCGCTGTCCCCCGCGTTTTAACCGCCAAGGGGATTACTCCCTAGTCTCCAGGC
ACGTGTCAGATATATACATCCTGTATGCATGCATATCATTCGATCGGAGCTCCCGA
TCGATGC

hmLinker nucleosome

CACGCGACTGTGTGCCCG/5mC/GATAATGGA**T**AATCCCGGTGCCGAGGCCGGCT
CAATTGGTCGTAGACAGCTCTAGCACCGCTTAAACGCACGTACGCGCTGTCCCCC
GCGTTTTAACCGCCAAGGGGATTACTCCCTAGTCTCCAGGCACGTGTCAGATATAT
A**T**ATCCTGTATGCATGCATATCATTCGATCGGAGCTCCCGATCGATGC

New_hmLinker nucleosome for structural study
CACGCGA/5mC/GATATTGTCGGTGGCTGGA**T**AATCCCGGTGCCGAGGCCGGCTCA
ATTGGTCGTAGACAGCTCTAGCACCGCTTAAACGCACGTACGCGCTGTCCCCCGC
GTTTTAACCGCCAAGGGGATTACTCCCTAGTCTCCAGGCACGTGTCAGATATATAT
A**T**CCTGTATGCATGCATATCATTCGATCGGAGCTCCCGATCGATGC

hmWrapped nucleosome
CACGCGACTGTGTGCTGGA/5mC/GAATCCCGGTCTGCAGGCCGCTCAATTGGTCG
TAGACAGCTCTAGCACCGCTTAAACGCACGTACGCGCTGTCCCCCGCGTTTTAAC
CGCCAAGGGGATTACTCCCTAGTCTCCAGGCACGTGTCAGATATATATA**T**CCTGTA
TGCATGCATATCATTCGATCGGAGCTCCCGATCGATGC

WT MeCP2 sequences
MVAGMLGLREEKSEDQDLQGLKDKPLKFKKVKKDKKEEKEGKHEPVQPSAHHSAEPA
EAGKAETSEGSGSAPAVPEASASPKQRRSIIRDRGPMYDDPTLPEGWTRKLKQRKSG
RSAGKYDVYLINPQGKAFRSKVELIAYFEKVGDTSLDPNDFDFTVTGRGSPSRREQKP
PKKPKSPKAPGTGRGRGRPKGSGTTRPKAATSEGVQVKRVLEKSPGKLLVKMPFQTS
PGGKAEGGGATTSTQVMVIKRPGRKRKAEADPQAIPKKRGRKPGSVVAAAAAEAKKK
AVKESSIRSVQETVLPIKKRKTRETVSIEVKEVVKPLLVSTLGEKSGKGLKTCKSPGRKS
KESSPKGRSSSASSPPKKEHHHHHHHSESPKAPVPLLPPLPPPPPEPESSEDPTSPPE
PQDLSSSVCKEEKMPRGGSLESDGCPKEPAKTQPAVATAATAAEKYKHRGEGERKDI
VSSSMPRPNREEPVDSRTPVTERVS

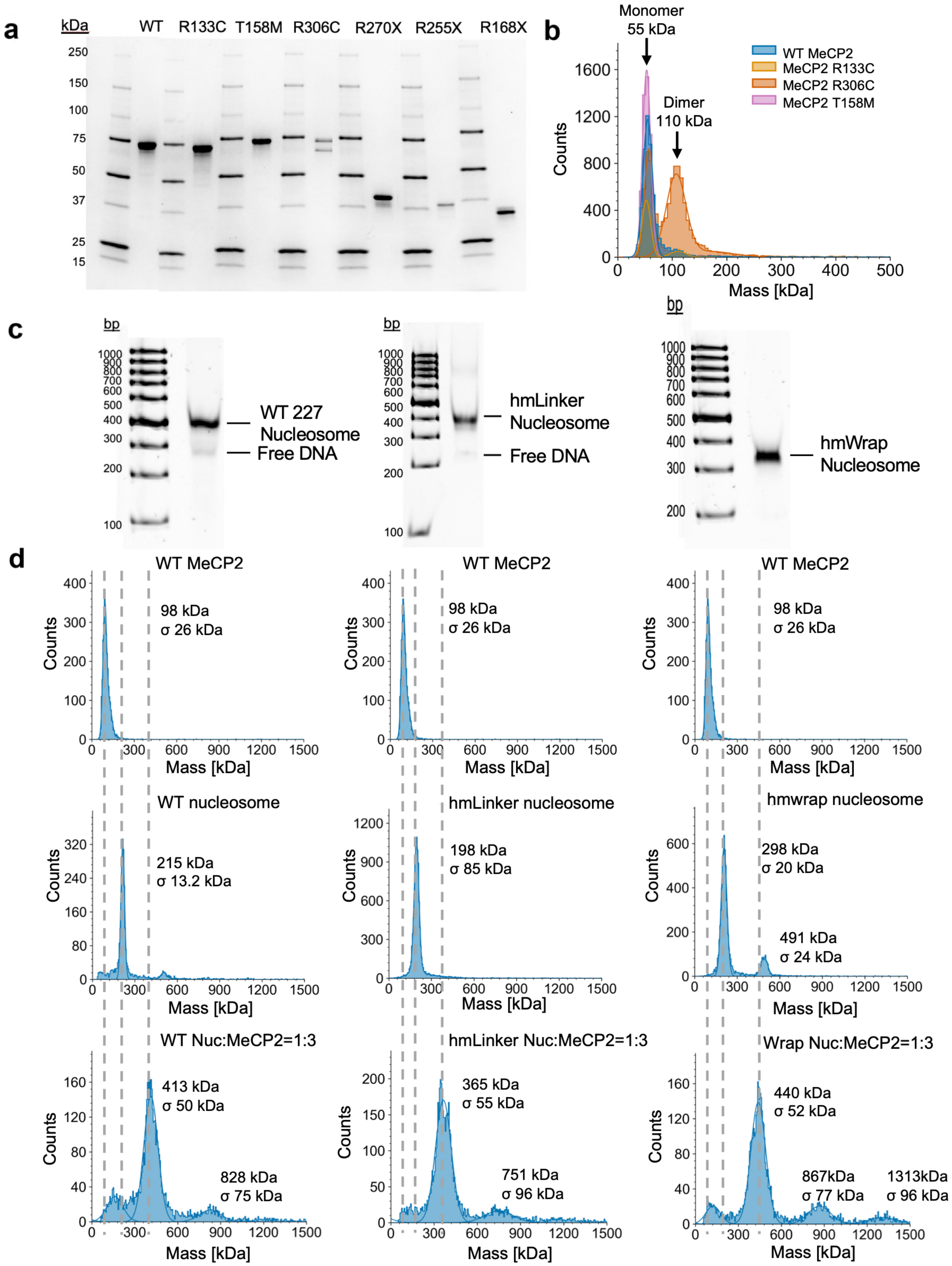


**Supplementary Figure 1: Validation of purified MeCP2, RTT variants, nucleosomes, and MeCP2-nucleosome complexes. (a)** Coomassie-stained SDS polyacrylamide gel of MeCP2, mutations, and truncations. **(b)** Mass photometry of MeCP2 and missense RTT variants. All samples are diluted to 20-50 nM in MP buffer 1 (25 mM HEPES, pH 7.5,

150 mM NaCl, 1 mM TECP). MeCP2 R306C shows both monomer (~55 kDa) and dimer peak (~110 kDa), whereas other protein constructs only show a monomer peak (~55 kDa). **(c)** 4% Native gel of WT, hmLinker, and hmWrap nucleosome. **(d)** Mass photometry of MeCP2, nucleosomes, and MeCP2 bound to nucleosomes with MeCP2 and nucleosomes at a 1:3 ratio. All samples are diluted to 20-50 nM in MP buffer 2 (25 mM HEPES, pH 7.5, 50 mM NaCl, 1 mM TECP). The MeCP2-only sample forms dimers at the current salt concentration. A clear shift in molecular weight indicates that MeCP2 binds to nucleosomes under the conditions we used for the structural study.

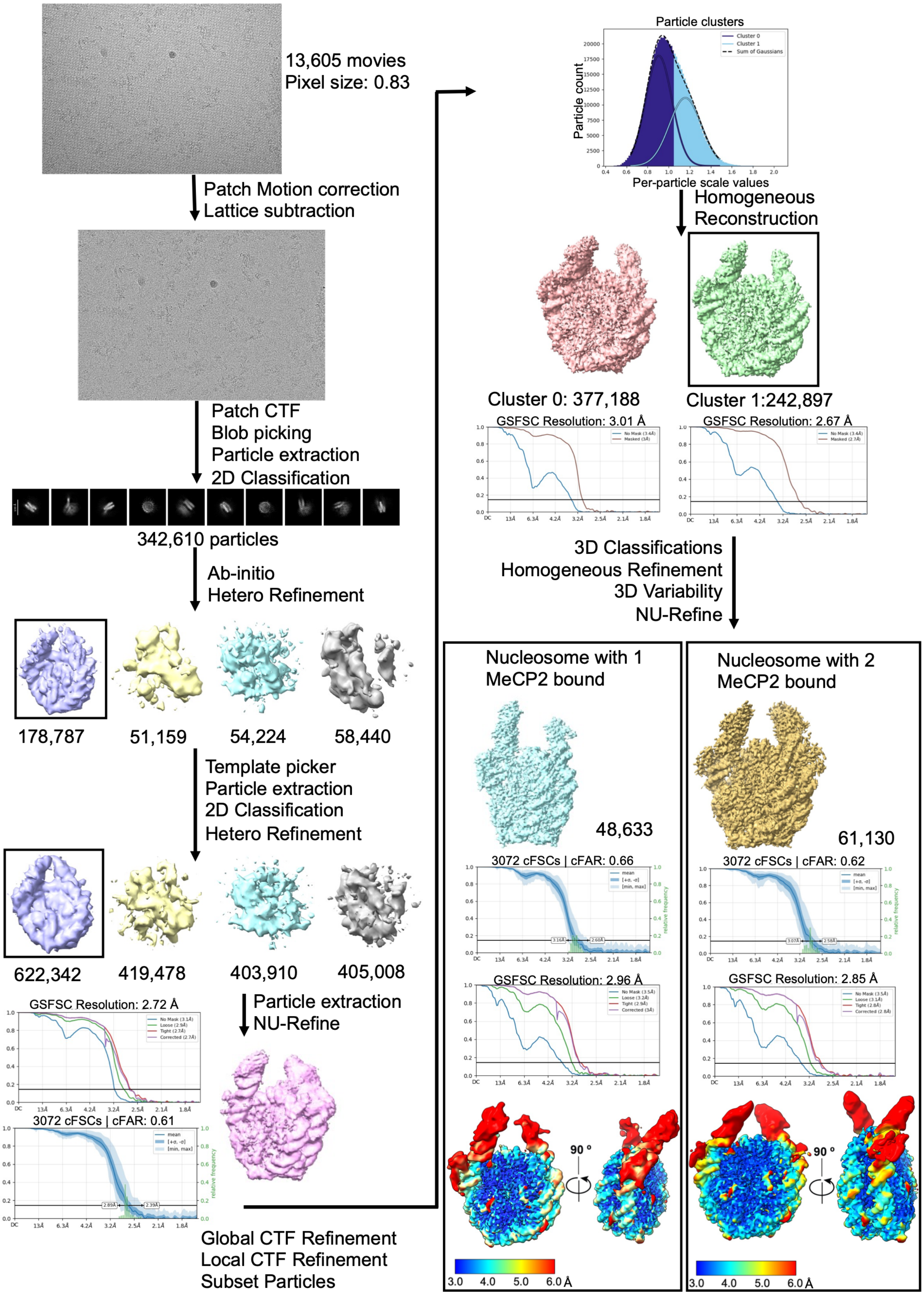
13,605 movies
Pixel size: 0.83
Patch Motion correction
Lattice subtraction
Patch CTF
Blob picking
Particle extraction
2D Classification
342,610 particles
Ab-initio
Hetero Refinement
178,787
51,159
54,224
58,440
Template picker
Particle extraction
2D Classification
Hetero Refinement
622,342
419,478
403,910
405,008
GSFSC Resolution: 2.72 Å
3072 cFSCs | cFAR: 0.61
Particle extraction
NU-Refine
Global CTF Refinement
Local CTF Refinement
Subset Particles
Particle clusters
Particle count
Per-particle scale values
Homogeneous
Reconstruction
Cluster 0: 377,188
Cluster 1:242,897
GSFSC Resolution: 3.01 Å
GSFSC Resolution: 2.67 Å
3D Classifications
Homogeneous Refinement
3D Variability
NU-Refine
Nucleosome with 1
MeCP2 bound
48,633
3072 cFSCs | cFAR: 0.66
GSFSC Resolution: 2.96 Å
Nucleosome with 2
MeCP2 bound
61,130
3072 cFSCs | cFAR: 0.62
GSFSC Resolution: 2.85 Å
90 °
3.0 4.0 5.0 6.0 Å

**Supplementary Figure 2.** Single-particle cryo-EM image processing workflow for MeCP2-WT nucleosome complex**.**

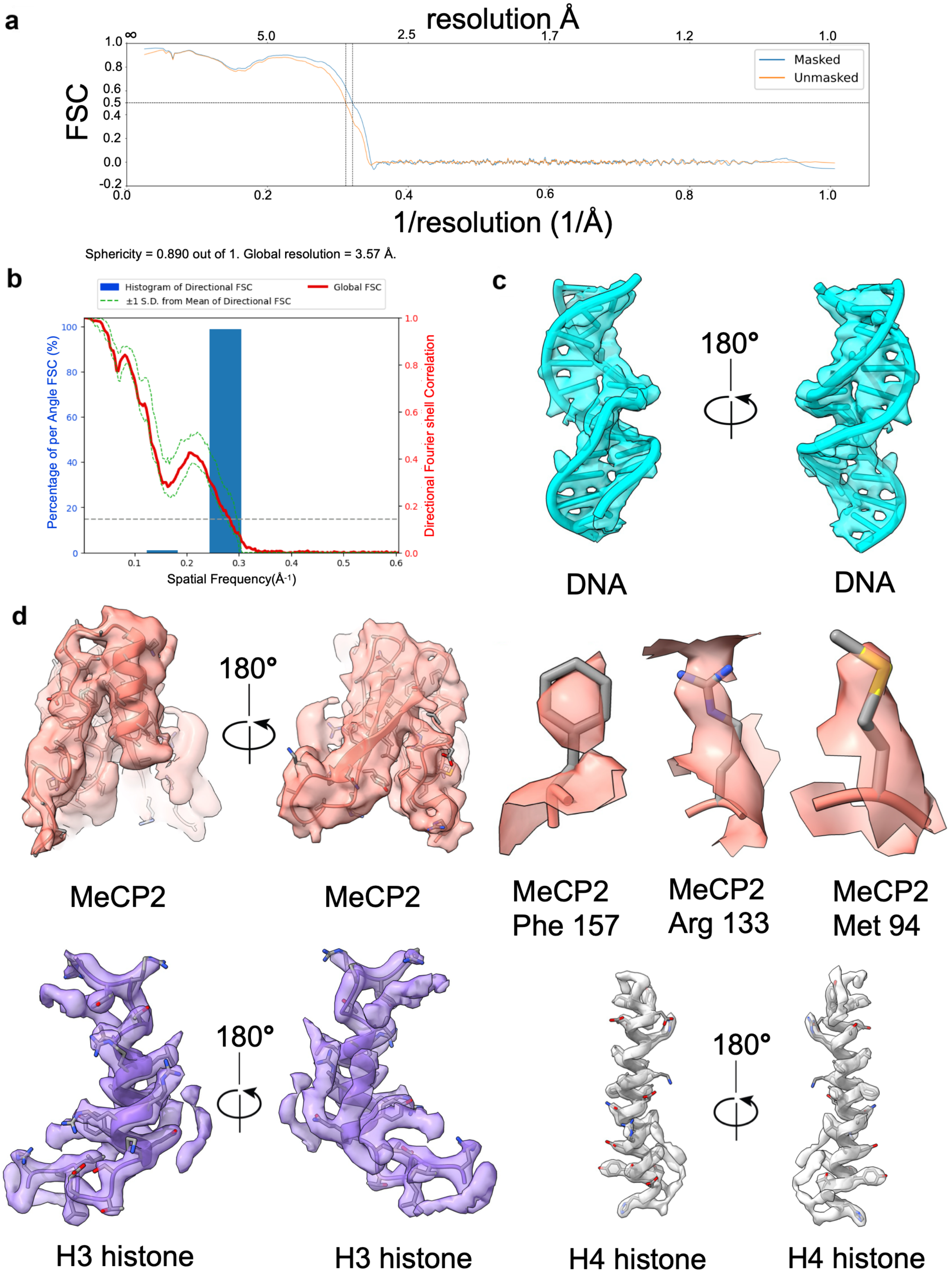
a
resolution Å
Masked
Unmasked
FSC
1/resolution (1/Å)
Sphericity = 0.890 out of 1. Global resolution = 3.57 Å.
b
Histogram of Directional FSC
Global FSC
±1 S.D. from Mean of Directional FSC
Percentage of per Angle FSC (%)
Directional Fourier shell Correlation
Spatial Frequency(Å-1)
c
180°
DNA
DNA
d
180°
MeCP2
MeCP2
MeCP2 Phe 157
MeCP2 Arg 133
MeCP2 Met 94
180°
H3 histone
H3 histone
180°
H4 histone
H4 histone

**Supplementary Figure 3: Validation of the MeCP2-WT nucleosome cryo-EM structural analysis. (a)** Model vs. map FSC for MeCP2-WT nucleosome. **(b)** Histogram and 3D FSC plot for MeCP2-WT nucleosome. **(c, d)** Examples of cryo-EM density from non-uniform refinement and built-in models for nucleosome DNA and different regions of MeCP2 and histone proteins.

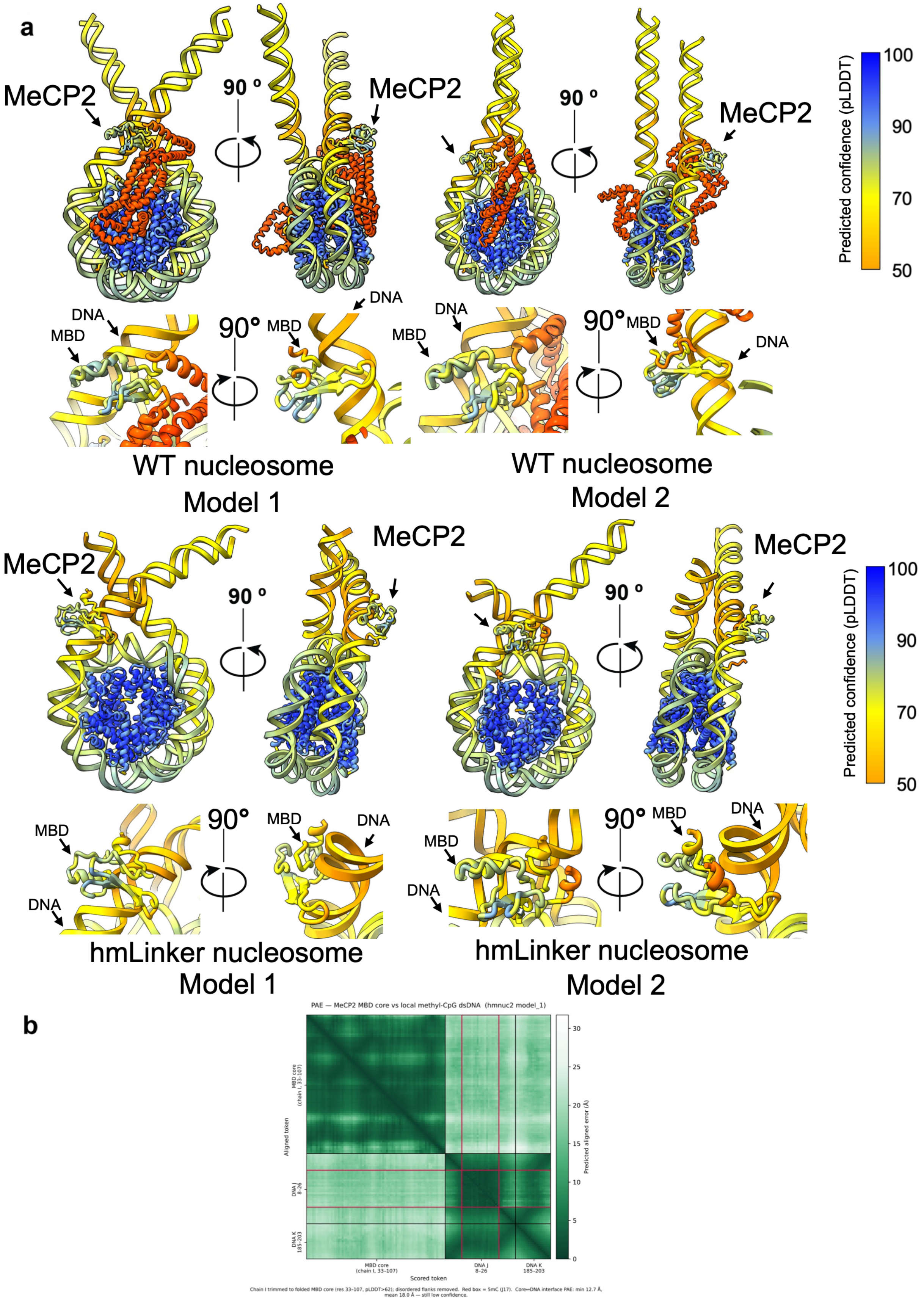
a
MeCP2
90 º
MeCP2
90 º
MeCP2
Predicted confidence (pLDDT)
100
90
70
50
DNA
MBD
90°
MBD
DNA
DNA
MBD
90°
MBD
DNA
WT nucleosome
Model 1
WT nucleosome
Model 2
MeCP2
90 º
MeCP2
90 º
MeCP2
Predicted confidence (pLDDT)
100
90
70
50
MBD
DNA
90°
MBD
DNA
MBD
DNA
90°
MBD
DNA
hmLinker nucleosome
Model 1
hmLinker nucleosome
Model 2
b
PAE — MeCP2 MBD core vs local methyl-CpG dsDNA (hmnuc2 model_1)
Aligned token
MBD core
(chain I, 33–107)
DNA J
8–26
DNA K
185–203
Scored token
Predicted aligned error (Å)
30
25
20
15
10
5
0
Chain I trimmed to folded MBD core (res 33–107, pLDDT>62); disordered flanks removed. Red box = 5mC (J17). Core↔DNA interface PAE: min 12.7 Å, mean 18.0 Å — still low confidence.

**Supplementary Figure 4: AlphaFold 3 prediction of MeCP2 binding to different nucleosomes. (a)** AlphaFold3 prediction of full-length MeCP2 binding to a mono-nucleosome. (top) Shown are the two best predicted models (iPTM=0.63; MeCP2 chain iPTM=0.17) and the close-up views of MeCP2 (MBD) binding to the nucleosome for the MeCP2-WT nucleosome. (bottom) Shown are the two best predicted models (iPTM=0.7; MeCP2(MBD) chain iPTM=0.4) and the close-up views of MeCP2 (MBD) binding to the nucleosome with a hemi-methylation site 15 bp from the entry/exit site of nucleosomal DNA. The predicted models show a different binding orientation of MeCP2 (MBD) relative to the nucleosome, likely due to poor predictive accuracy. The models are colored by their predicted Local Distance Difference Test (pLDDT) scores, and the position of MeCP2 (MBD) is marked with a black arrow. **(b)** A representative Predicted Aligned Error (PAE) plot for MeCP2 (MBD) binding to the hmLinker nucleosome from the AlphaFold 3 prediction clearly shows the high alignment error for the position of MBD near DNA. The PAE plot shown is restricted to aa 33-107 of MBD. The red box contains the 5mC in one strand of DNA (chain J; position 17), and the MBD+DNA interface PAE ranges from 12.7 Å (min) to 18 Å (mean), indicating low confidence.

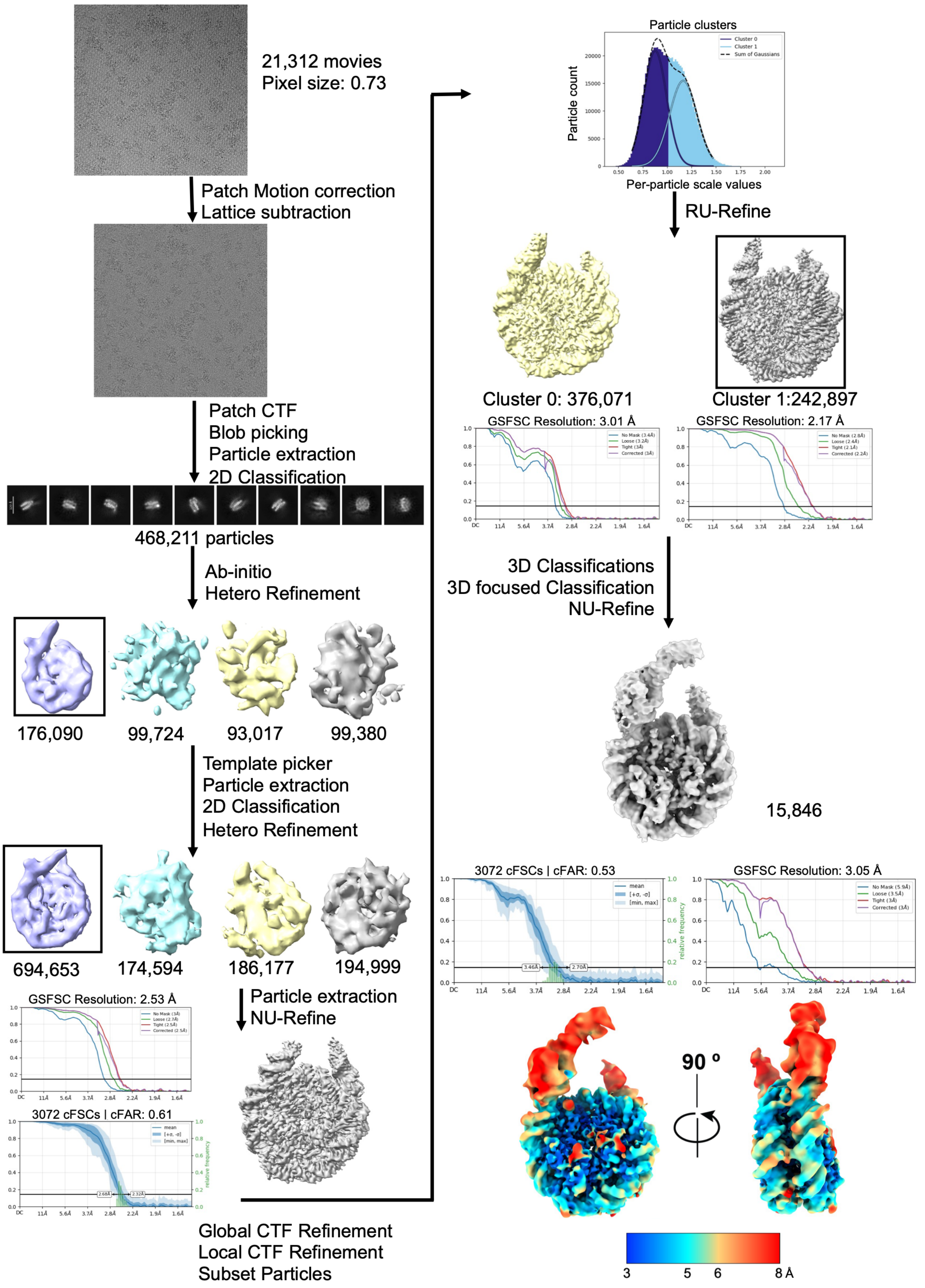
21,312 movies
Pixel size: 0.73
Patch Motion correction
Lattice subtraction
Patch CTF
Blob picking
Particle extraction
2D Classification
468,211 particles
Ab-initio
Hetero Refinement
176,090
99,724
93,017
99,380
Template picker
Particle extraction
2D Classification
Hetero Refinement
694,653
174,594
186,177
194,999
GSFSC Resolution: 2.53 Å
3072 cFSCs | cFAR: 0.61
Particle extraction
NU-Refine
Global CTF Refinement
Local CTF Refinement
Subset Particles
Particle clusters
Cluster 0
Cluster 1
Sum of Gaussians
Particle count
Per-particle scale values
RU-Refine
Cluster 0: 376,071
Cluster 1:242,897
GSFSC Resolution: 3.01 Å
GSFSC Resolution: 2.17 Å
3D Classifications
3D focused Classification
NU-Refine
15,846
3072 cFSCs | cFAR: 0.53
GSFSC Resolution: 3.05 Å
90 º
3
5
6
8 Å

**Supplementary Figure 5.** Single-particle cryo-EM image processing workflow for MeCP2-hmLinker nucleosome complex**.**

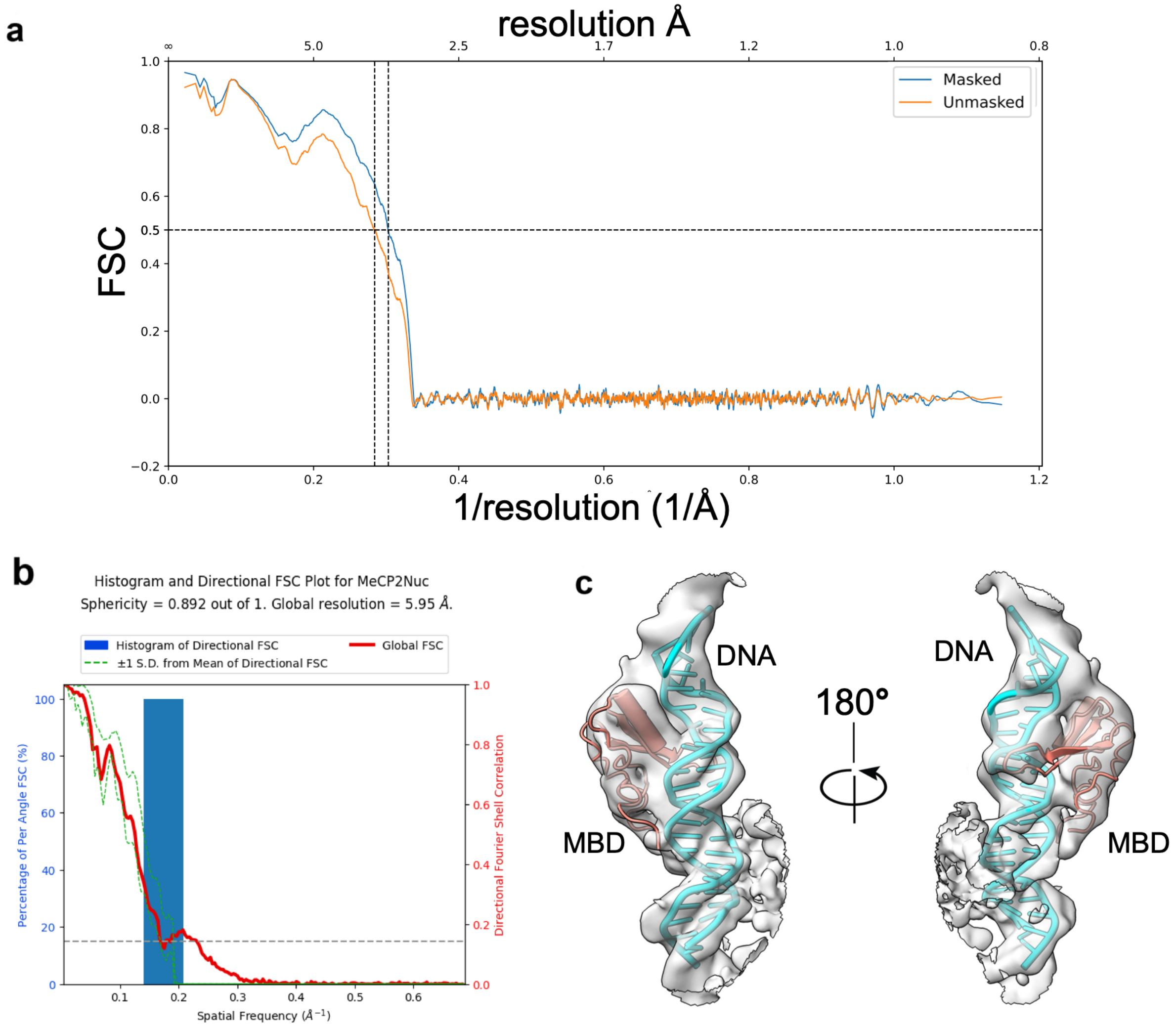


**Supplementary Figure 6: Validation of the MeCP2-hmLinker nucleosome cryo-EM structural analysis. (a)** Model vs. map FSC for the MeCP2-hmLinker nucleosome. **(b)** Histogram and 3D FSC plot for MeCP2-hmLinker nucleosome. **(c)** Close-up view of the cryo-EM density from the locally resolution-filtered map and docked model of MBD with the built-in model for nucleosome DNA.

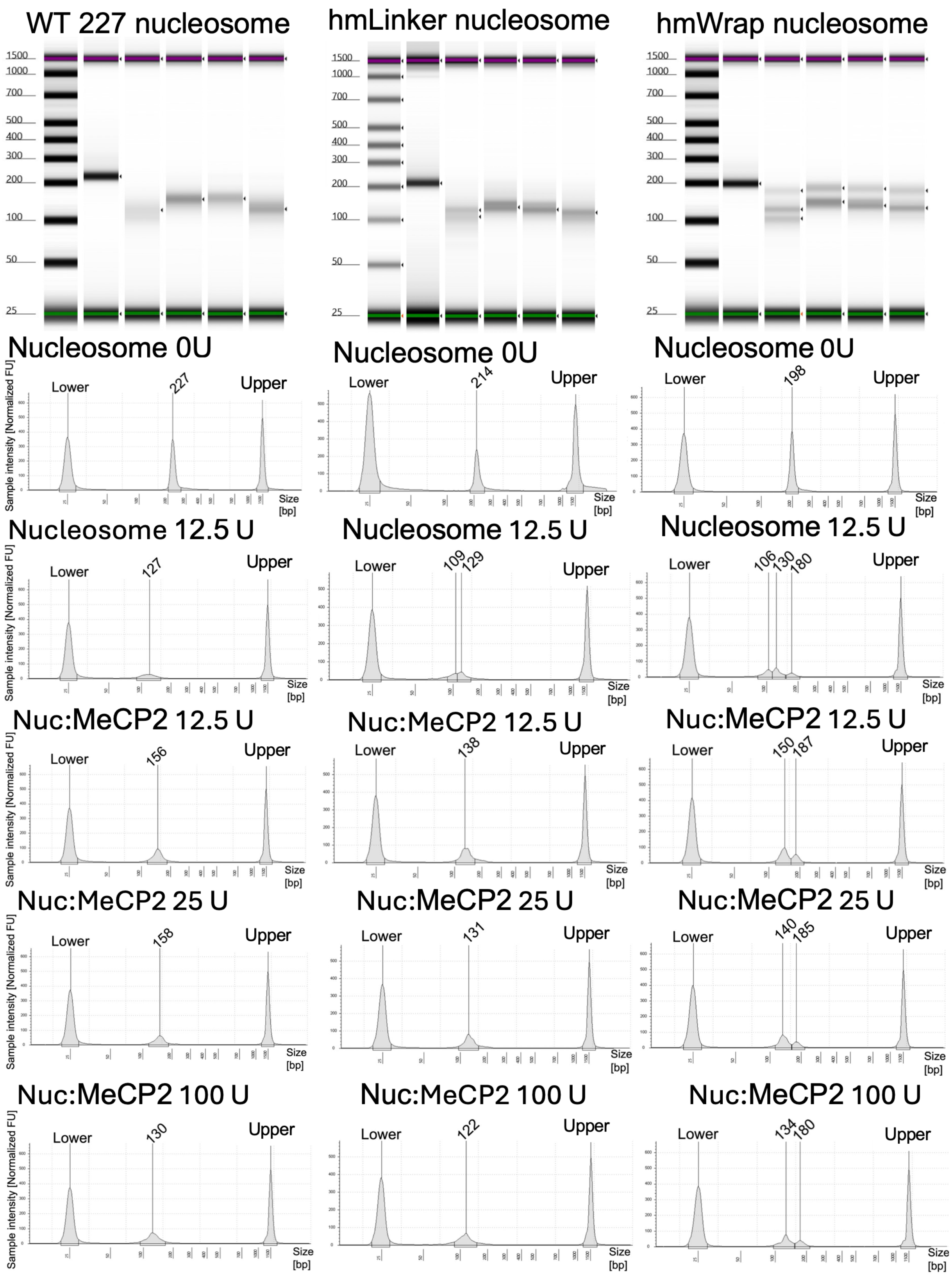

WT 227 nucleosome
hmLinker nucleosome
hmWrap nucleosome
1500
1000
700
500
400
300
200
100
50
25
Nucleosome 0U
Lower
227
214
198
Upper
Nucleosome 12.5 U
127
109
129
106
130
180
Nuc:MeCP2 12.5 U
156
138
150
187
Nuc:MeCP2 25 U
158
131
140
185
Nuc:MeCP2 100 U
130
122
134
180
Sample intensity [Normalized FU]
Size [bp]

**Supplementary Figure 7: MNase footprinting of MeCP2 bound to nucleosomes (Replicate 1).** For each nucleosomes, 5 samples are loaded in sequence: 0.5 μM nucleosomes that have not been treated with Mnase (nucleosome 0U), 0.5 μM nucleosomes treated with 12.5 U Mnase (nucleosome 12.5U), 0.5 uM nucleosomes+1.5 μM of MeCP2 treated with 12.5 U Mnase (Nucleosome: MeCP2 12.5 U), 0.5 μM nucleosomes+1.5 μM of MeCP2 treated with 25 U Mnase (Nucleosome: MeCP2 25 U), 0.5 μM nucleosomes+1.5 μM of MeCP2 treated with 100 U Mnase (Nucleosome: MeCP2 100 U). Each graph shows the lower and upper limits and the actual peaks of the reaction. The lower (35 bp) and upper (10,380 bp) limit peaks belong to internal standards in the loading buffer.

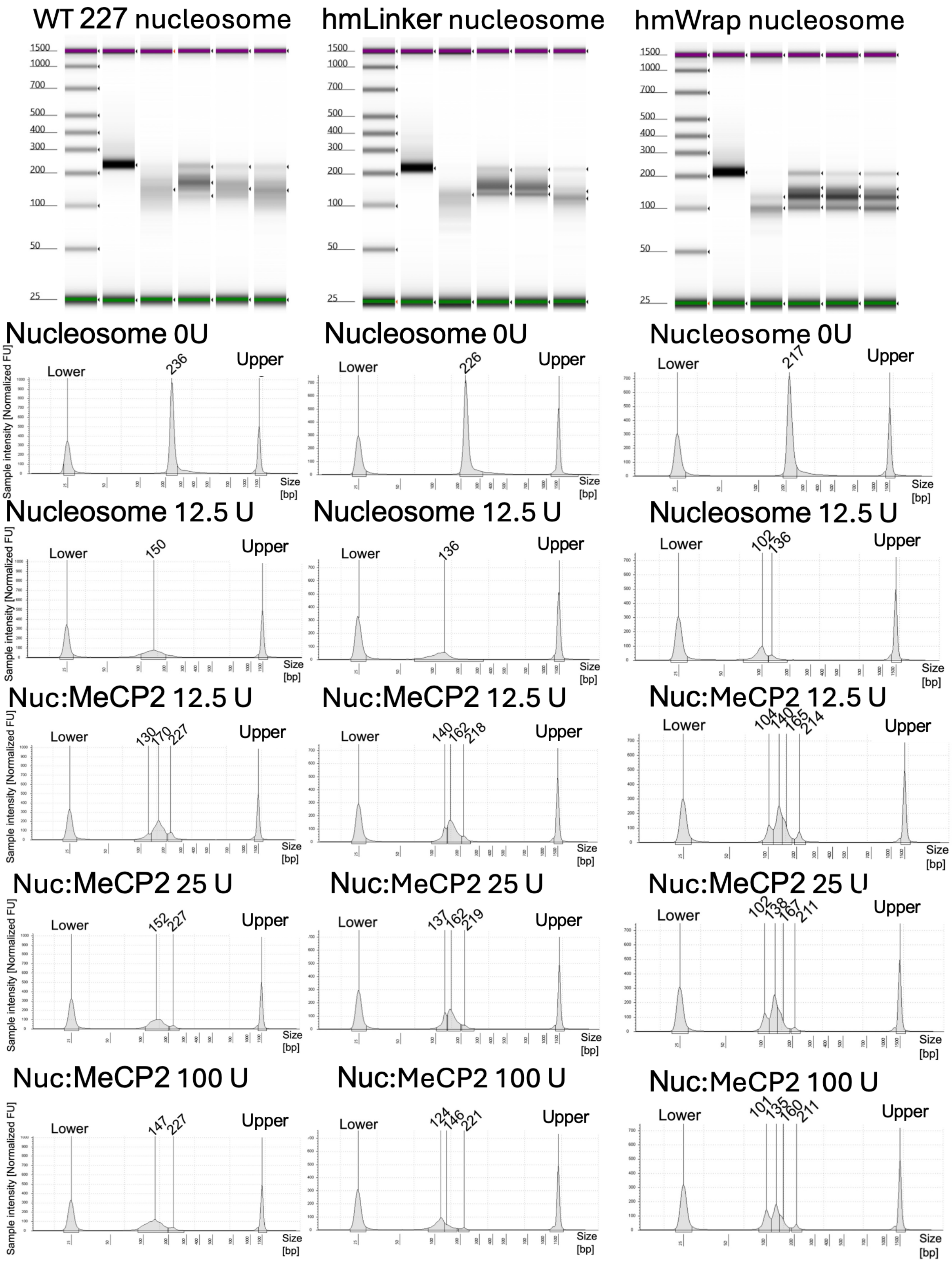

WT 227 nucleosome
hmLinker nucleosome
hmWrap nucleosome
Nucleosome 0U
Nucleosome 12.5 U
Nuc:MeCP2 12.5 U
Nuc:MeCP2 25 U
Nuc:MeCP2 100 U
Lower
Upper
Sample intensity [Normalized FU]
Size [bp]
236
226
217
150
136
102
136
130
170
227
140
162
218
104
140
165
214
152
227
137
162
219
102
138
167
211
147
227
124
146
221
101
135
160
211

**Supplementary Figure 8: MNase footprinting of MeCP2 bound to nucleosomes (Replicate 2; from independent MeCP2 prep and independent nucleosome reconstitutions).** For each nucleosomes, 5 samples are loaded in sequence: 0.5 μM nucleosomes that have not been treated with Mnase (nucleosome 0U), 0.5 μM nucleosomes treated with 12.5 U Mnase (nucleosome 12.5U), 0.5 uM nucleosomes+1.5 μM of MeCP2 treated with 12.5 U Mnase (Nucleosome: MeCP2 12.5 U), 0.5 μM nucleosomes+1.5 μM of MeCP2 treated with 25 U Mnase (Nucleosome: MeCP2 25 U), 0.5 μM nucleosomes+1.5 μM of MeCP2 treated with 100 U Mnase (Nucleosome: MeCP2 100 U). Each graph shows the lower and upper limits and the actual peaks of the reaction. The lower (35 bp) and upper (10,380 bp) limit peaks belong to internal standards in the loading buffer.

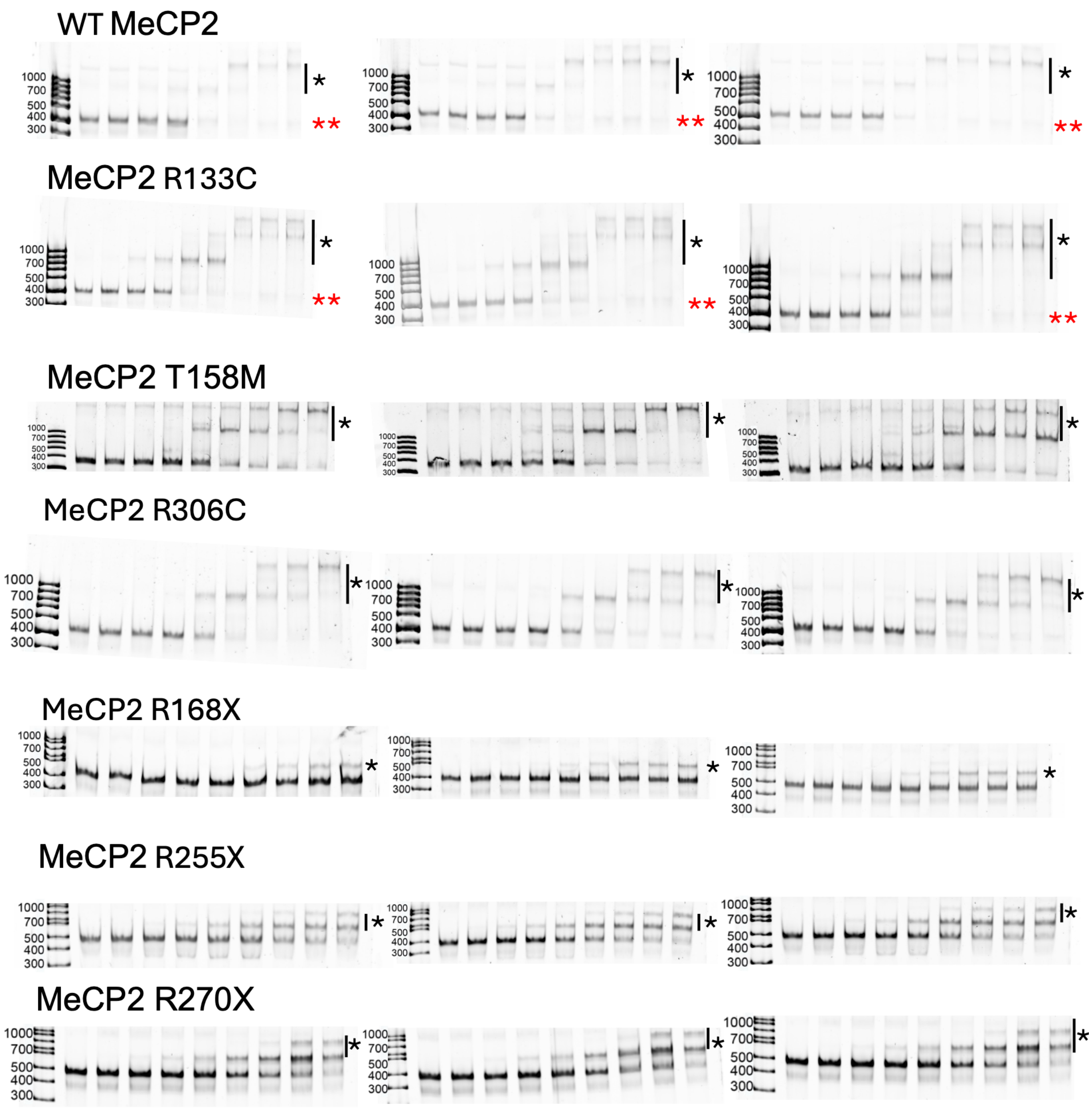


**Supplementary Figure 9: Triplicate EMSA assays for different constructs of MeCP2 and WT nucleosome.** Each lane contains 50 nM labeled nucleosome and increasing concentrations of MeCP2 (0–250 nM). Asterisks (*) indicate complete gel-shift to the bound species; red asterisks (**) mark species arising from non-specific binding to 27-bp competitor DNA.

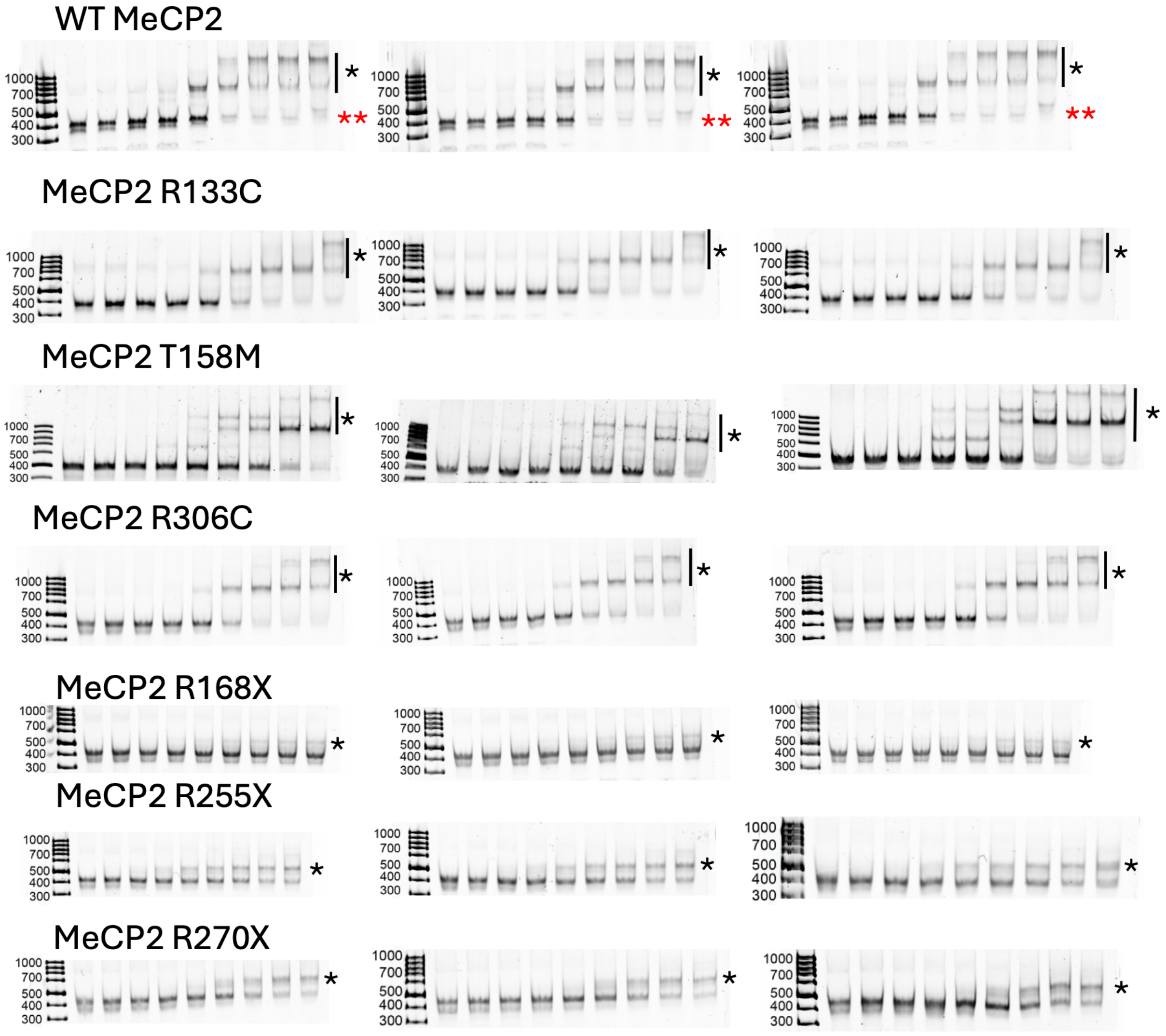


**Supplementary Figure 10: Triplicate EMSA assays for different constructs of MeCP2 and hmLinker nucleosome.** Each lane contains 50 nM labeled nucleosome and increasing concentrations of MeCP2 (0–250 nM). Asterisks (*) indicate complete gel-shift to the bound species; red asterisks (**) mark species arising from non-specific binding to 27-bp competitor DNA.

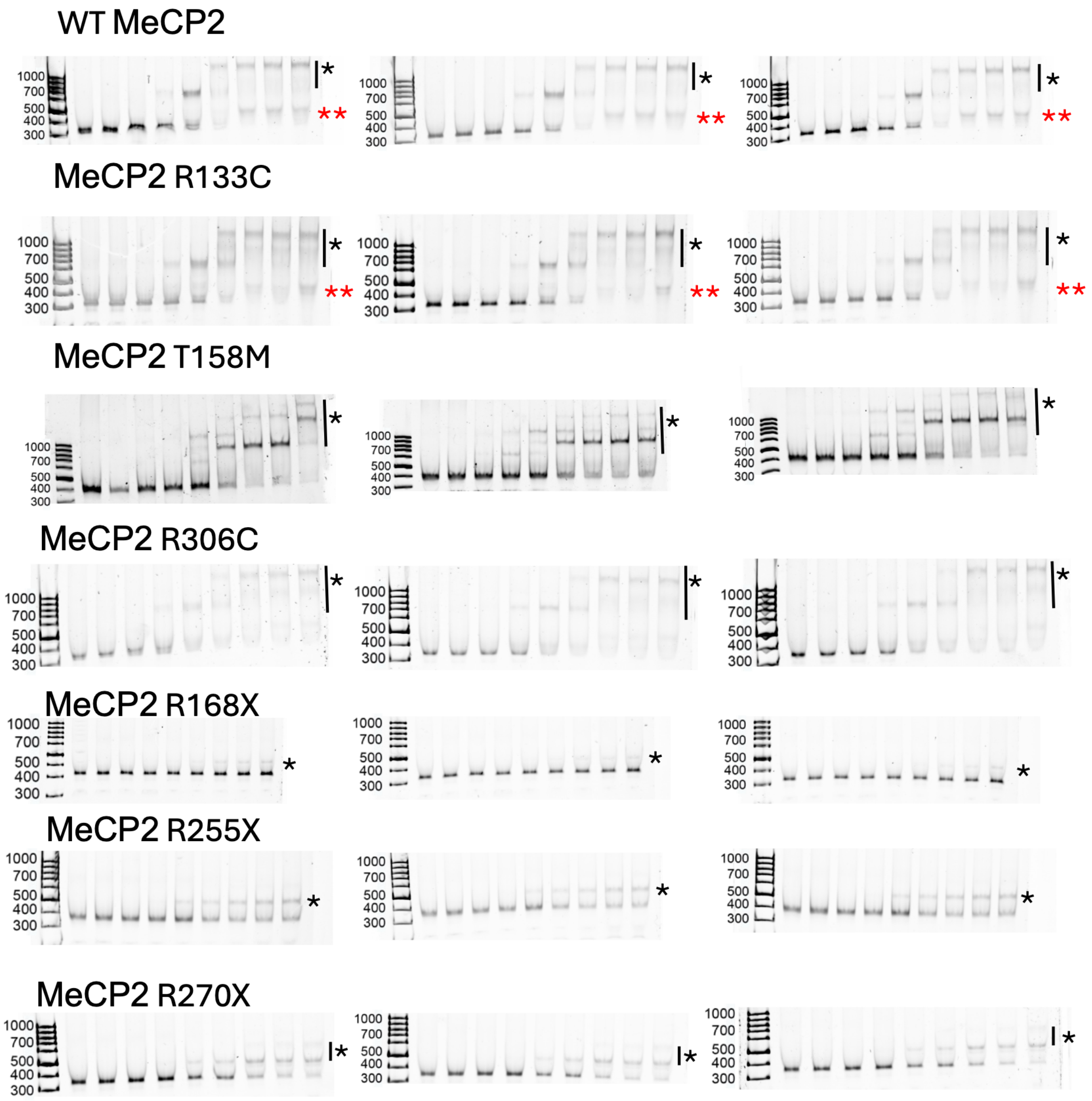


**Supplementary Figure 11: Triplicate EMSA assays for different constructs of MeCP2 and hmWrap nucleosome.** Each lane contains 50 nM labeled nucleosome and increasing concentrations of MeCP2 (0–250 nM). Asterisks (*) indicate the bound species; red asterisks (**) mark species arising from non-specific binding to 27-bp competitor DNA.

### Supplementary Table S1. EMSA fit parameters

*Kd reported as mean ± SE of per-replicate fits. Model = majority model across 3 replicates. AIC criterion: Hill selected if ΔAIC(Hill−Morrison) < −2. Nucleosome [L] = 50 nM. R168X hmWrap = N.D. (Fmax < 15%, uninterpretable). R168X 227WTnuc/hmLinker = >1 μM.*

| Nucleosome | Variant | Model (majority) | Kd (nM) mean | Kd SE (nM) | Kd_Rep1 (nM) | Kd_Rep2 (nM) | Kd_Rep3 (nM) | Fmax mean | Hill n mean | $R^2$ mean | ΔAIC (Hill−Morrison) | N reps |
|---|---|---|---|---|---|---|---|---|---|---|---|---|
| **WT 227 Nucleosome (40-N-40)** | | | | | | | | | | | | |
| WT 227 Nucleosome (40-N-40) | WT | Hill | 88.80 | 1.27 | 90.80 | 86.45 | 89.15 | 0.99 | 4.09 | 1.00 | -37.07 | 3 |
| WT 227 Nucleosome (40-N-40) | R133C | Hill | 64.30 | 5.39 | 57.35 | 74.91 | 60.64 | 1.02 | 2.35 | 0.99 | -9.80 | 3 |
| WT 227 Nucleosome (40-N-40) | T158M | Hill | 187.82 | 8.46 | 176.75 | 204.43 | 182.27 | 1.50 | 1.85 | 0.96 | -7.10 | 3 |
| WT 227 Nucleosome (40-N-40) | R255X | Hill | 266.52 | 12.45 | 255.59 | 252.60 | 291.37 | 1.50 | 1.71 | 0.99 | -12.97 | 3 |
| WT 227 Nucleosome (40-N-40) | R306C | Hill | 178.14 | 7.75 | 189.81 | 181.15 | 163.48 | 1.40 | 2.97 | 0.98 | -14.93 | 3 |
| WT 227 Nucleosome (40-N-40) | R270X | Hill | 262.39 | 23.57 | 290.77 | 280.81 | 215.60 | 1.37 | 2.00 | 0.99 | -19.33 | 3 |
| WT 227 Nucleosome (40-N-40) | R168X | Hill | >1000 | n.d | >1000 | >1000 | >1000 | 1.12 | 2.35 | 0.98 | -6.90 | 3 |
| **hmLinker Nucleosome (23-N-40)** | | | | | | | | | | | | |
| hmLinker Nucleosome (23-N-40) | WT | Hill | 103.24 | 2.72 | 98.76 | 102.80 | 108.16 | 1.09 | 2.75 | 0.99 | -18.90 | 3 |
| hmLinker Nucleosome (23-N-40) | R133C | Hill | 131.03 | 3.62 | 129.16 | 138.02 | 125.90 | 1.04 | 4.41 | 1.00 | -40.80 | 3 |
| hmLinker Nucleosome (23-N-40) | T158M | Hill | 232.23 | 11.10 | 213.48 | 231.31 | 251.90 | 1.50 | 3.64 | 0.92 | -8.30 | 3 |
| hmLinker Nucleosome (23-N-40) | R255X | Morrison | 283.05 | 56.00 | 350.67 | 171.91 | 326.56 | 1.31 | | 0.96 | 1.53 | 3 |
| hmLinker Nucleosome (23-N-40) | R306C | Hill | 150.31 | 1.68 | 153.61 | 149.13 | 148.18 | 1.04 | 6.00 | 1.00 | -47.73 | 3 |
| hmLinker Nucleosome (23-N-40) | R270X | Hill | 247.78 | 17.49 | 212.91 | 262.73 | 267.69 | 1.41 | 2.43 | 0.99 | -24.50 | 3 |
| hmLinker Nucleosome (23-N-40) | R168X | Morrison | >1000 | n.d | >1000 | >1000 | >1000 | 0.96 | | 0.95 | 1.76 | 3 |
| **hmWrap Nucleosome (14-N-40)** | | | | | | | | | | | | |
| hmWrap Nucleosome (14-N-40) | WT | Hill | 85.50 | 5.57 | 96.65 | 79.92 | 79.94 | 1.01 | 4.21 | 1.00 | -39.87 | 3 |
| hmWrap Nucleosome (14-N-40) | R133C | Hill | 116.46 | 7.27 | 130.03 | 114.18 | 105.16 | 1.19 | 2.17 | 0.99 | -18.90 | 3 |
| hmWrap Nucleosome (14-N-40) | T158M | Hill | 185.40 | 20.59 | 144.33 | 208.53 | 203.34 | 1.36 | 2.54 | 0.98 | -12.40 | 3 |
| hmWrap Nucleosome (14-N-40) | R255X | Hill | 248.50 | 35.33 | 309.90 | 187.52 | 248.07 | 0.71 | 2.67 | 0.99 | -27.40 | 3 |
| hmWrap Nucleosome (14-N-40) | R306C | Hill | 103.33 | 25.06 | 153.42 | 77.10 | 79.47 | 1.16 | 1.66 | 0.98 | -3.63 | 3 |
| hmWrap Nucleosome (14-N-40) | R270X | Hill | 213.06 | 20.69 | 192.46 | 254.44 | 192.28 | 1.11 | 4.62 | 1.00 | -37.07 | 3 |
| hmWrap Nucleosome (14-N-40) | R168X | Hill | >1000 | n.d | >1000 | >1000 | >1000 | 0.59 | 3.27 | 0.97 | -13.47 | 3 |

| length | competitor | protein | DNA | replicates | Kd_nM | Kd_SE_nM |
|---|---|---|---|---|---|---|
| 15bp | competitor | R133C | Hemi-methylated | 3 | 81.26 | 4.50 |
| 15bp | competitor | R133C | Methylated | 3 | 78.28 | 8.31 |
| 15bp | competitor | R133C | Unmethylated | 3 | 81.18 | 7.01 |
| 15bp | competitor | R168X | Hemi-methylated | 3 | 218.30 | 8.34 |
| 15bp | competitor | R168X | Methylated | 3 | 100.36 | 5.11 |
| 15bp | competitor | R168X | Unmethylated | 3 | 300.73 | 14.91 |
| 15bp | competitor | R255X | Hemi-methylated | 3 | 102.12 | 2.67 |
| 15bp | competitor | R255X | Methylated | 3 | 108.24 | 4.33 |
| 15bp | competitor | R255X | Unmethylated | 3 | 119.65 | 8.43 |
| 15bp | competitor | R270X | Hemi-methylated | 3 | 100.88 | 0.85 |
| 15bp | competitor | R270X | Methylated | 3 | 85.41 | 2.96 |
| 15bp | competitor | R270X | Unmethylated | 3 | 108.93 | 0.84 |
| 15bp | competitor | R306C | Hemi-methylated | 3 | 71.00 | 9.43 |
| 15bp | competitor | R306C | Methylated | 3 | 56.52 | 7.32 |
| 15bp | competitor | R306C | Unmethylated | 3 | 79.41 | 9.62 |
| 15bp | competitor | T158M | Hemi-methylated | 3 | 67.30 | 5.18 |
| 15bp | competitor | T158M | Methylated | 3 | 60.75 | 4.61 |
| 15bp | competitor | T158M | Unmethylated | 3 | 71.34 | 6.67 |
| 15bp | competitor | WT | Hemi-methylated | 3 | 51.56 | 2.46 |
| 15bp | competitor | WT | Methylated | 3 | 47.51 | 0.17 |
| 15bp | competitor | WT | Unmethylated | 3 | 51.17 | 0.84 |
| 15bp | none | R133C | Hemi-methylated | 3 | 2.19 | 0.33 |
| 15bp | none | R133C | Methylated | 3 | 2.13 | 0.38 |
| 15bp | none | R133C | Unmethylated | 3 | 2.57 | 0.48 |
| 15bp | none | R168X | Hemi-methylated | 3 | 90.02 | 10.02 |
| 15bp | none | R168X | Methylated | 3 | 15.69 | 2.23 |
| 15bp | none | R168X | Unmethylated | 3 | 297.86 | 20.32 |
| 15bp | none | R255X | Hemi-methylated | 3 | 51.14 | 3.54 |
| 15bp | none | R255X | Methylated | 3 | 57.61 | 2.19 |
| 15bp | none | R255X | Unmethylated | 3 | 64.01 | 6.67 |
| 15bp | none | R270X | Hemi-methylated | 3 | 2.16 | 0.33 |
| 15bp | none | R270X | Methylated | 3 | 1.35 | 0.15 |
| 15bp | none | R270X | Unmethylated | 3 | 7.06 | 1.12 |
| 15bp | none | R306C | Hemi-methylated | 3 | 0.66 | 0.09 |
| 15bp | none | R306C | Methylated | 3 | 1.35 | 0.54 |
| 15bp | none | R306C | Unmethylated | 3 | 1.11 | 0.16 |
| 15bp | none | T158M | Hemi-methylated | 3 | 2.00 | 0.46 |
| 15bp | none | T158M | Methylated | 3 | 2.22 | 0.17 |
| 15bp | none | T158M | Unmethylated | 3 | 2.00 | 0.67 |
| 15bp | none | WT | Hemi-methylated | 3 | 1.04 | 0.06 |
| 15bp | none | WT | Methylated | 3 | 1.44 | 0.36 |
| 15bp | none | WT | Unmethylated | 3 | 1.38 | 0.13 |
| 27bp | competitor | R133C | Hemi-methylated | 3 | 72.62 | 0.20 |
| 27bp | competitor | R133C | Methylated | 3 | 65.29 | 1.05 |

| | | | | | | |
|---|---|---|---|---|---|---|
| 27bp | competitor | R133C | Unmethylated | 3 | 58.95 | 3.29 |
| 27bp | competitor | R168X | Hemi-methylated | 3 | 219.21 | 6.78 |
| 27bp | competitor | R168X | Methylated | 3 | 306.60 | 21.14 |
| 27bp | competitor | R168X | Unmethylated | 3 | 286.45 | 5.59 |
| 27bp | competitor | R255X | Hemi-methylated | 3 | 112.78 | 16.62 |
| 27bp | competitor | R255X | Methylated | 3 | 116.21 | 18.72 |
| 27bp | competitor | R255X | Unmethylated | 3 | 119.49 | 19.09 |
| 27bp | competitor | R270X | Hemi-methylated | 3 | 149.16 | 6.27 |
| 27bp | competitor | R270X | Methylated | 3 | 161.03 | 5.43 |
| 27bp | competitor | R270X | Unmethylated | 3 | 163.59 | 5.66 |
| 27bp | competitor | R306C | Hemi-methylated | 3 | 57.57 | 2.11 |
| 27bp | competitor | R306C | Methylated | 3 | 47.94 | 0.70 |
| 27bp | competitor | R306C | Unmethylated | 3 | 51.54 | 3.42 |
| 27bp | competitor | T158M | Hemi-methylated | 3 | 66.73 | 2.77 |
| 27bp | competitor | T158M | Methylated | 3 | 56.19 | 0.45 |
| 27bp | competitor | T158M | Unmethylated | 3 | 65.89 | 3.46 |
| 27bp | competitor | WT | Hemi-methylated | 3 | 74.94 | 3.46 |
| 27bp | competitor | WT | Methylated | 3 | 72.20 | 2.02 |
| 27bp | competitor | WT | Unmethylated | 3 | 59.95 | 1.77 |
| 27bp | none | R133C | Hemi-methylated | 3 | 1.55 | 0.28 |
| 27bp | none | R133C | Methylated | 3 | 2.69 | 0.44 |
| 27bp | none | R133C | Unmethylated | 3 | 2.27 | 0.63 |
| 27bp | none | R168X | Hemi-methylated | 3 | 31.02 | 6.61 |
| 27bp | none | R168X | Methylated | 3 | 146.88 | 12.74 |
| 27bp | none | R168X | Unmethylated | 3 | 108.32 | 25.43 |
| 27bp | none | R255X | Hemi-methylated | 3 | 79.61 | 5.17 |
| 27bp | none | R255X | Methylated | 3 | 93.88 | 2.77 |
| 27bp | none | R255X | Unmethylated | 3 | 85.32 | 6.62 |
| 27bp | none | R270X | Hemi-methylated | 3 | 4.00 | 0.57 |
| 27bp | none | R270X | Methylated | 3 | 7.59 | 0.83 |
| 27bp | none | R270X | Unmethylated | 3 | 6.06 | 0.58 |
| 27bp | none | R306C | Hemi-methylated | 3 | 1.24 | 0.58 |
| 27bp | none | R306C | Methylated | 3 | 1.86 | 0.26 |
| 27bp | none | R306C | Unmethylated | 3 | 1.65 | 0.53 |
| 27bp | none | T158M | Hemi-methylated | 3 | 0.56 | 0.31 |
| 27bp | none | T158M | Methylated | 3 | 1.21 | 0.17 |
| 27bp | none | T158M | Unmethylated | 3 | 1.82 | 0.53 |
| 27bp | none | WT | Hemi-methylated | 3 | 0.73 | 0.05 |
| 27bp | none | WT | Methylated | 3 | 0.76 | 0.03 |
| 27bp | none | WT | Unmethylated | 3 | 0.72 | 0.21 |

**Supplementary Table S3. Actual Kd for each replicate and different variants of MeCP2 binding to DNA in FP**

| length | competitor | protein | dna | replicate | Kd_nM | Kd_SE | model | $R^2$ |
|---|---|---|---|---|---|---|---|---|
| 15bp | competitor | R133C | Hemi-methylated | 1 | 84.74 | 1.98 | Hill | 1.00 |
| 15bp | competitor | R133C | Hemi-methylated | 2 | 72.34 | 2.58 | Hill | 1.00 |
| 15bp | competitor | R133C | Hemi-methylated | 3 | 86.69 | 2.56 | Hill | 1.00 |
| 15bp | competitor | R133C | Methylated | 1 | 81.35 | 1.87 | Hill | 1.00 |
| 15bp | competitor | R133C | Methylated | 2 | 62.60 | 1.94 | Hill | 1.00 |
| 15bp | competitor | R133C | Methylated | 3 | 90.88 | 1.82 | Hill | 1.00 |
| 15bp | competitor | R133C | Unmethylated | 1 | 78.20 | 1.91 | Hill | 1.00 |
| 15bp | competitor | R133C | Unmethylated | 2 | 70.82 | 2.23 | Hill | 1.00 |
| 15bp | competitor | R133C | Unmethylated | 3 | 94.54 | 2.25 | Hill | 1.00 |
| 15bp | competitor | R168X | Hemi-methylated | 1 | 225.99 | 24.22 | Hill | 0.99 |
| 15bp | competitor | R168X | Hemi-methylated | 2 | 201.64 | 15.71 | Hill | 0.99 |
| 15bp | competitor | R168X | Hemi-methylated | 3 | 227.28 | 12.99 | Hill | 0.99 |
| 15bp | competitor | R168X | Methylated | 1 | 100.67 | 9.74 | Hill | 0.99 |
| 15bp | competitor | R168X | Methylated | 2 | 91.37 | 8.78 | Hill | 0.99 |
| 15bp | competitor | R168X | Methylated | 3 | 109.05 | 8.79 | Hill | 1.00 |
| 15bp | competitor | R168X | Unmethylated | 1 | 330.14 | 19.84 | Hill | 1.00 |
| 15bp | competitor | R168X | Unmethylated | 2 | 290.29 | 10.05 | Hill | 1.00 |
| 15bp | competitor | R168X | Unmethylated | 3 | 281.76 | 12.11 | Hill | 1.00 |
| 15bp | competitor | R255X | Hemi-methylated | 1 | 106.93 | 4.04 | Hill | 1.00 |
| 15bp | competitor | R255X | Hemi-methylated | 2 | 97.72 | 2.91 | Hill | 1.00 |
| 15bp | competitor | R255X | Hemi-methylated | 3 | 101.72 | 2.24 | Hill | 1.00 |
| 15bp | competitor | R255X | Methylated | 1 | 116.89 | 2.57 | Hill | 1.00 |
| 15bp | competitor | R255X | Methylated | 2 | 104.17 | 3.50 | Hill | 1.00 |
| 15bp | competitor | R255X | Methylated | 3 | 103.64 | 2.28 | Hill | 1.00 |
| 15bp | competitor | R255X | Unmethylated | 1 | 136.39 | 3.03 | Hill | 1.00 |
| 15bp | competitor | R255X | Unmethylated | 2 | 113.04 | 2.67 | Hill | 1.00 |
| 15bp | competitor | R255X | Unmethylated | 3 | 109.53 | 3.56 | Hill | 0.99 |
| 15bp | competitor | R270X | Hemi-methylated | 1 | 102.34 | 3.45 | Hill | 1.00 |
| 15bp | competitor | R270X | Hemi-methylated | 2 | 100.89 | 3.39 | Hill | 1.00 |
| 15bp | competitor | R270X | Hemi-methylated | 3 | 99.41 | 4.94 | Hill | 0.99 |
| 15bp | competitor | R270X | Methylated | 1 | 90.93 | 4.90 | Hill | 0.99 |
| 15bp | competitor | R270X | Methylated | 2 | 80.78 | 8.24 | Hill | 0.98 |
| 15bp | competitor | R270X | Methylated | 3 | 84.50 | 4.37 | Hill | 0.99 |
| 15bp | competitor | R270X | Unmethylated | 1 | 109.36 | 4.96 | Hill | 0.99 |
| 15bp | competitor | R270X | Unmethylated | 2 | 110.11 | 5.21 | Hill | 0.99 |
| 15bp | competitor | R270X | Unmethylated | 3 | 107.31 | 4.12 | Hill | 0.99 |
| 15bp | competitor | R306C | Hemi-methylated | 1 | 68.63 | 1.92 | Hill | 1.00 |
| 15bp | competitor | R306C | Hemi-methylated | 2 | 55.99 | 2.69 | Hill | 0.99 |
| 15bp | competitor | R306C | Hemi-methylated | 3 | 88.39 | 2.45 | Hill | 1.00 |
| 15bp | competitor | R306C | Methylated | 1 | 51.07 | 2.45 | Hill | 0.99 |
| 15bp | competitor | R306C | Methylated | 2 | 47.48 | 3.40 | Hill | 0.99 |
| 15bp | competitor | R306C | Methylated | 3 | 71.02 | 1.64 | Hill | 1.00 |
| 15bp | competitor | R306C | Unmethylated | 1 | 91.64 | 1.75 | Hill | 1.00 |
| 15bp | competitor | R306C | Unmethylated | 2 | 86.18 | 1.44 | Hill | 1.00 |
| 15bp | competitor | R306C | Unmethylated | 3 | 60.43 | 2.03 | Hill | 1.00 |
| 15bp | competitor | T158M | Hemi-methylated | 1 | 74.67 | 2.36 | Hill | 1.00 |
| 15bp | competitor | T158M | Hemi-methylated | 2 | 57.32 | 1.00 | Hill | 1.00 |
| 15bp | competitor | T158M | Hemi-methylated | 3 | 69.90 | 1.25 | Hill | 1.00 |
| 15bp | competitor | T158M | Methylated | 1 | 53.89 | 1.49 | Hill | 1.00 |

| | | | | | | | | |
|---|---|---|---|---|---|---|---|---|
| 15bp | competitor | T158M | Methylated | 2 | 58.86 | 1.14 | Hill | 1.00 |
| 15bp | competitor | T158M | Methylated | 3 | 69.50 | 2.96 | Hill | 0.99 |
| 15bp | competitor | T158M | Unmethylated | 1 | 75.68 | 3.25 | Hill | 0.99 |
| 15bp | competitor | T158M | Unmethylated | 2 | 58.24 | 1.93 | Hill | 1.00 |
| 15bp | competitor | T158M | Unmethylated | 3 | 80.10 | 4.27 | Hill | 0.99 |
| 15bp | competitor | WT | Hemi-methylated | 1 | 48.84 | 1.83 | Hill | 1.00 |
| 15bp | competitor | WT | Hemi-methylated | 2 | 49.37 | 1.99 | Hill | 1.00 |
| 15bp | competitor | WT | Hemi-methylated | 3 | 56.48 | 1.30 | Hill | 1.00 |
| 15bp | competitor | WT | Methylated | 1 | 47.18 | 2.12 | Hill | 1.00 |
| 15bp | competitor | WT | Methylated | 2 | 47.64 | 2.28 | Hill | 0.99 |
| 15bp | competitor | WT | Methylated | 3 | 47.71 | 1.66 | Hill | 1.00 |
| 15bp | competitor | WT | Unmethylated | 1 | 52.73 | 1.76 | Hill | 1.00 |
| 15bp | competitor | WT | Unmethylated | 2 | 50.93 | 1.40 | Hill | 1.00 |
| 15bp | competitor | WT | Unmethylated | 3 | 49.86 | 1.60 | Hill | 1.00 |
| 15bp | none | R133C | Hemi-methylated | 1 | 1.82 | 0.11 | Hill | 0.99 |
| 15bp | none | R133C | Hemi-methylated | 2 | 1.92 | 0.11 | Hill | 0.99 |
| 15bp | none | R133C | Hemi-methylated | 3 | 2.85 | 0.11 | Hill | 1.00 |
| 15bp | none | R133C | Methylated | 1 | 1.74 | 0.04 | Hill | 1.00 |
| 15bp | none | R133C | Methylated | 2 | 1.76 | 0.05 | Hill | 1.00 |
| 15bp | none | R133C | Methylated | 3 | 2.90 | 0.11 | Hill | 1.00 |
| 15bp | none | R133C | Unmethylated | 1 | 2.08 | 0.17 | Hill | 0.99 |
| 15bp | none | R133C | Unmethylated | 2 | 2.12 | 0.12 | Hill | 1.00 |
| 15bp | none | R133C | Unmethylated | 3 | 3.52 | 0.29 | Hill | 0.99 |
| 15bp | none | R168X | Hemi-methylated | 1 | 73.17 | 5.64 | Morrison | 1.00 |
| 15bp | none | R168X | Hemi-methylated | 2 | 89.05 | 8.23 | Morrison | 0.99 |
| 15bp | none | R168X | Hemi-methylated | 3 | 107.84 | 13.07 | Hill | 1.00 |
| 15bp | none | R168X | Methylated | 1 | 18.87 | 2.89 | Morrison | 0.98 |
| 15bp | none | R168X | Methylated | 2 | 16.79 | 3.10 | Morrison | 0.98 |
| 15bp | none | R168X | Methylated | 3 | 11.40 | 1.68 | Morrison | 0.99 |
| 15bp | none | R168X | Unmethylated | 1 | 275.09 | 19.34 | Morrison | 1.00 |
| 15bp | none | R168X | Unmethylated | 2 | 338.40 | 37.68 | Morrison | 0.99 |
| 15bp | none | R168X | Unmethylated | 3 | 280.09 | 29.45 | Morrison | 0.99 |
| 15bp | none | R255X | Hemi-methylated | 1 | 49.80 | 2.86 | Hill | 0.99 |
| 15bp | none | R255X | Hemi-methylated | 2 | 45.78 | 2.69 | Hill | 1.00 |
| 15bp | none | R255X | Hemi-methylated | 3 | 57.83 | 2.71 | Hill | 0.99 |
| 15bp | none | R255X | Methylated | 1 | 61.75 | 4.58 | Hill | 0.99 |
| 15bp | none | R255X | Methylated | 2 | 54.32 | 4.25 | Hill | 0.99 |
| 15bp | none | R255X | Methylated | 3 | 56.76 | 4.49 | Hill | 0.99 |
| 15bp | none | R255X | Unmethylated | 1 | 59.99 | 2.36 | Hill | 1.00 |
| 15bp | none | R255X | Unmethylated | 2 | 55.00 | 1.64 | Hill | 1.00 |
| 15bp | none | R255X | Unmethylated | 3 | 77.04 | 3.74 | Hill | 1.00 |
| 15bp | none | R270X | Hemi-methylated | 1 | 1.53 | 0.37 | Morrison | 0.97 |
| 15bp | none | R270X | Hemi-methylated | 2 | 2.65 | 0.47 | Hill | 0.98 |
| 15bp | none | R270X | Hemi-methylated | 3 | 2.29 | 0.40 | Hill | 0.98 |
| 15bp | none | R270X | Methylated | 1 | 1.65 | 0.38 | Morrison | 0.97 |
| 15bp | none | R270X | Methylated | 2 | 1.22 | 0.30 | Morrison | 0.97 |
| 15bp | none | R270X | Methylated | 3 | 1.17 | 0.24 | Morrison | 0.98 |
| 15bp | none | R270X | Unmethylated | 1 | 9.28 | 4.01 | Morrison | 0.86 |
| 15bp | none | R270X | Unmethylated | 2 | 5.70 | 1.47 | Hill | 0.98 |
| 15bp | none | R270X | Unmethylated | 3 | 6.19 | 1.75 | Hill | 0.97 |
| 15bp | none | R306C | Hemi-methylated | 1 | 0.61 | 0.12 | Morrison | 0.99 |

| | | | | | | | | |
|---|---|---|---|---|---|---|---|---|
| 15bp | none | R306C | Hemi-methylated | 2 | 0.84 | 0.09 | Hill | 0.99 |
| 15bp | none | R306C | Hemi-methylated | 3 | 0.53 | 0.12 | Morrison | 0.99 |
| 15bp | none | R306C | Methylated | 1 | 2.00 | 0.10 | Hill | 0.99 |
| 15bp | none | R306C | Methylated | 2 | 0.28 | 0.07 | Morrison | 0.99 |
| 15bp | none | R306C | Methylated | 3 | 1.77 | 0.10 | Hill | 0.99 |
| 15bp | none | R306C | Unmethylated | 1 | 1.12 | 0.18 | Morrison | 0.99 |
| 15bp | none | R306C | Unmethylated | 2 | 1.39 | 0.19 | Hill | 0.98 |
| 15bp | none | R306C | Unmethylated | 3 | 0.83 | 0.20 | Morrison | 0.98 |
| 15bp | none | T158M | Hemi-methylated | 1 | 2.10 | 0.16 | Hill | 0.99 |
| 15bp | none | T158M | Hemi-methylated | 2 | 1.16 | 0.25 | Morrison | 0.98 |
| 15bp | none | T158M | Hemi-methylated | 3 | 2.75 | 0.16 | Hill | 1.00 |
| 15bp | none | T158M | Methylated | 1 | 2.05 | 0.11 | Hill | 0.99 |
| 15bp | none | T158M | Methylated | 2 | 2.05 | 0.13 | Hill | 0.99 |
| 15bp | none | T158M | Methylated | 3 | 2.56 | 0.16 | Hill | 0.99 |
| 15bp | none | T158M | Unmethylated | 1 | 3.33 | 0.30 | Hill | 0.97 |
| 15bp | none | T158M | Unmethylated | 2 | 1.30 | 0.28 | Morrison | 0.98 |
| 15bp | none | T158M | Unmethylated | 3 | 1.37 | 0.21 | Morrison | 0.99 |
| 15bp | none | WT | Hemi-methylated | 1 | 0.92 | 0.12 | Hill | 0.99 |
| 15bp | none | WT | Hemi-methylated | 2 | 1.09 | 0.16 | Hill | 0.97 |
| 15bp | none | WT | Hemi-methylated | 3 | 1.10 | 0.20 | Morrison | 0.99 |
| 15bp | none | WT | Methylated | 1 | 1.12 | 0.14 | Hill | 0.98 |
| 15bp | none | WT | Methylated | 2 | 1.03 | 0.07 | Hill | 0.99 |
| 15bp | none | WT | Methylated | 3 | 2.16 | 0.19 | Hill | 0.98 |
| 15bp | none | WT | Unmethylated | 1 | 1.35 | 0.18 | Hill | 0.99 |
| 15bp | none | WT | Unmethylated | 2 | 1.18 | 0.18 | Hill | 0.98 |
| 15bp | none | WT | Unmethylated | 3 | 1.63 | 0.34 | Morrison | 0.98 |
| 27bp | competitor | R133C | Hemi-methylated | 1 | 72.25 | 3.63 | Hill | 0.99 |
| 27bp | competitor | R133C | Hemi-methylated | 2 | 72.64 | 2.95 | Hill | 1.00 |
| 27bp | competitor | R133C | Hemi-methylated | 3 | 72.96 | 2.43 | Hill | 1.00 |
| 27bp | competitor | R133C | Methylated | 1 | 66.85 | 1.56 | Hill | 1.00 |
| 27bp | competitor | R133C | Methylated | 2 | 63.28 | 1.90 | Hill | 1.00 |
| 27bp | competitor | R133C | Methylated | 3 | 65.73 | 2.92 | Hill | 1.00 |
| 27bp | competitor | R133C | Unmethylated | 1 | 53.46 | 3.39 | Hill | 0.99 |
| 27bp | competitor | R133C | Unmethylated | 2 | 58.56 | 2.51 | Hill | 0.99 |
| 27bp | competitor | R133C | Unmethylated | 3 | 64.82 | 3.65 | Hill | 0.98 |
| 27bp | competitor | R168X | Hemi-methylated | 1 | 217.83 | 7.60 | Hill | 1.00 |
| 27bp | competitor | R168X | Hemi-methylated | 2 | 231.59 | 23.70 | Hill | 1.00 |
| 27bp | competitor | R168X | Hemi-methylated | 3 | 208.23 | 19.30 | Hill | 0.99 |
| 27bp | competitor | R168X | Methylated | 1 | 335.17 | 23.03 | Morrison | 1.00 |
| 27bp | competitor | R168X | Methylated | 2 | 265.32 | 41.44 | Hill | 0.99 |
| 27bp | competitor | R168X | Methylated | 3 | 319.32 | 70.00 | Morrison | 0.98 |
| 27bp | competitor | R168X | Unmethylated | 1 | 295.70 | 35.25 | Hill | 0.99 |
| 27bp | competitor | R168X | Unmethylated | 2 | 287.27 | 21.13 | Hill | 1.00 |
| 27bp | competitor | R168X | Unmethylated | 3 | 276.38 | 24.89 | Hill | 1.00 |
| 27bp | competitor | R255X | Hemi-methylated | 1 | 135.41 | 5.74 | Hill | 0.99 |
| 27bp | competitor | R255X | Hemi-methylated | 2 | 122.54 | 5.29 | Hill | 0.99 |
| 27bp | competitor | R255X | Hemi-methylated | 3 | 80.39 | 2.33 | Hill | 1.00 |
| 27bp | competitor | R255X | Methylated | 1 | 143.32 | 8.30 | Hill | 0.99 |
| 27bp | competitor | R255X | Methylated | 2 | 125.03 | 4.45 | Hill | 1.00 |
| 27bp | competitor | R255X | Methylated | 3 | 80.29 | 2.03 | Hill | 1.00 |
| 27bp | competitor | R255X | Unmethylated | 1 | 152.04 | 3.25 | Hill | 1.00 |

| | | | | | | | | |
|---|---|---|---|---|---|---|---|---|
| 27bp | competitor | R255X | Unmethylated | 2 | 120.52 | 2.50 | Hill | 1.00 |
| 27bp | competitor | R255X | Unmethylated | 3 | 85.92 | 3.08 | Hill | 1.00 |
| 27bp | competitor | R270X | Hemi-methylated | 1 | 155.59 | 6.82 | Hill | 0.99 |
| 27bp | competitor | R270X | Hemi-methylated | 2 | 155.28 | 3.67 | Hill | 1.00 |
| 27bp | competitor | R270X | Hemi-methylated | 3 | 136.62 | 3.38 | Hill | 1.00 |
| 27bp | competitor | R270X | Methylated | 1 | 154.85 | 7.39 | Hill | 0.99 |
| 27bp | competitor | R270X | Methylated | 2 | 171.85 | 8.45 | Hill | 0.99 |
| 27bp | competitor | R270X | Methylated | 3 | 156.39 | 7.64 | Hill | 0.99 |
| 27bp | competitor | R270X | Unmethylated | 1 | 166.93 | 7.81 | Hill | 0.99 |
| 27bp | competitor | R270X | Unmethylated | 2 | 152.56 | 4.83 | Hill | 1.00 |
| 27bp | competitor | R270X | Unmethylated | 3 | 171.29 | 6.74 | Hill | 1.00 |
| 27bp | competitor | R306C | Hemi-methylated | 1 | 53.36 | 0.97 | Hill | 1.00 |
| 27bp | competitor | R306C | Hemi-methylated | 2 | 59.48 | 1.36 | Hill | 1.00 |
| 27bp | competitor | R306C | Hemi-methylated | 3 | 59.87 | 1.23 | Hill | 1.00 |
| 27bp | competitor | R306C | Methylated | 1 | 48.69 | 1.32 | Hill | 1.00 |
| 27bp | competitor | R306C | Methylated | 2 | 48.59 | 1.45 | Hill | 1.00 |
| 27bp | competitor | R306C | Methylated | 3 | 46.56 | 0.76 | Hill | 1.00 |
| 27bp | competitor | R306C | Unmethylated | 1 | 45.63 | 3.98 | Hill | 0.99 |
| 27bp | competitor | R306C | Unmethylated | 2 | 57.47 | 1.14 | Hill | 1.00 |
| 27bp | competitor | R306C | Unmethylated | 3 | 51.53 | 1.50 | Hill | 1.00 |
| 27bp | competitor | T158M | Hemi-methylated | 1 | 62.57 | 2.03 | Hill | 1.00 |
| 27bp | competitor | T158M | Hemi-methylated | 2 | 65.65 | 1.15 | Hill | 1.00 |
| 27bp | competitor | T158M | Hemi-methylated | 3 | 71.97 | 1.30 | Hill | 1.00 |
| 27bp | competitor | T158M | Methylated | 1 | 56.45 | 1.89 | Hill | 1.00 |
| 27bp | competitor | T158M | Methylated | 2 | 55.31 | 2.46 | Hill | 0.99 |
| 27bp | competitor | T158M | Methylated | 3 | 56.81 | 1.97 | Hill | 1.00 |
| 27bp | competitor | T158M | Unmethylated | 1 | 59.82 | 1.12 | Hill | 1.00 |
| 27bp | competitor | T158M | Unmethylated | 2 | 66.05 | 1.15 | Hill | 1.00 |
| 27bp | competitor | T158M | Unmethylated | 3 | 71.81 | 1.72 | Hill | 1.00 |
| 27bp | competitor | WT | Hemi-methylated | 1 | 74.11 | 1.67 | Hill | 1.00 |
| 27bp | competitor | WT | Hemi-methylated | 2 | 81.30 | 2.08 | Hill | 1.00 |
| 27bp | competitor | WT | Hemi-methylated | 3 | 69.39 | 1.72 | Hill | 1.00 |
| 27bp | competitor | WT | Methylated | 1 | 72.56 | 3.16 | Hill | 0.99 |
| 27bp | competitor | WT | Methylated | 2 | 75.50 | 3.32 | Hill | 0.99 |
| 27bp | competitor | WT | Methylated | 3 | 68.53 | 1.72 | Hill | 1.00 |
| 27bp | competitor | WT | Unmethylated | 1 | 58.32 | 1.43 | Hill | 1.00 |
| 27bp | competitor | WT | Unmethylated | 2 | 63.49 | 2.28 | Hill | 0.99 |
| 27bp | competitor | WT | Unmethylated | 3 | 58.04 | 1.68 | Hill | 1.00 |
| 27bp | none | R133C | Hemi-methylated | 1 | 1.24 | 0.05 | Hill | 1.00 |
| 27bp | none | R133C | Hemi-methylated | 2 | 1.29 | 0.03 | Hill | 1.00 |
| 27bp | none | R133C | Hemi-methylated | 3 | 2.11 | 0.19 | Hill | 0.99 |
| 27bp | none | R133C | Methylated | 1 | 2.23 | 0.10 | Hill | 1.00 |
| 27bp | none | R133C | Methylated | 2 | 2.28 | 0.54 | Morrison | 0.97 |
| 27bp | none | R133C | Methylated | 3 | 3.57 | 0.17 | Hill | 1.00 |
| 27bp | none | R133C | Unmethylated | 1 | 1.56 | 0.09 | Hill | 0.99 |
| 27bp | none | R133C | Unmethylated | 2 | 1.72 | 0.10 | Hill | 1.00 |
| 27bp | none | R133C | Unmethylated | 3 | 3.52 | 0.31 | Hill | 0.99 |
| 27bp | none | R168X | Hemi-methylated | 1 | 35.83 | 6.17 | Morrison | 0.98 |
| 27bp | none | R168X | Hemi-methylated | 2 | 39.28 | 9.46 | Morrison | 0.96 |
| 27bp | none | R168X | Hemi-methylated | 3 | 17.95 | 3.39 | Morrison | 0.98 |
| 27bp | none | R168X | Methylated | 1 | 151.54 | 132.26 | Hill | 0.96 |

| | | | | | | | | |
|---|---|---|---|---|---|---|---|---|
| 27bp | none | R168X | Methylated | 2 | 166.24 | 157.54 | Hill | 0.96 |
| 27bp | none | R168X | Methylated | 3 | 122.85 | 86.27 | Hill | 0.98 |
| 27bp | none | R168X | Unmethylated | 1 | 83.77 | 15.09 | Hill | 1.00 |
| 27bp | none | R168X | Unmethylated | 2 | 159.16 | 56.19 | Hill | 0.99 |
| 27bp | none | R168X | Unmethylated | 3 | 82.02 | 31.13 | Hill | 0.98 |
| 27bp | none | R255X | Hemi-methylated | 1 | 86.19 | 10.16 | Hill | 0.99 |
| 27bp | none | R255X | Hemi-methylated | 2 | 69.41 | 2.46 | Hill | 1.00 |
| 27bp | none | R255X | Hemi-methylated | 3 | 83.23 | 11.60 | Morrison | 0.99 |
| 27bp | none | R255X | Methylated | 1 | 99.11 | 4.74 | Hill | 1.00 |
| 27bp | none | R255X | Methylated | 2 | 92.86 | 6.66 | Hill | 1.00 |
| 27bp | none | R255X | Methylated | 3 | 89.67 | 4.88 | Hill | 1.00 |
| 27bp | none | R255X | Unmethylated | 1 | 82.66 | 10.57 | Hill | 0.99 |
| 27bp | none | R255X | Unmethylated | 2 | 75.42 | 4.74 | Hill | 1.00 |
| 27bp | none | R255X | Unmethylated | 3 | 97.88 | 10.14 | Hill | 0.99 |
| 27bp | none | R270X | Hemi-methylated | 1 | 4.96 | 0.59 | Morrison | 0.99 |
| 27bp | none | R270X | Hemi-methylated | 2 | 4.07 | 0.45 | Morrison | 0.99 |
| 27bp | none | R270X | Hemi-methylated | 3 | 2.98 | 0.33 | Morrison | 0.99 |
| 27bp | none | R270X | Methylated | 1 | 9.22 | 1.12 | Morrison | 0.99 |
| 27bp | none | R270X | Methylated | 2 | 7.09 | 5.04 | Morrison | 0.64 |
| 27bp | none | R270X | Methylated | 3 | 6.46 | 0.88 | Morrison | 0.99 |
| 27bp | none | R270X | Unmethylated | 1 | 5.02 | 0.74 | Morrison | 0.99 |
| 27bp | none | R270X | Unmethylated | 2 | 6.16 | 0.92 | Morrison | 0.99 |
| 27bp | none | R270X | Unmethylated | 3 | 7.01 | 1.30 | Morrison | 0.98 |
| 27bp | none | R306C | Hemi-methylated | 1 | 1.96 | 0.10 | Hill | 1.00 |
| 27bp | none | R306C | Hemi-methylated | 2 | 0.10 | 0.04 | Morrison | 0.99 |
| 27bp | none | R306C | Hemi-methylated | 3 | 1.65 | 0.10 | Hill | 0.99 |
| 27bp | none | R306C | Methylated | 1 | 2.28 | 0.10 | Hill | 1.00 |
| 27bp | none | R306C | Methylated | 2 | 1.37 | 0.09 | Hill | 0.99 |
| 27bp | none | R306C | Methylated | 3 | 1.92 | 0.05 | Hill | 1.00 |
| 27bp | none | R306C | Unmethylated | 1 | 2.32 | 0.14 | Hill | 0.99 |
| 27bp | none | R306C | Unmethylated | 2 | 0.61 | 0.08 | Morrison | 1.00 |
| 27bp | none | R306C | Unmethylated | 3 | 2.02 | 0.06 | Hill | 1.00 |
| 27bp | none | T158M | Hemi-methylated | 1 | 1.14 | 0.07 | Hill | 0.99 |
| 27bp | none | T158M | Hemi-methylated | 2 | 0.10 | 0.04 | Morrison | 0.99 |
| 27bp | none | T158M | Hemi-methylated | 3 | 0.44 | 0.08 | Morrison | 0.99 |
| 27bp | none | T158M | Methylated | 1 | 0.98 | 0.22 | Morrison | 0.98 |
| 27bp | none | T158M | Methylated | 2 | 1.55 | 0.08 | Hill | 1.00 |
| 27bp | none | T158M | Methylated | 3 | 1.11 | 0.18 | Morrison | 0.99 |
| 27bp | none | T158M | Unmethylated | 1 | 0.84 | 0.22 | Morrison | 0.98 |
| 27bp | none | T158M | Unmethylated | 2 | 1.96 | 0.09 | Hill | 1.00 |
| 27bp | none | T158M | Unmethylated | 3 | 2.66 | 0.19 | Hill | 0.99 |
| 27bp | none | WT | Hemi-methylated | 1 | 0.65 | 0.04 | Hill | 0.99 |
| 27bp | none | WT | Hemi-methylated | 2 | 0.82 | 0.08 | Hill | 0.98 |
| 27bp | none | WT | Hemi-methylated | 3 | 0.71 | 0.04 | Hill | 0.99 |
| 27bp | none | WT | Methylated | 1 | 0.82 | 0.05 | Hill | 0.99 |
| 27bp | none | WT | Methylated | 2 | 0.74 | 0.04 | Hill | 0.99 |
| 27bp | none | WT | Methylated | 3 | 0.73 | 0.04 | Hill | 0.99 |
| 27bp | none | WT | Unmethylated | 1 | 0.97 | 0.09 | Hill | 0.99 |
| 27bp | none | WT | Unmethylated | 2 | 0.90 | 0.04 | Hill | 1.00 |
| 27bp | none | WT | Unmethylated | 3 | 0.30 | 0.03 | Morrison | 1.00 |

**Supplementary Table S4. Fold change between different variants of MeCP2 and WT MeCP2 in FP assay**

| length | protein | DNA | FC1 (Kd(+comp)/Kd(no-comp)) | FC1_SEM | n | p_value | significance | FC2 (Kd(+comp, variant)/Kd(+comp, WT)) | FC2_SEM | n | p_value | significance |
|---|---|---|---|---|---|---|---|---|---|---|---|---|
| 27bp | WT | Hemi-methylated | 103.37 | 5.03 | 3 | 0.0001 | *** | 1.00 | 0.00 | 3 | | |
| 27bp | R133C | Hemi-methylated | 49.69 | 7.62 | 3 | 0.0019 | ** | 0.97 | 0.05 | 3 | 0.589 | ns |
| 27bp | R306C | Hemi-methylated | 31.78 | 4.55 | 2 | 0.0266 | * | 0.77 | 0.05 | 3 | 0.0124 | * |
| 27bp | T158M | Hemi-methylated | 109.35 | 54.35 | 2 | 0.0759 | ns | 0.90 | 0.07 | 3 | 0.1348 | ns |
| 27bp | R255X | Hemi-methylated | 1.43 | 0.24 | 3 | 0.2172 | ns | 1.50 | 0.19 | 3 | 0.1276 | ns |
| 27bp | R270X | Hemi-methylated | 38.45 | 4.18 | 3 | 0.0009 | *** | 1.99 | 0.06 | 3 | 0.0004 | *** |
| 27bp | R168X | Hemi-methylated | 7.86 | 1.87 | 3 | 0.0118 | * | 2.93 | 0.04 | 3 | 0.0001 | *** |
| 15bp | WT | Hemi-methylated | 49.89 | 2.43 | 3 | 0.0002 | *** | 1.00 | 0.00 | 3 | | |
| 15bp | R133C | Hemi-methylated | 38.27 | 4.70 | 3 | 0.0012 | ** | 1.58 | 0.08 | 3 | 0.004 | ** |
| 15bp | R306C | Hemi-methylated | 115.15 | 28.59 | 3 | 0.0031 | ** | 1.37 | 0.13 | 3 | 0.1361 | ns |
| 15bp | T158M | Hemi-methylated | 36.76 | 6.91 | 3 | 0.0029 | ** | 1.31 | 0.11 | 3 | 0.0598 | ns |
| 15bp | R255X | Hemi-methylated | 2.01 | 0.13 | 3 | 0.0087 | ** | 1.99 | 0.11 | 3 | 0.0008 | *** |
| 15bp | R270X | Hemi-methylated | 49.46 | 8.84 | 3 | 0.0019 | ** | 1.97 | 0.10 | 3 | 0.0038 | ** |
| 15bp | R168X | Hemi-methylated | 2.49 | 0.30 | 3 | 0.0167 | * | 4.25 | 0.19 | 3 | 0 | *** |
| 27bp | WT | Methylated | 95.05 | 3.85 | 3 | 0.0001 | *** | 1.00 | 0.00 | 3 | | |
| 27bp | R133C | Methylated | 25.39 | 3.56 | 3 | 0.0022 | ** | 0.91 | 0.04 | 3 | 0.0496 | * |
| 27bp | R306C | Methylated | 27.03 | 4.29 | 3 | 0.0022 | ** | 0.66 | 0.01 | 3 | 0.001 | *** |
| 27bp | T158M | Methylated | 48.11 | 6.48 | 3 | 0.0014 | ** | 0.78 | 0.03 | 3 | 0.0083 | ** |
| 27bp | R255X | Methylated | 1.23 | 0.17 | 3 | 0.3413 | ns | 1.60 | 0.23 | 3 | 0.1212 | ns |
| 27bp | R270X | Methylated | 21.75 | 2.48 | 3 | 0.0016 | ** | 2.23 | 0.05 | 3 | 0.0001 | *** |
| 27bp | R168X | Methylated | 2.14 | 0.29 | 3 | 0.0357 | * | 4.26 | 0.38 | 3 | 0.0007 | *** |
| 15bp | WT | Methylated | 36.76 | 7.45 | 3 | 0.0042 | ** | 1.00 | 0.00 | 3 | | |
| 15bp | R133C | Methylated | 37.85 | 4.59 | 3 | 0.0011 | ** | 1.65 | 0.17 | 3 | 0.0476 | * |
| 15bp | R306C | Methylated | 32.83 | 7.34 | 2 | 0.0417 | * | 1.19 | 0.15 | 3 | 0.3303 | ns |
| 15bp | T158M | Methylated | 27.36 | 0.72 | 3 | 0.0001 | *** | 1.28 | 0.09 | 3 | 0.0841 | ns |
| 15bp | R255X | Methylated | 1.88 | 0.03 | 3 | 0.0005 | *** | 2.28 | 0.10 | 3 | 0.0021 | ** |
| 15bp | R270X | Methylated | 64.33 | 4.99 | 3 | 0.0004 | *** | 1.80 | 0.07 | 3 | 0.0032 | ** |
| 15bp | R168X | Methylated | 6.78 | 1.39 | 3 | 0.0103 | * | 2.11 | 0.11 | 3 | 0.0045 | ** |
| 27bp | WT | Unmethylated | 108.14 | 43.00 | 3 | 0.0065 | ** | 1.00 | 0.00 | 3 | | |
| 27bp | R133C | Unmethylated | 28.88 | 5.24 | 3 | 0.0038 | ** | 0.99 | 0.07 | 3 | 0.7812 | ns |
| 27bp | R306C | Unmethylated | 22.6 | 2.95 | 2 | 0.0268 | * | 0.86 | 0.04 | 3 | 0.132 | ns |
| 27bp | T158M | Unmethylated | 43.89 | 13.69 | 3 | 0.0062 | ** | 1.10 | 0.07 | 3 | 0.2187 | ns |
| 27bp | R255X | Unmethylated | 1.44 | 0.29 | 3 | 0.2981 | ns | 2.00 | 0.33 | 3 | 0.0531 | ns |
| 27bp | R270X | Unmethylated | 27.49 | 2.88 | 3 | 0.0009 | *** | 2.74 | 0.17 | 3 | 0 | *** |
| 27bp | R168X | Unmethylated | 2.9 | 0.55 | 3 | 0.042 | * | 4.79 | 0.16 | 3 | 0 | *** |
| 15bp | WT | Unmethylated | 37.68 | 3.75 | 3 | 0.0008 | *** | 1.00 | 0.00 | 3 | | |
| 15bp | R133C | Unmethylated | 32.63 | 3.14 | 3 | 0.0008 | *** | 1.59 | 0.16 | 3 | 0.0293 | * |
| 15bp | R306C | Unmethylated | 72.31 | 5.71 | 3 | 0.0004 | *** | 1.55 | 0.17 | 3 | 0.0802 | ns |
| 15bp | T158M | Unmethylated | 42.08 | 10.43 | 3 | 0.0059 | ** | 1.40 | 0.14 | 3 | 0.0775 | ns |
| 15bp | R255X | Unmethylated | 1.92 | 0.26 | 3 | 0.0475 | * | 2.33 | 0.13 | 3 | 0.0046 | ** |
| 15bp | R270X | Unmethylated | 16.15 | 2.25 | 3 | 0.0029 | ** | 2.13 | 0.03 | 3 | 0 | *** |
| 15bp | R168X | Unmethylated | 1.02 | 0.10 | 3 | 0.9148 | ns | 5.87 | 0.20 | 3 | 0.0002 | *** |
